# A deep learning framework for understanding cochlear implants

**DOI:** 10.1101/2025.07.16.665227

**Authors:** Annesya Banerjee, Mark R. Saddler, Julie G. Arenberg, Josh H. McDermott

**Affiliations:** Program in Speech and Hearing Biosciences and Technology, Harvard University; Department of Brain and Cognitive Sciences, Massachusetts Institute of Technology; K. Lisa Yang Brain-Body Center, Massachusetts Institute of Technology; McGovern Institute for Brain Research, Massachusetts Institute of Technology; Department of Health Technology, Technical University of Denmark; Mass Eye and Ear, Harvard Medical School, Department of Otolaryngology Head and Neck Surgery; Center for Brains, Minds and Machines, Massachusetts Institute of Technology

## Abstract

Sensory prostheses replace dysfunctional sensory organs with electrical stimulation but currently fail to restore normal perception. Outcomes may be limited by stimulation strategies, neural degeneration, or suboptimal decoding by the brain. We propose a deep learning framework to evaluate these issues by estimating best-case outcomes with task-optimized decoders operating on simulated prosthetic input. We applied the framework to cochlear implants – the standard treatment for deafness – by training artificial neural networks to recognize and localize sounds using simulated auditory nerve input. The resulting models exhibited speech recognition and sound localization that was worse than that of normal hearing listeners, and on par with the best human cochlear implant users, with similar results across the three main stimulation strategies in current use. Speech recognition depended heavily on the extent of decoder optimization for implant input, with lesser influence from other factors. The results identify performance limits of current devices and demonstrate a model-guided approach for understanding the limitations and potential of sensory prostheses.

## Introduction

Over 70 million people worldwide are deaf, and over 39 million people are blind^1^. In most cases the root cause lies in sensory receptor dysfunction. Sensory prostheses aim to restore perception in the deficient sense, by supplanting faulty receptors with machine sensors and conveying a stimulus directly to peripheral neurons via electrical stimulation^2^. For instance, in many deaf individuals the hair cells that normally transduce the mechanical vibrations of sound do not function, but the auditory nerve, which normally carries signals from the hair cells to the brain, is at least partially intact. The nerve can thus be electrically stimulated, causing the perception of sound^3^, i.e., “electrical hearing” (Fig. 1A).

**Fig. 1.**
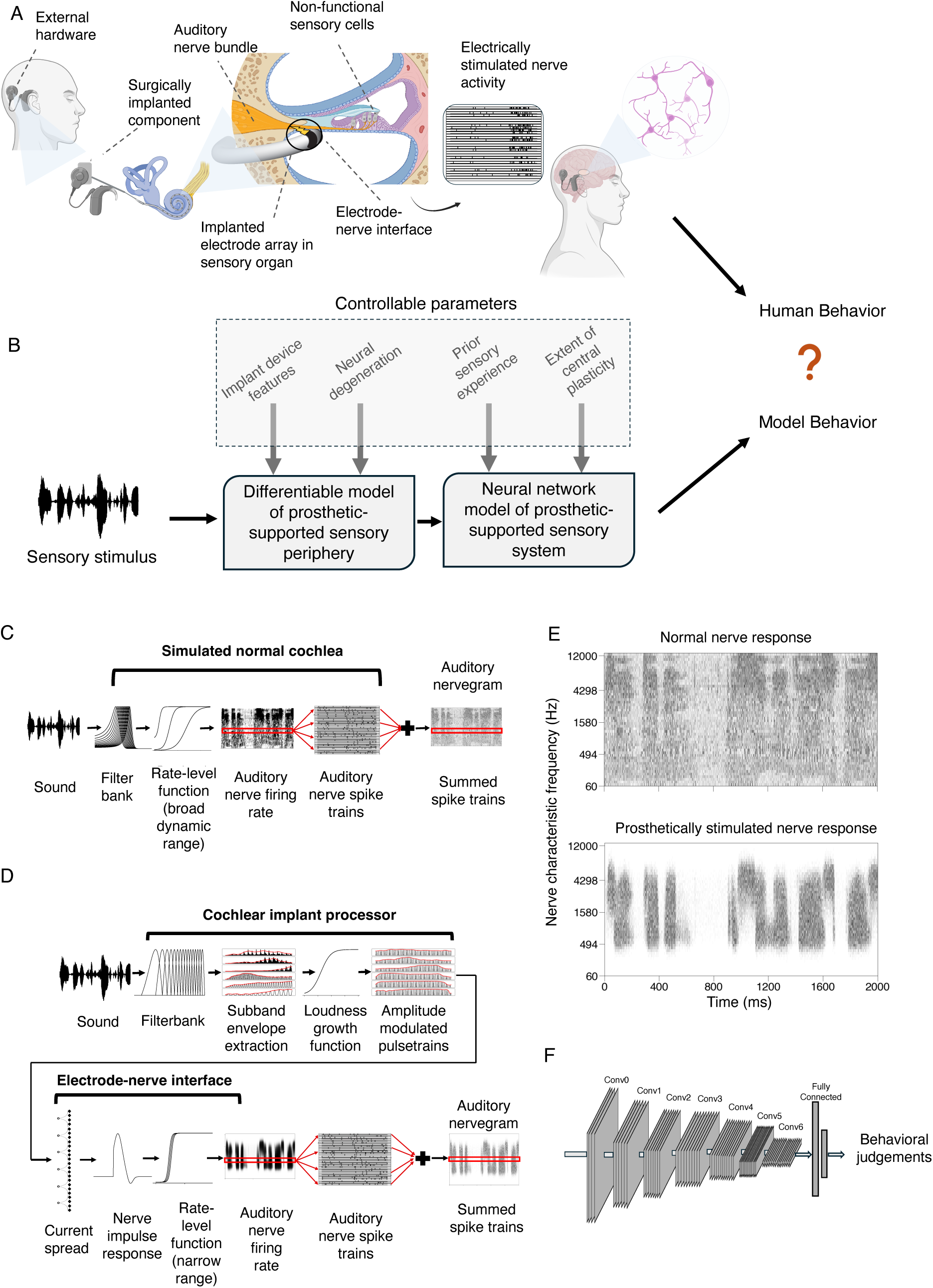
Simulation of peripheral stimulation and central decoding. **A.** Sensory transduction in prosthetic-aided sensory systems. Center panel shows cross section of cochlea. Figure created in http://biorender.com/. **B.** Modeling framework for studying cochlear-implant-mediated hearing. **C.** Simulation of normal hearing auditory nerve response. The responses of multiple nerve fibers are sampled from the firing rate at a characteristic frequency, and are then summed. Here and in D, the red boxes and spike trains show this process for an example characteristic frequency. **D.** Simulation of cochlear-implant-stimulated auditory nerve response, comprising stages of signal processing within a cochlear implant processor, and the electrode-nerve interface that produces nerve responses from electrode signals. The rate-level function for electrical stimulation has a narrower dynamic range compared to the acoustically driven response of the normal cochlea (because electrical stimulation bypasses the amplitude compression of the cochlea). **E.** Example simulated nerve responses for a normal cochlea (top) and a cochlear implant (bottom) in response to an excerpt of speech. **F.** Schematic of deep convolutional neural network model of central auditory decoding.

Cochlear implants are by far the most successful and widely used sensory prosthesis, with over one million^4^ devices in use worldwide. By comparison, fewer than 1000 retinal implants^5^ and fewer than 2000 auditory brainstem implants^6^ have been implanted globally. Cochlear implants are a remarkable feat of science and engineering, in the best cases enabling previously deaf individuals to engage in conversations over a telephone^3^ (for which they are entirely reliant on auditory information). On the other hand, outcomes are highly variable, and even in the best cases current implants are far from restoring “normal” hearing^7,8^.

The factors limiting outcomes with sensory prostheses remain uncertain. Prostheses rely on the neural pathways that carry sensory signals to the brain, but these pathways often degenerate when they are deprived of input for a prolonged period of time^9,10^, potentially limiting perceptual abilities. There is also considerable evidence that people benefit from learning to extract information from prosthetically stimulated nerve activity^11–14^. For instance, in cochlear implant recipients, speech recognition scores typically improve over months to years following implantation^15^, suggestive of brain plasticity that enables better decoding of the implant stimulation. It is also likely that the strategies for converting external sensory stimuli into electrical signals are suboptimal^4^. For instance, contemporary cochlear implants discard some of the cues to a sound’s location^16^, which might be expected to limit how well sounds can be localized. Moreover, the resolution with which a sound’s frequency content can be encoded is thought to be limited by the spatial spread of excitation along the cochlea^7^, which depends in part on the strategy for encoding sound in electrode signals.

Each of these factors plausibly limits on how well a person can perceive the world through a prosthesis, but it is difficult to know their importance. Moreover, because the best outcomes from a particular device strategy require extended periods of use by a human, and because outcomes are highly variable across individuals^15^, it is intractable to empirically test large sets of alternative strategies in humans. It thus remains unclear whether substantially better outcomes might be possible with alternative strategies. These issues apply broadly to all sensory prostheses.

In this paper, we propose to understand sensory prostheses using models of auditory perception built using deep learning^17–20^. The core idea is to train an artificial neural network to perform real-world auditory tasks using input from a simulated prosthesis (Fig. 1B). The performance of the resulting model should approach an upper bound on how well a human with the same prosthesis should be able to perform. The model could then be used to explore alternative stimulation strategies at scale, to assess whether alternatives not in current use might give substantially better outcomes. Such a model could in principle also be used to test the potential effect of internal factors that might account for suboptimal performance by human prosthesis users, by measuring their effect on performance in a working system in which individual factors can be manipulated. The primary purpose of this paper is to illustrate the framework by using it to evaluate best-case scenario behavioral outcomes for cochlear implant strategies that are currently in widespread use. We sought to answer four main scientific questions. The first question was how well contemporary cochlear implants should in principle enable performance on two important auditory tasks (word recognition and sound localization). The second question was whether any performance limits could be clearly linked to specific device-related factors. The third question was whether high-performing human cochlear implant users currently come close to these best-case scenario performance levels. The fourth question was whether gaps between individual human users and best-case scenario performance levels could be accounted for in a model with the various internal factors proposed to limit cochlear implant outcomes. There is no guarantee that the effect of these factors in deep learning models will replicate those in the brain, but model outcomes nonetheless provide an example of what might be expected to happen in a working sensory system.

We built a simulation of cochlear implant stimulation of the auditory nerve, leveraging the decades of research that have characterized these effects in the auditory system^21–28^. We then trained neural networks to decode speech and sound location from the simulated nerve responses. Models optimized for input from contemporary cochlear implants recognized speech and localized sounds better than typical human cochlear implant users, but in both cases performed substantially worse than models optimized for normal hearing input. Results were similar for the three main stimulation strategies in widespread use, revealing limitations in what can be decoded from current devices. The approach provides a general framework for studying and improving sensory prostheses.

## Results

### Overall approach

The models we consider begin by simulating the representations of sound in the auditory nerve in both normal and electrical hearing with established stages of signal processing (Fig. 1C&D). It is apparent from glancing at example simulated nerve responses to speech (Fig. 1E) that the input to the brain from a cochlear implant differs drastically from that obtained from the normal ear. These differences are a consequence of how electrical stimulation induces action potentials in the nerve, producing reduced frequency resolution and dynamic range, among other differences^22,29^. Such differences lend plausibility to the idea that good performance using cochlear implant input might require a different “decoder” than that used for normal hearing, as might result from plasticity following the implant. Moreover, it is not obvious what levels of performance should be possible given optimal decoding.

We use machine learning to derive optimized decoders for real-world perceptual problems using simulated nerve representations as input (Fig. 1F). The solutions arrived at by machine learning have been repeatedly shown to yield human-like behavior in many settings when optimized given input from a simulated normal ear^17–20^, consistent with the idea that both humans and machine learning models end up relatively well optimized for natural tasks performed on input from healthy ears. Such results give credence to the idea that similar models optimized for cochlear implant input should approach the best performance that is possible with such input. However, it was not clear what performance levels to expect. One previous attempt to model cochlear implant-mediated speech recognition with neural networks was plausibly hindered by training datasets that are small by modern standards^30^, and that likely limit the model predictions in diverse realistic conditions^31^. Another recent such attempt may have been limited by unrealistic simulations of cochlear implant input^32^. It was thus unclear what would happen in a model with realistic input optimized in realistic conditions. We evaluated candidate central decoders by comparing them to a model of normal hearing that replicates the absolute performance of human listeners, and to the performance of human cochlear implant users.

### Normal hearing auditory nerve model stage

Models of normal hearing began with a fixed stage of simulated auditory nerve responses developed in prior work^20^. It was intended to be as realistic as possible subject to the practical constraints of being efficient to run in a deep learning setting. We simulated 32,000 spiking nerve fibers with each of the three qualitative fiber types found in the mammalian ear^33^.

### Cochlear implant and electrode-to-nerve interface model stages

Models of hearing through a cochlear implant began with a fixed stage of simulated electrically stimulated nerve responses. This stage can be conceived as containing two sub-stages: one consisting of the cochlear implant device, which generates electrode signals from sound input, and one consisting of the electrode-to-nerve interface, which generates auditory nerve responses from the electrode signals. These stages were again intended to be as realistic as possible subject to constraints of computational efficiency.

The simulated cochlear implant processor was based on the signal processing operations in current commercially available devices, which include audio subband generation using a filter bank (one filter per electrode), extraction of subband envelopes, amplitude compression, and amplitude modulation of pulse trains^3,7,8^. Amplitude compression serves to map sound energy onto the relatively small (10-20 dB) dynamic range of electrical stimulation (outside of which the auditory nerve either does not respond, or is saturated)^34–37^. We set the parameters of this mapping by maximizing the variance in the nerve response across stimuli, and then confirming that these parameters were sufficient to account for psychophysical abilities of human cochlear implant users (see below).

The resulting compressed envelopes were used to amplitude-modulate a train of pulses. Our baseline model used the continuous interleaved sampling (CIS)-based sound coding strategy^38^, in which pulses across electrodes are interleaved in time to minimize electrical interference. The sound coding strategies in widespread use in current commercial cochlear implants are all based on CIS. We also implemented models using the strategies used in the three main devices available today: Advanced Combination Encoder (ACE; used in Cochlear Ltd. devices)^39^, Fine Structure Processing (FSP; used in MED-EL devices)^40^, and High Resolution with Fidelity 120 (HiRES120; used in Advanced Bionics devices)^41^. Each of these three strategies is a variation on CIS. The specific versions of FSP and HiRES120 that we implemented are slightly older versions of the strategies available in new devices from MED-EL and Advanced Bionics, but the versions we used are a) representative of those in most devices currently in use, and b) only modestly different from the latest versions^42^.

The electrode-to-nerve interface captured the established behavior of auditory nerve responses under electrical stimulation^21–28^. Electrical pulses at an electrode create an electric field that spreads across space and excites neurons over some portion of the cochlea. We simulated this spread with exponential decay^43–46^, setting the decay rate to simulate the spread estimated to occur for the monopolar stimulation common to contemporary implants (we test the effect of decreasing this spread in subsequent experiments). We then summed the electrical fields from each electrode and simulated the neural response at each place along the cochlea by convolving the time-varying electrical field with an impulse response matched to the timescale of the nerve response^47^ (see Methods). The result was converted to firing rates using a rate-level function with a narrow dynamic range. These components are similar to those in previous models of electrically stimulated nerve responses^31,45,48–54^, but omit some details (e.g. auditory nerve refractoriness) due to computational constraints. See Methods for complete description and Discussion for the possible ramifications of these omissions. We sampled spikes from the resulting firing rates as in the normal auditory nerve model stage (see Methods).

### Validating current settings with intensity discrimination thresholds

Most of the free parameters in the simulation of nerve responses could be set from known values or estimated from prior work (see Methods). The current settings were the lone exception. A cochlear implant maps a representation of audio (subband envelopes, in the case of the contemporary devices studied in this paper) onto electrical current. In a human cochlear implant user, the absolute current levels are set by an audiologist based on behavioral feedback. The audiologist can set the minimum current to something that is just barely audible, and the maximum level to that just below what is uncomfortably loud (maximum comfortable level; MCL). In practice, because the difference between these levels (i.e., the usable dynamic range) tends to be fairly consistent across human users (∼20 dB), an audiologist often determines only the maximum comfortable level based on patient feedback, and then sets the minimum current level to a fixed fraction of the maximum. For example, in Advanced Bionics devices, the threshold is set to be a tenth of the maximum comfortable level^55^. This process cannot be directly replicated in a model, because we do not know how loudness relates to the auditory nerve response. In most previous models of cochlear implants, the current parameters have been set based on an assumed neural correlate of the maximum comfortable level that has yet to be validated^45,56^.

We found empirically that the current settings could have a substantial effect on model behavior, so it seemed important to set them in a principled way. We opted to set current levels based on two constraints. First, we assumed that current level settings in humans enable different sounds to be distinguished from the nerve responses they produce – in other words, the current levels avoid responses that are pinned near zero or that are saturated. Second, we required that the current settings in the model be sufficient to reproduce intensity discrimination abilities of human cochlear implant users^57,58^, on the assumption that such abilities are largely determined by the characteristics of peripheral responses^59,60^.

We first measured the variance of nerve responses across a set of speech stimuli. We conducted a grid search over maximum current levels, constraining the minimum level to achieve a dynamic range of 20 dB to be consistent with the ranges that are used in humans. This analysis revealed a set of current settings that produced high variance across stimuli. We then measured intensity discrimination thresholds using the nerve representations from the set of candidate current settings, training a classifier on a set of training stimuli and then measuring psychometric functions on a set of held-out stimuli (Fig. 2A-D). As shown in Fig. 2E, some of the current settings led to model intensity discrimination on par with that observed in humans in previous work^57,61^, matching both absolute thresholds and the variation in threshold with intensity (within the dynamic range that resulted from the settings). We chose one of these settings to use in the default model (see Methods), but explore the effect of alternative settings later.

**Fig. 2.**
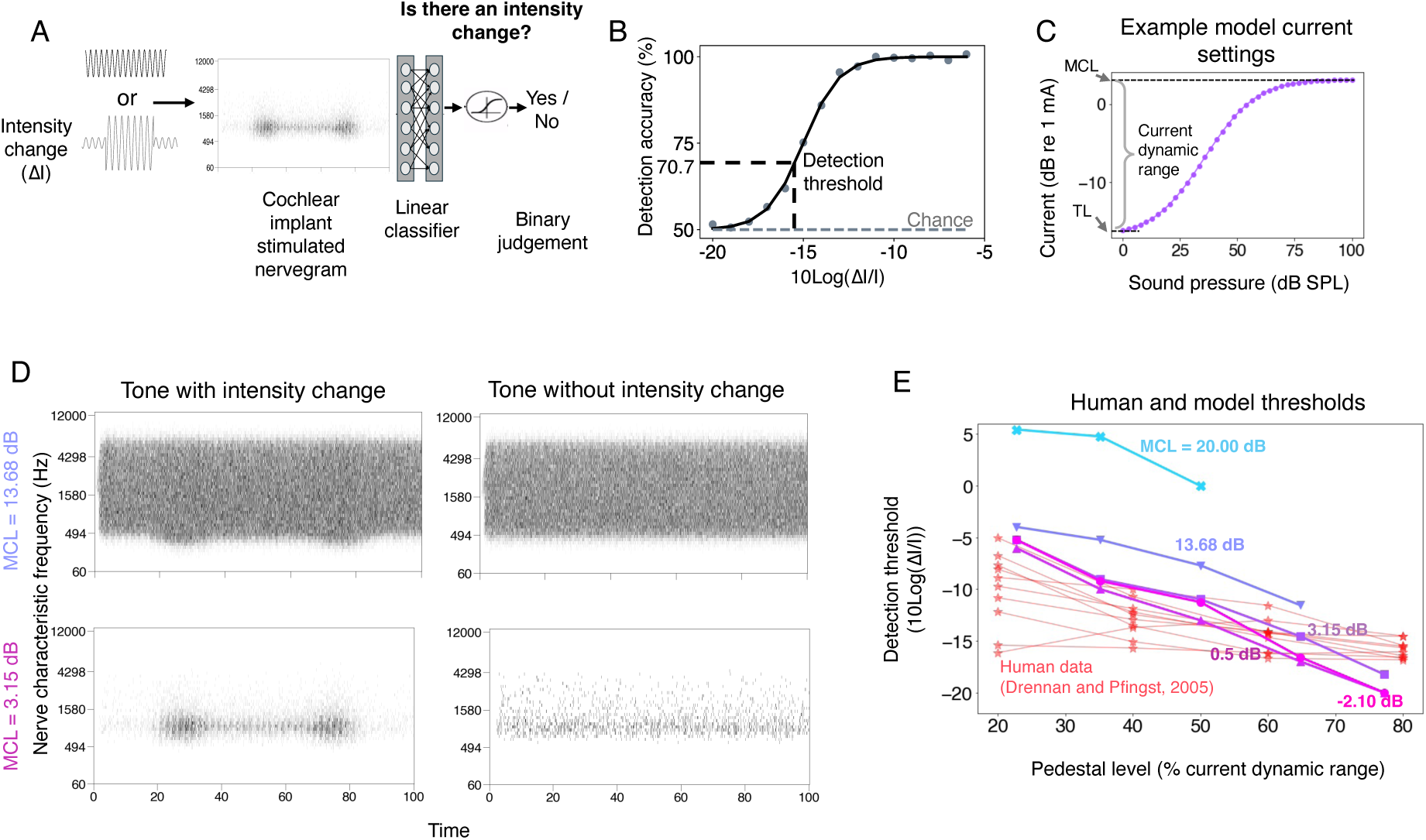
Models produced human-like intensity discrimination thresholds. **A.** Schematic of intensity discrimination experiment. Cochlear-implant-stimulated nerve responses corresponding to target tone stimuli (with/without intensity increment) were passed through a linear classifier. The classifier produced a binary judgment of whether the intensity changed or not. **B.** An example psychometric function from the model classifier. Detection thresholds were estimated from the psychometric function for an experimental condition. **C.** Example “loudness growth” functions (mapping acoustic sound pressure levels to current levels). The dynamic range of the current remained the same across different settings (20 dB); only the maximum current levels (MCL) changed, with the minimum (threshold level; TL) set to be 20 dB below the MCL. **D.** Example nerve responses of tone stimuli with and without an intensity change at two different current level settings: one for which the two stimuli are clearly distinguishable; another for which the two stimuli are less distinguishable. **E.** Comparison of intensity discrimination thresholds in the model for different current-level choices to those for human cochlear implant users (data from Drennan and Pfingst, 2005^61^). Human data are plotted for ten individual human participants. Error bars on model results plot standard error (not visible as they are smaller than the marker size). Error bars were not available for human data as they were scanned in from an earlier publication.

### Neural network decoding of speech and location from cochlear implant input

We optimized standard deep convolutional neural networks to decode auditory nerve input. When optimized for naturalistic auditory tasks, this type of model architecture has previously been shown to reproduce a wide range of human behaviors^17–20^, and to provide the best current predictions of human auditory cortical responses^17,62^ (though they also exhibit some significant discrepancies with humans^63^). The neural network consisted of a sequence of convolution, rectification, pooling, and normalization operations. The model weights were optimized using either normal or cochlear-implant-stimulated nerve input to perform one of two tasks: word recognition in noise^17,64^ or sound localization^19^. We used separate models for the two tasks for ease of implementation. We used at least three different model architectures for each task and device setting. These architectures were the result of architecture searches conducted in previous work that searched across a larger set of architectures to find those that worked well for the tasks^17,19^. The use of multiple architectures helped to ensure that the results were not idiosyncratic to a specific model architecture. The different architectures were treated akin to multiple human participants (i.e., we averaged results across architectures). In practice results were similar across architectures.

In the word recognition task, the model input was an excerpt of monaural audio, and the task was to identify the word spoken in the middle of the excerpt. In the sound localization task, the model input was an excerpt of binaural audio and the task was to identify the location (in azimuth and elevation) of the underlying sound source. In both cases the models were trained with supervised learning on a large set of labeled audio signals. The training data was intended to reflect the sorts of natural sounds that plausibly shaped the human auditory system over evolution and development. The models were optimized only for overall task performance, and were not fit to match human behavior in any way (apart from ensuring that the cochlear implant settings enabled human-like intensity resolution, as described above).

### Comparison to human cochlear implant users

To enable a direct comparison to humans, we tested humans and our models on the same experiments. For word recognition, we measured the performance of human cochlear implant users in our lab. Participants listened to speech excerpts presented in noise through their cochlear implant device connected to a computer via Bluetooth (see Methods). Word recognition accuracy was analyzed as a function of the signal-to-noise ratio (SNR). For sound localization, we tested models on a previously published experiment measuring the localization of noise bursts by listeners with normal hearing^65^ or cochlear implants^66^ (discussed below).

### Optimized decoders support speech recognition on par with the best human CI users

We first assessed the best-case scenario for perception through a prosthesis, by fully reoptimizing the neural network for prosthetic input, akin to a situation where the central auditory system is fully plastic, and where the plasticity serves to achieve the best possible task performance on (Fig. 3A). Fig. 3B shows word recognition performance in different noise levels for humans and models with normal hearing and cochlear implant input. As in prior work^17,20^, the model optimized for normal-hearing nerve input performed comparably to normal-hearing human listeners, even in noise. This model also reproduced patterns of speech intelligibility across different types of noise and speech distortions, specifically noise vocoding with different numbers of channels (Supplementary Fig. 1).

**Fig. 3.**
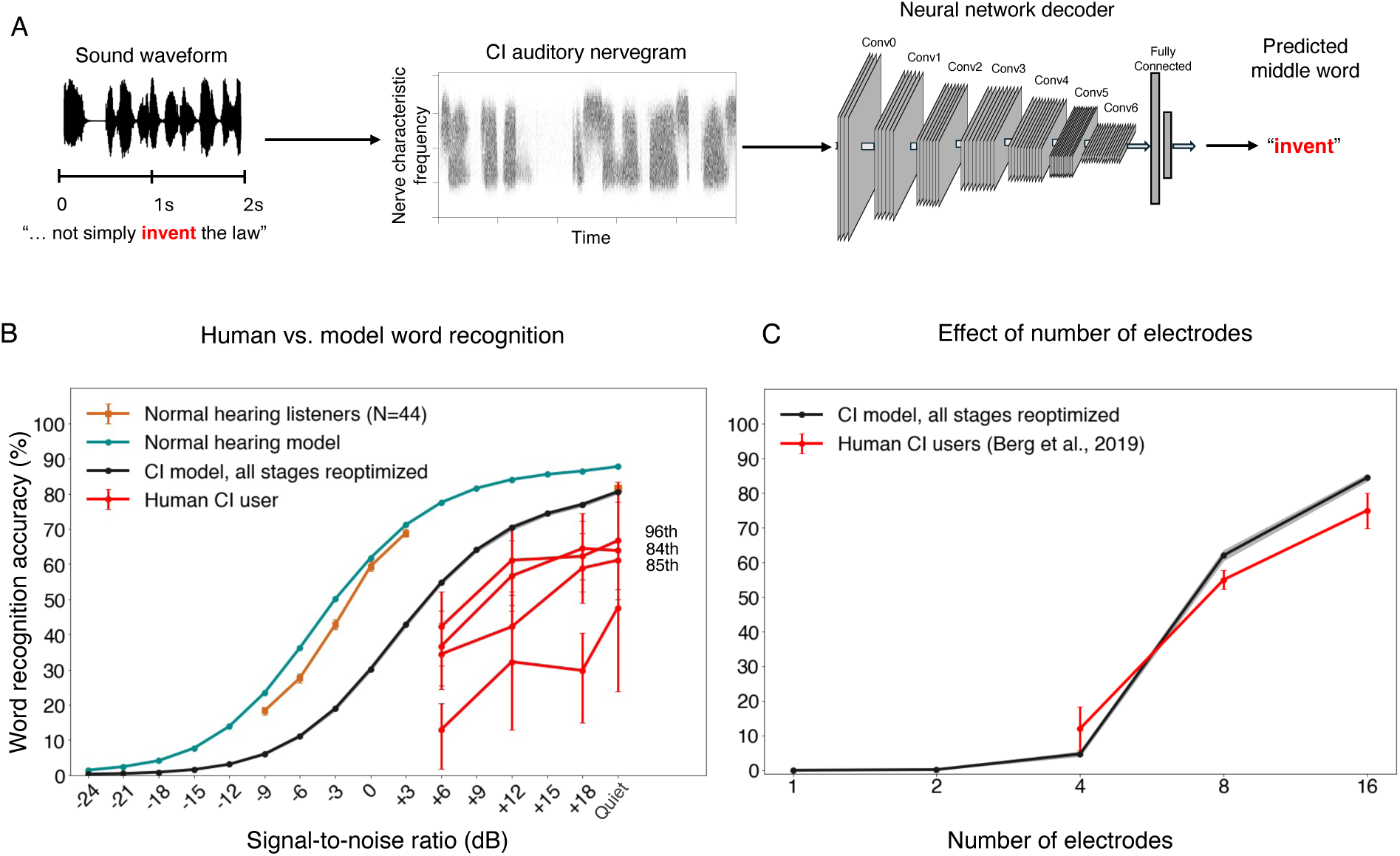
Optimized word recognition decoder outperforms human cochlear implant users. **A.** Task and model setup for word recognition. The task was to report the middle word in a 2s speech excerpt that was superimposed with some amount of additive noise. **B.** Word recognition accuracy as a function of the signal-to-noise ratio for normal hearing human participants (N=44)^20^, human cochlear implant users (N=4), and models optimized for either a simulation of the normal auditory nerve, or a cochlear-implant-stimulated auditory nerve. Error bars plot standard error across either model architectures or stimuli (for human participants). Percentile ranks for clinical word recognition scores in quiet are provided for the three cochlear implant users for whom this data was available. **C.** Effect of number of stimulated electrodes. Human data is replotted from a previous publication^67^. Model results are for the middle-word task used throughout this paper. Humans performed a different word recognition task, so absolute performance levels cannot be compared, but the qualitative effect of number of electrodes is similar for humans and the model.

The model fully reoptimized for cochlear implant input performed worse than the normal hearing models (the SNR yielding 50% correct was 7.92 dB lower), but far above chance even in moderate noise levels. This result indicates that the information provided by the cochlear implant can support relatively good recognition of speech in both quiet and noise provided it is appropriately decoded. There was substantial variability across human cochlear implant users, as expected^15^, and their word recognition performance remained far from that of normal-hearing listeners (Fig 3B), consistent with many prior such experiments. However, the best participants approached the performance level of the model (these participants were also high-performing as gauged by standard clinical word recognition scores; percentile ranks are shown in Fig. 3B).

To further probe whether the model captured the factors that constrain speech recognition in human cochlear implant users, we simulated an experiment in which stimulation was restricted to subsets of electrodes^67^ (see Methods). Model performance was poor for one or two electrodes but then improved as the number of stimulated electrodes increased, mirroring the qualitative trend found in human cochlear implant users (Fig. 3C).

### Optimized decoders also support sound localization on par with the best human CI users

We next conducted analogous experiments on models of sound localization. Cochlear implant users are known to perform poorly in sound localization tasks. Bilaterally implanted users show some benefits over unilateral users but are still not sufficient to achieve normal hearing listener-like performance^16,68^. There is reason to think that the strategies in current cochlear implants discard some of the most important cues to sound localization, in particular interaural time differences from stimulus fine structure. Nonetheless, it remains unclear what abilities should be expected with an optimized decoder.

We optimized models to perform localization of natural sounds in diffuse background noise (Fig. 4A). The models were trained using binaural nerve responses from spatialized audio. Sounds were spatialized using a room simulator and were filtered using head-related transfer functions to incorporate natural location cues. Models were trained to predict the azimuth and elevation location of the target sounds^19,20^.

**Fig. 4.**
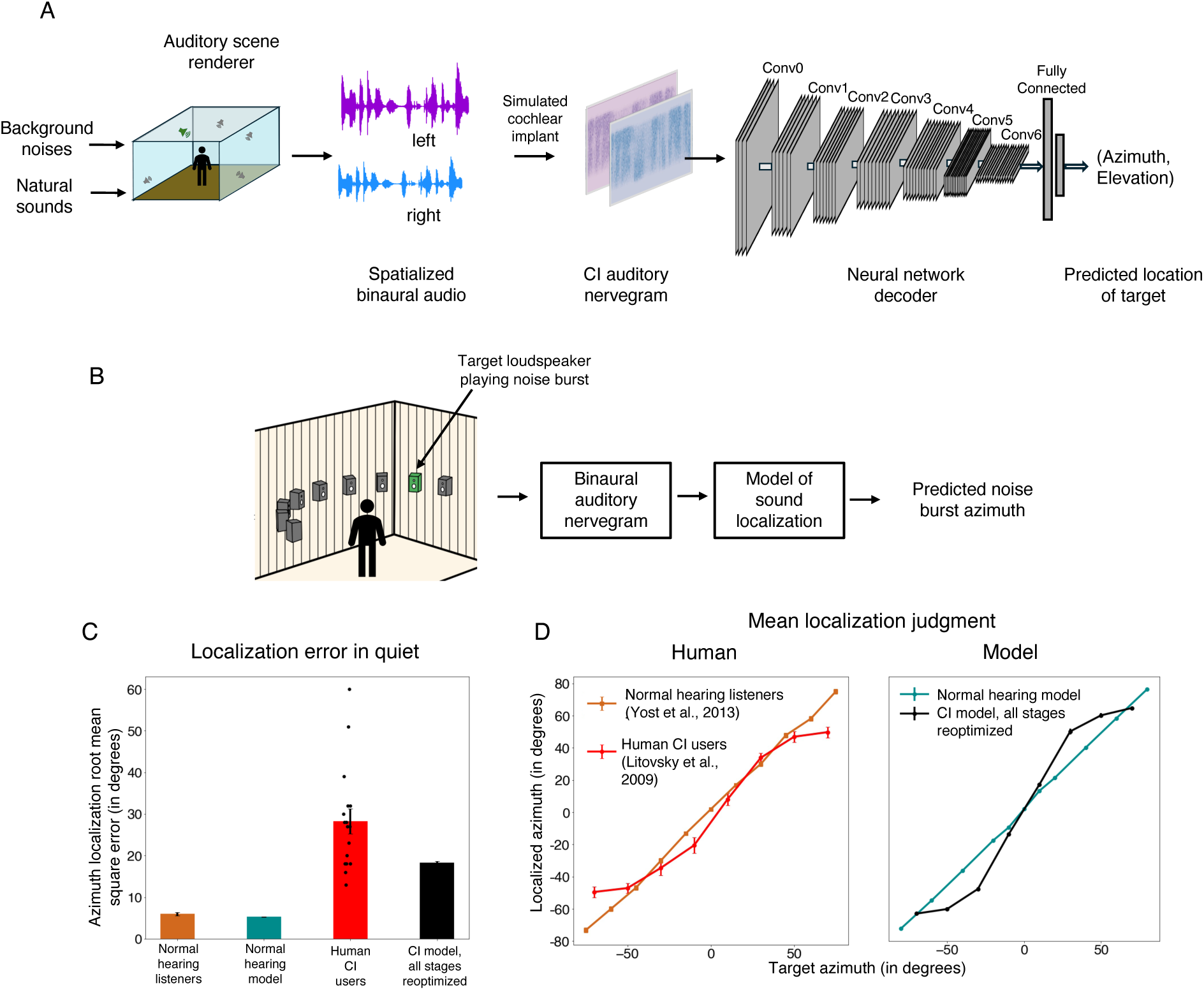
Model sound localization performance remains poor compared to normal hearing conditions. **A.** Training setup for sound localization models. Training data were generated using a room simulator that rendered the binaural audio signal that should be produced by a sound source to be localized (shown in green), when presented in a room, accompanied by noise sources (shown in gray). **B**. Localization experiment schematic. Neural network optimized to perform sound localization using binaural auditory nerve representation of virtually rendered auditory scenes. Left and right ear nerve representations were available as two distinct channels to the first stage of the neural network. **C.** Azimuthal localization error (calculated as root-mean-squared error) for human and model localization judgments of noise burst stimuli. Here and in panel D, error bars plot standard error across model architectures or human participants. Black dots plot results for individual human cochlear implant users. Human data are replotted from a previous publication^65,66^. **D**. Localized azimuth angle as a function of the target azimuth angle for humans and models.

Model localization performance was tested on an experiment measuring the localization of noise bursts in the azimuthal plane, that had previously been conducted with cochlear implant users^66^ (Fig. 4B). Performance was quantified using mean absolute localization error in degrees.

As in prior work^19^, and as seen for word recognition, the normal-hearing model of sound localization performed comparably to normal-hearing humans. The fully reoptimized cochlear implant model exhibited substantially worse performance than the normal hearing model (Fig. 4C), with roughly three times worse accuracy. As was the case for word recognition, the best-performing human cochlear implant users performed similarly to the model, with many participants who were substantially worse. The model also exhibited similar biases to those seen in humans (Fig. 4D), with distinct peripheral locations tending to be mapped onto similar perceived locations. These results confirm the idea that sound localization performance is likely limited by cochlear implant sound coding strategies that leave insufficient information in the auditory nerve representation.

### Similar outcomes across leading sound coding algorithms

Most cochlear implants in current use are made by one of three manufacturers (Fig. 5A), each of which uses a different sound coding algorithm (variants of continuous interleaved sampling): ACE (Cochlear), FSP (MED-EL), and HiRES120 (Advanced Bionics); see Supplementary Fig. 2 for descriptions of the algorithms and Supplementary Fig. 3 for example electrodograms. Previous comparisons of these devices have found no evidence of differences in overall speech comprehension, but have been limited to clean speech^69,70^. It remains possible that the ability to detect differences between devices was limited by the variability across human participants, or narrowly focused experimental assessments, and that performance differences would be evident for optimized decoders, for noisy conditions, or for sound localization. We optimized models for each of the three sound coding strategies, for both the word recognition and sound localization tasks (Fig. 5B).

**Fig. 5.**
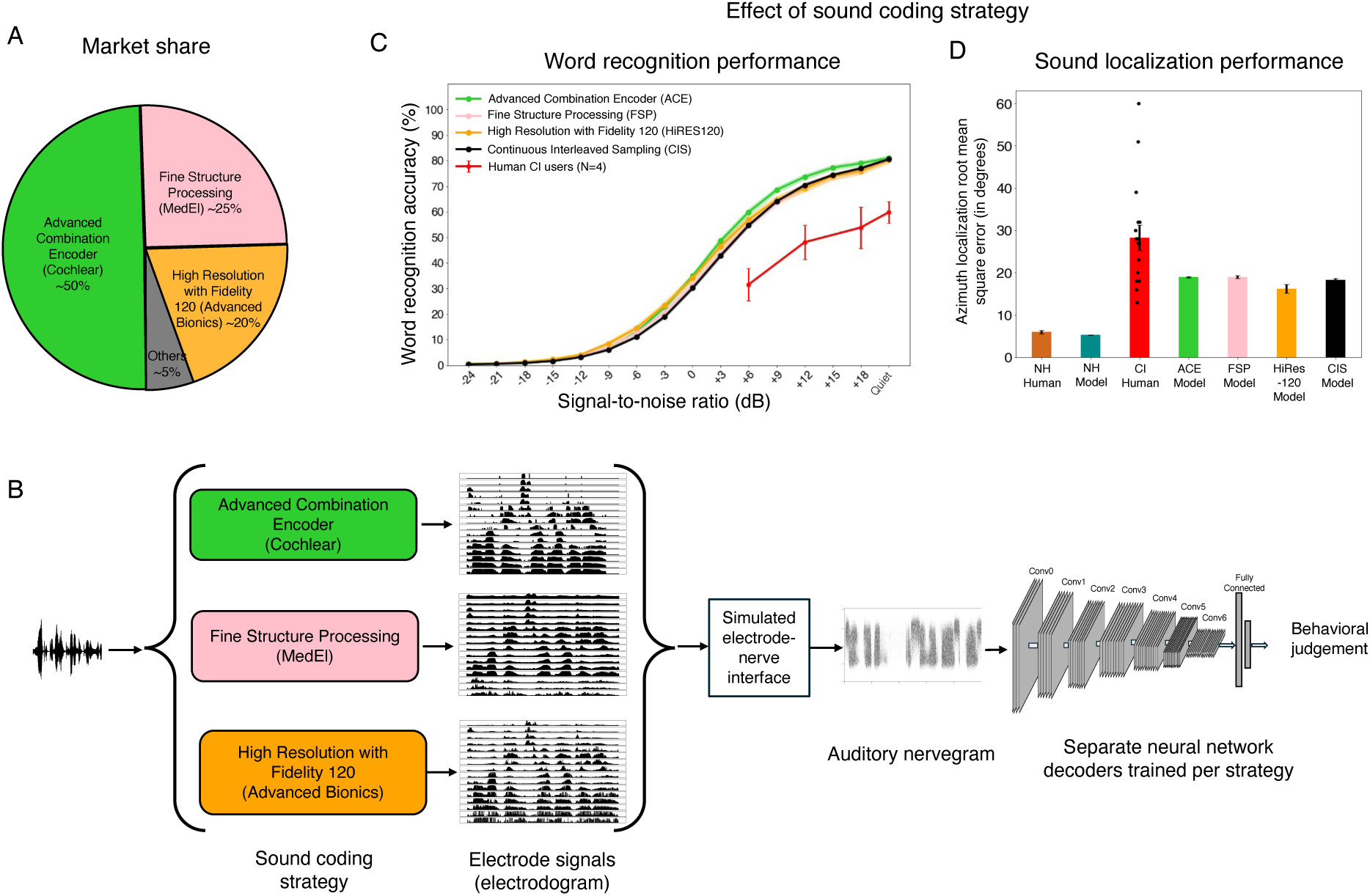
Similar word recognition and sound localization outcomes across leading sound coding strategies. **A.** Market share for the three leading device manufacturers. **B**. Models were optimized for each of the three main currently commercially available sound coding strategies: Advanced Combination Encoder from Cochlear, Fine Structure Processing from MED-EL, and High Resolution with Fidelity 120 from Advanced Bionics. **C.** Model word recognition performance across the three sound coding strategies compared to the baseline CIS strategy. Here and in D, error bars plot standard error across model architectures, and human data are replotted from Figure 3 and 4. Red line plots mean of the four human participants shown individually in Fig. 3B. **D.** Model sound localization performance across the three sound coding strategies compared to the baseline CIS strategy. Black dots plot performance for individual human cochlear implant users.

For both tasks, model results were quite similar across the three sound coding algorithms in use by the three major cochlear implant manufacturers. With optimized decoders, models showed comparable word recognition performance irrespective of the sound coding algorithm (Fig. 5C), as well as comparable sound localization (Fig. 5D). This result suggests that there is no major advantage to any of the three currently available device types for either speech recognition or sound localization.

### Factors limiting cochlear implant outcomes

We next sought to explore the factors that limit cochlear-implant-mediated auditory behavior. For both tasks, the performance gaps of cochlear implant models relative to normal hearing models suggest limitations in the way the device encodes sound. However, the fact that only the very best participants approach the cochlear implant model’s performance raise the possibility of limitations in the brain. Both types of factors have been widely discussed in the cochlear implant literature^15,71,72^, but the modeling framework introduced here provides a new avenue for testing them.

### Current cochlear implants do not support the use of interaural time differences

Human sound localization is known to be dependent in part on differences in time and sound level between the sound arriving at the two ears (Fig. 6A&B). Interaural time and level difference cues are complementary, being most reliable at low and high frequencies, respectively (“duplex theory”^73^). Cochlear implant users are thought to have poor sensitivity to time difference cues because of limitations in implant processing strategies. Specifically, cochlear implants could in principle encode interaural time differences via the timing of individual pulses (the electrodogram “fine structure”), but most current devices encode time information only in the electrode envelope (Fig. 6B). The one exception is the FSP strategy by MED-EL, which varies pulse timing in apical electrodes, but it remains unclear whether this information is usable. Prior behavioral studies on cochlear implant users have shown that cochlear implant users are sensitive to level differences but not time differences^71,74^. We sought to assess whether this cue dependence was present even with an optimized decoder for sound location. Such a result would suggest that there is little information to exploit in the envelope time differences that are in theory provided by current implants (because otherwise the optimized decoder would learn to exploit it).

**Fig. 6.**
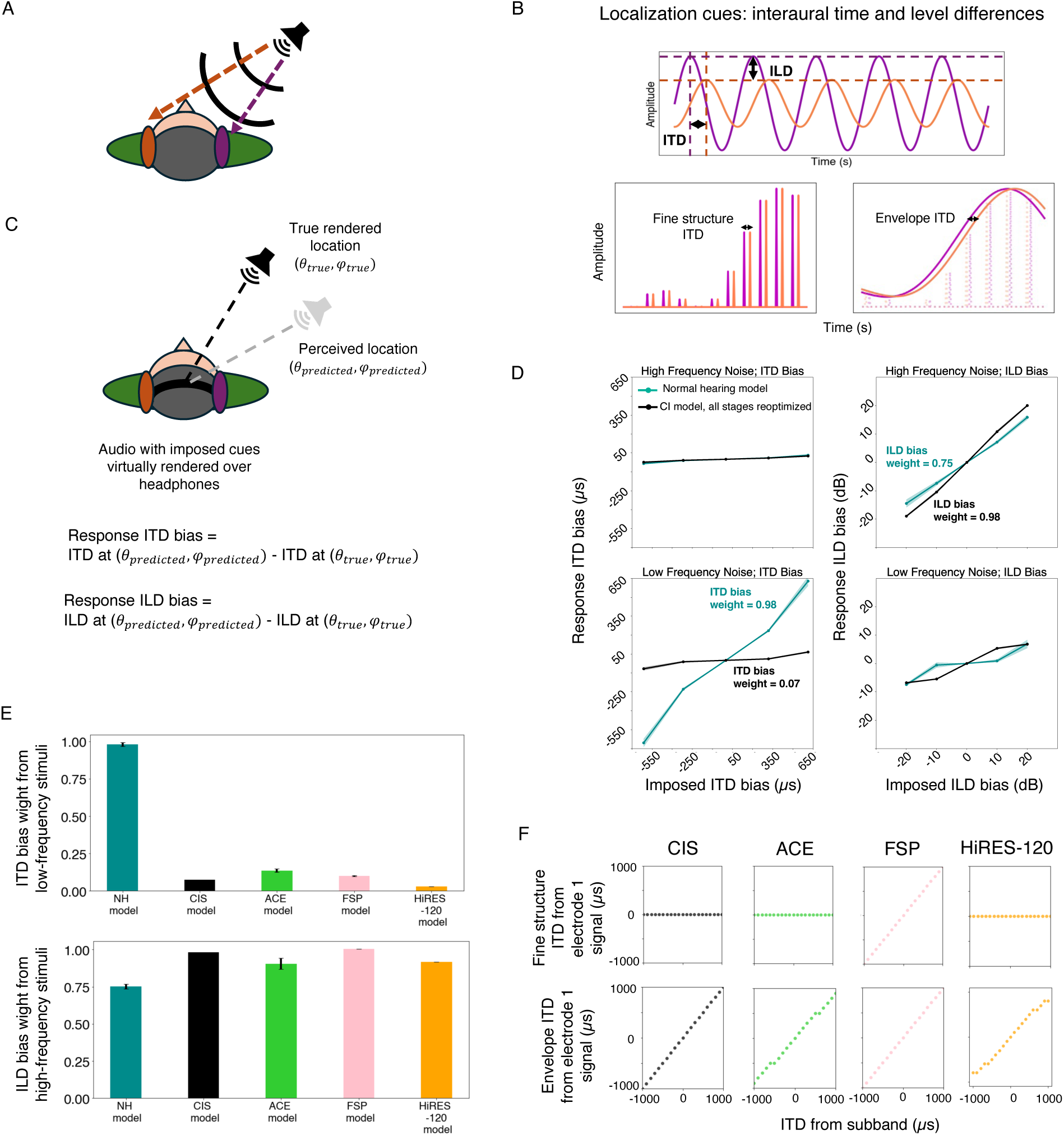
Models exhibit sensitivity to interaural level differences but not interaural time differences. **A.** Sound localization cues available to human listeners. **B.** Top: Interaural time and level difference (ITD and ILD, respectively) used for azimuth localization are shown for pure tones recorded at the left and right ears. Bottom: Temporal fine structure and envelope ITD cues as could be encoded by cochlear implant electrode pulse trains in the left (orange) and right (purple) ears. Right panel shows how ITD could be encoded in the timing of electrode pulses. Left panel depicts electrode signals from continuous interleaved sampling, in which pulse timings are uninformative, but in which the electrode envelope can encode ITD. **C.** Macpherson and Middlebrooks (2002) cue weighting experimental setup. **D.** Perceived ITD and ILD bias in response to imposed ITD and ILD bias measured for the normal hearing and cochlear implant models using low-frequency and high-frequency noise stimuli. Here and in E, error bars plot standard errors across model architectures. **E.** Comparison of ITD and ILD bias weights between normal hearing and cochlear implant models with different sound coding strategies. Graphs plot ITD bias weights measured for low-frequency stimuli and ILD bias weights measured for high-frequency stimuli, respectively. **F.** Envelope and fine structure ITD cues encoded by an example apical electrode (electrode 1) for different strategies. X-axis shows the ITDs obtained from the corresponding subband signal and Y-axis shows the ITDs obtained from the fine structure or envelope of the electrode signals. Co-variation between the stimulus ITD and the electrode ITD indicates that the electrode encodes the stimulus ITD.

We simulated the experimental paradigm of Macpherson and Middlebrooks (2002)^73^, who analyzed interaural time and level difference sensitivity in normal hearing listeners. As depicted in Figure 6C, participants heard noise bursts with artificially imposed additional ITDs or ILDs and reported the location of the sound source. The responses were then analyzed to determine if the added ITD or ILD influenced the perceived location (quantified as the change in ITD or ILD that would normally correspond to the shift in location). Macpherson and Middlebrooks tested both low-and high-frequency noise bursts, finding that ITDs influenced the localization of low frequencies, while ILDs influenced the localization of high frequencies, consistent with classical duplex theory. These effects have previously been shown to also occur in neural network models optimized to localize sound given simulated normal cochlear input^19^. We ran the experiment on models optimized with the classic CIS strategy along with the ACE, FSP, and HiRes120 strategies used in current devices.

A priori it was not completely clear what to expect. Models optimized for sound localization with limited auditory nerve phase locking show no sensitivity to interaural time differences^20^, but also have the benefit of normal level difference cues, as well as normal spectral cues from pinna filtering. If time differences between the envelopes of the left and right ear input are useful for localization, an optimized decoder might learn to use them. It also seemed plausible that the fine structure information encoded by the FSP strategy would provide usable cues. The Macpherson and Middlebrooks experiment should reflect both sensitivity to envelope and fine structure ITDs, as both are present in the noise stimuli that were used.

As shown in Fig. 6D, the cochlear implant model had negligible sensitivity to ITD cues in either frequency range despite near-normal sensitivity to ILD cues for high frequencies. We quantified this dependence as the cue bias weight (see Methods for details), and used this to summarize the experiment result for CIS as well as the three current stimulation strategies. The ITD cue weight was close to zero for CIS as well as all three contemporary sound coding strategies (ACE, FSP, HiRes120; Fig. 6E and Supplementary Fig. 4).

To confirm that ITD information was present in the electrode responses of the cochlear implant simulations, we measured both envelope and fine structure ITDs from left-right pairs of individual electrodes for each of the four strategies, using bands of noise centered at each electrode and spatialized to different positions. Results for an example apical electrode are shown in Fig. 6F. As expected, all four stimulation strategies convey ITDs in the envelope of the electrode response when such ITDs are present in the acoustic stimulus. Moreover, the FSP strategy also conveys ITDs via the fine structure of apical electrode responses (i.e., the pulse timings), whereas the other strategies do not (again as expected). See Supplementary Figs. 5-7 for results from all electrodes.

Overall, these results suggest that the ITD cues conveyed by current cochlear implants (even the fine structure ITDs conveyed by the FSP strategy) are not sufficient to substantially aid localization^75,76^. Even though the cues are present in the electrodes, they do not contribute to the localization behavior of a decoder that is optimized to work with them. Two factors seem likely as root causes of this limitation: one is that the cues are corrupted by the electrode-nerve interface, such that cues that are evident in the electrodes are not extractable from the nerve. The other is that cues that are evident in the clean conditions we analyzed in Fig. 6F are not robust to realistic conditions with noise and reverberation, and so are not used by a decoder optimized for realistic conditions.

### Model performance is limited by spatial spread of cochlear stimulation

Another device constraint often thought to limit behavioral outcomes is the extent of spatial spread across the cochlea of the electrical field from an electrode, as this plausibly limits the spectral resolution with which sound can be represented in the auditory nerve. The model results shown thus far were intended to match spatial spread appropriate for monopolar stimulation used in most current devices^77^, but the spread is thought to vary somewhat between devices, and between electrodes, based on the distance of the electrode from the nerve, and the current level^78^. However, because it is difficult to measure the spread of excitation in an individual implant user^79–81^, and because tests of current spread manipulations in humans are typically restricted to isolated sessions (rather than letting a human live with decreased spread for a period of time before testing its effects), it is not clear how much the spread in fact limits behavioral performance^82–85^.

To quantify what effect such variation might have on performance, we separately trained models with spread thought to be appropriate for bipolar and tripolar stimulation, which is substantially narrower than monopolar stimulation^77,86^ (Fig. 7A), but not usually used in commercial devices because of power consumption concerns and device voltage limits. We also hoped that this degree of difference in spread might roughly capture the range of spreads that can occur for different electrode conditions^44,87,88^. As shown in Fig. 7B, decreasing the spread of excitation improved model word recognition, substantially narrowing the gap with the normal hearing model (the SNR yielding 50% correct decreased by 3.11 dB for bipolar and 4 dB for tripolar). By contrast, there was little effect of current spread on sound localization (Fig. 7C), potentially because when localizing a single sound in the horizontal plane, the cues are fairly consistent across frequencies, and so are not degraded much when spectral resolution is decreased. This result confirms the expectation that spatial spread is a fundamental limitation of current cochlear implants, and suggests that reducing the spread with alternative coding strategies^86^ might produce measurable benefits in performance.

**Fig. 7.**
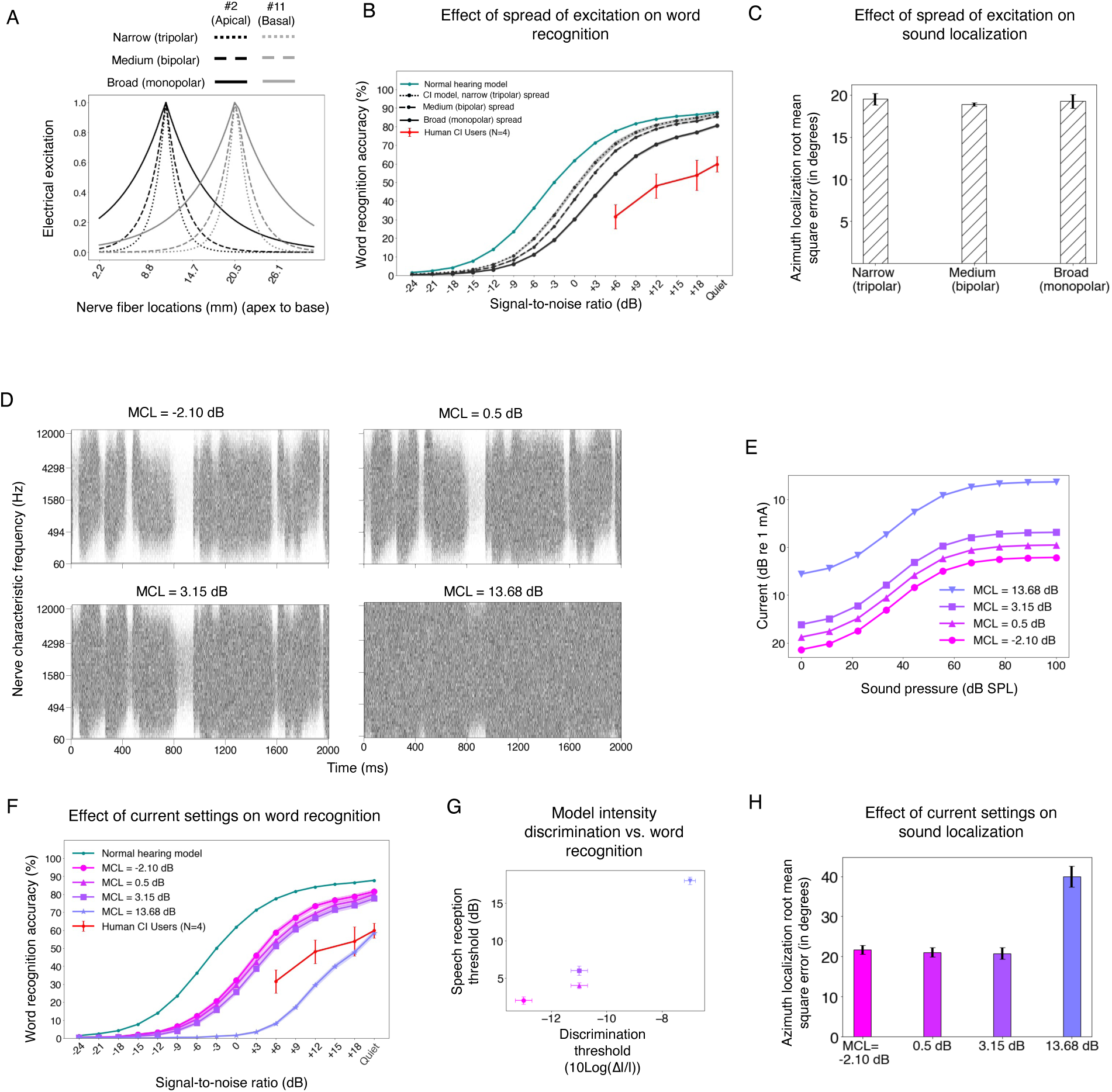
Effects of current spread and current levels on word recognition and sound localization performance. **A.** Illustration of three different extents of spatial spread of electrical excitation. X-axis shows nerve fiber locations (in mm) along the cochlea. Y-axis shows the strength of the electrical field at each location. **B.** Word recognition performance for different spatial spreads of excitation. Here and in F, red line plots mean of the four human participants shown individually in Fig. 3B. **C.** Noise burst localization performance for the same three spatial spreads of electrical excitation in B. **D.** Example auditory nerve responses to an excerpt of speech for four different current setting choices. For current level settings that yield worse intensity discrimination than that of human implant users (bottom right), the speech signal appears less evident in the nerve response. **E.** Loudness growth functions for three current settings that produce human-like intensity discrimination (MCL values closer to 0 dB) and one current setting that produces substantially worse intensity discrimination than that of human implant users (MCL value significantly higher). **F.** Word recognition performance with the four current settings from C and D. Error bars plot standard error across model architectures. **G.** Scatter plots of model speech reception thresholds vs. intensity discrimination thresholds. **H.** Noise burst localization performance for the same four current level settings in F.

### Sub-optimal choices of current levels affect recognition and localization performance

Another device-related factor limiting human outcomes could be the way in which the stimulation current range is set. We found in Fig. 2 that some current-level choices resulted in intensity discrimination thresholds similar to those of human cochlear implant users, but that sub-optimal choices resulted in much higher thresholds. However, it was unclear how these current level choices would translate to the performance of real-world auditory tasks. We assessed their effect on speech recognition and sound localization by separately optimizing models with current levels that either yielded human-like intensity discrimination, or that produced substantially worse intensity discrimination (Fig. 7D&E).

As shown in Fig. 7F&G, current level choices that produced good intensity discrimination also yielded good word recognition performance. Similar results were observed for sound localization (Fig. 7H). Moreover, the one current setting we tested that produced substantially worse intensity discrimination than humans also produced substantially worse word recognition and sound localization performance compared to the other current settings. This result is potentially clinically relevant, in that it suggests that an audiologist does not need to optimize current settings separately for speech recognition provided they afford good intensity discrimination.

### Model-based tests of limitations on speech recognition

We next considered the factors internal to the auditory system that might account for suboptimal cochlear-implant-mediated speech recognition in humans relative to optimized decoders. We investigated two commonly cited such factors: limited plasticity and neural degeneration, both of which can be simulated in a model. It remains possible that the simulation of these factors deviates in important ways from the brain, but it nonetheless provides an example of their effect in a working system that solves some of the tasks facing humans, which could provide useful clues.

### Model with limited plasticity replicates speech recognition of human implant users

The improvement of human speech recognition scores following implantation^15^ suggests an important role for learning to decode information from the implant, but learning might be limited. In particular, there is some reason to think that early stages of sensory systems are less plastic than later stages^89–92^. However, the extent to which plasticity limits behavioral outcomes remains uncertain, particularly in comparison to other factors such as neural degeneration. Inspired by evidence of coarse correspondence between stages of feedforward artificial neural networks and stages of the auditory system^17,62^, we simulated the possibility of plasticity that is restricted to later stages of the system by varying the number of decoder stages that were reoptimized, in each case retraining the model until asymptotic performance was reached (Fig. 8A). The approach embodies the hypothesis that much of experience-induced plasticity following cochlear implantation can be modeled by reoptimization for behaviorally important tasks, potentially with anatomical constraints on the locus of plasticity. Each model stage was either fully reoptimized (i.e., until asymptotic performance was reached) or not optimized at all. We also included a condition in which no stages were reoptimized, which might simulate a user’s performance of an auditory task immediately after the activation of the implant, or in a scenario in which plasticity is not possible.

**Fig. 8.**
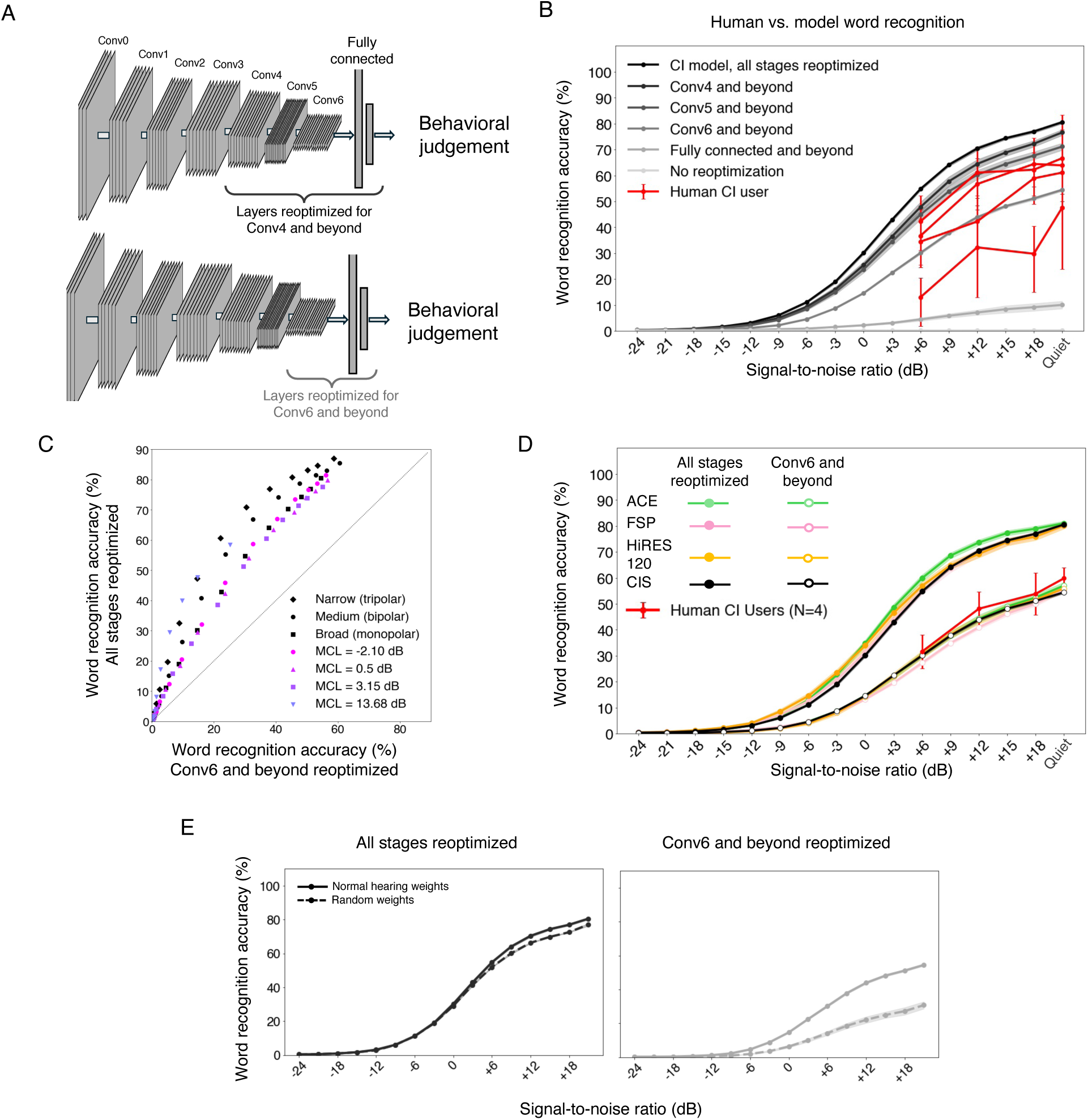
Model behavior is critically dependent on plasticity and benefits from normal-hearing weight initialization. **A.** Schematic of two different plasticity conditions: one in which Conv4 and beyond were re-optimized (top) and one in which Conv6 and beyond were re-optimized (bottom). **B.** Word recognition accuracy as a function of the signal-to-noise ratio for human cochlear implant users (N=4; data replotted from Fig. 3B), and models with different plasticity cutoffs. Error bars plot standard error across either model architectures or stimuli (for human participants). **C.** Comparison of model word recognition performance for two plasticity conditions with different device settings and stimulus SNRs. The performance improvement for the fully plastic decoder is fairly consistent across conditions. **D.** Model word recognition performance for the three leading sound coding strategies in current use, for full and partially plastic decoders. The sound coding strategies produce similar results for both types of decoders. Red line plots mean of the four human participants shown individually in Fig. 8B. Error bars plot standard error across either model architectures or human participants. **E.** Word recognition performance for models with normal hearing weight initialization (solid lines) compared to that for models with random initialization (dashed lines), for two plasticity conditions.

As shown in Fig. 8B, the extent of model plasticity had an enormous effect on the model’s word recognition performance. Without any reoptimization, the model’s performance was at floor. Increasing the number of reoptimized stages increased performance. This result is consistent with the idea that the wide range of outcomes observed in human cochlear implant users could reflect variation in the extent of brain plasticity. We note that the near-chance model performance without any reoptimization might be viewed as inconsistent with human cochlear implant users who have some ability to recognize speech immediately following the implant activation^15^ (along with many users whose performance is very poor immediately after activation, and who eventually achieve much better recognition). It is possible that there is some form of very rapid plasticity in biological auditory systems (perhaps more akin to gain control) that could help reconcile these results (see Discussion).

We trained partially plastic models with several current settings and with both broad and narrow spread (limiting reoptimization to the Conv6 stage and beyond), and the effect of partial plasticity produced a consistent decrease in performance (Fig. 8C). We also trained partially plastic models with each of the three leading sound coding strategies in current use, and found that they again yielded similar results (in each case performing worse than if the decoder was fully reoptimized; Fig. 8D). These results indicate that conclusions drawn with fully optimized decoders about the relative performance for different device settings are likely to apply to a considerable extent in conditions where the decoder is not fully reoptimized. This result is practically relevant as it suggests that using models to search for new coding strategies that improve best-case scenario outcomes should produce benefits even in conditions when plasticity is not as extensive.

### Partially plastic models show benefit of normal hearing weight initialization

The effects of limited plasticity interacted with whether the decoder weights had been initially optimized with normal hearing input. We compared two sets of models optimized for cochlear implant input: those initialized with weights that had been optimized for normal hearing input, and those with randomly initialized weights. When the model was fully plastic, there was minimal benefit to normal hearing weight initialization over random weight initialization (Fig. 8E). However, when model plasticity was limited to late model stages, normal hearing weight initialization yielded better word recognition performance than random initialization, indicating that early processing stages learned for normal hearing have some benefit to cochlear implant decoding. This result is qualitatively consistent with evidence that normal auditory experience prior to deafness can in some cases improve cochlear implant outcomes.

### Simulated neural degeneration yields only modest degradation in speech recognition

Another factor that could account for the gap in human cochlear implant user performance and that of the fully re-optimized model is neural degeneration. Extended periods of deafness are believed to lead to some degree of neural degeneration, particularly in the auditory nerve^9,10,93^, but the effect on behavior is uncertain, in part because there is currently no way to measure degeneration in humans while they are alive. We modeled peripheral neural degeneration by silencing a fraction of nerve fibers in the input to the neural network (Fig. 9A). The silenced fibers were randomly selected from the entire frequency range. In a separate condition we also modeled central neural degeneration by randomly silencing a portion of the frequency-channel combinations in each stage of the neural network (Fig. 9B), conceptualizing these “units” as loosely analogous to neurons. The resulting models were reoptimized, either entirely (end-to-end), or with reoptimization restricted to late model stages (to assess whether degeneration effects might be more pronounced in this setting). We simulated loss of 90% of auditory nerve fibers, along with a loss of 50% of central neurons. The extent of degeneration is likely to vary depending on the duration of deafness, but such losses are consistent with the upper end of documented neural loss in cases of prolonged periods of deafness^93–96^.

**Fig. 9.**
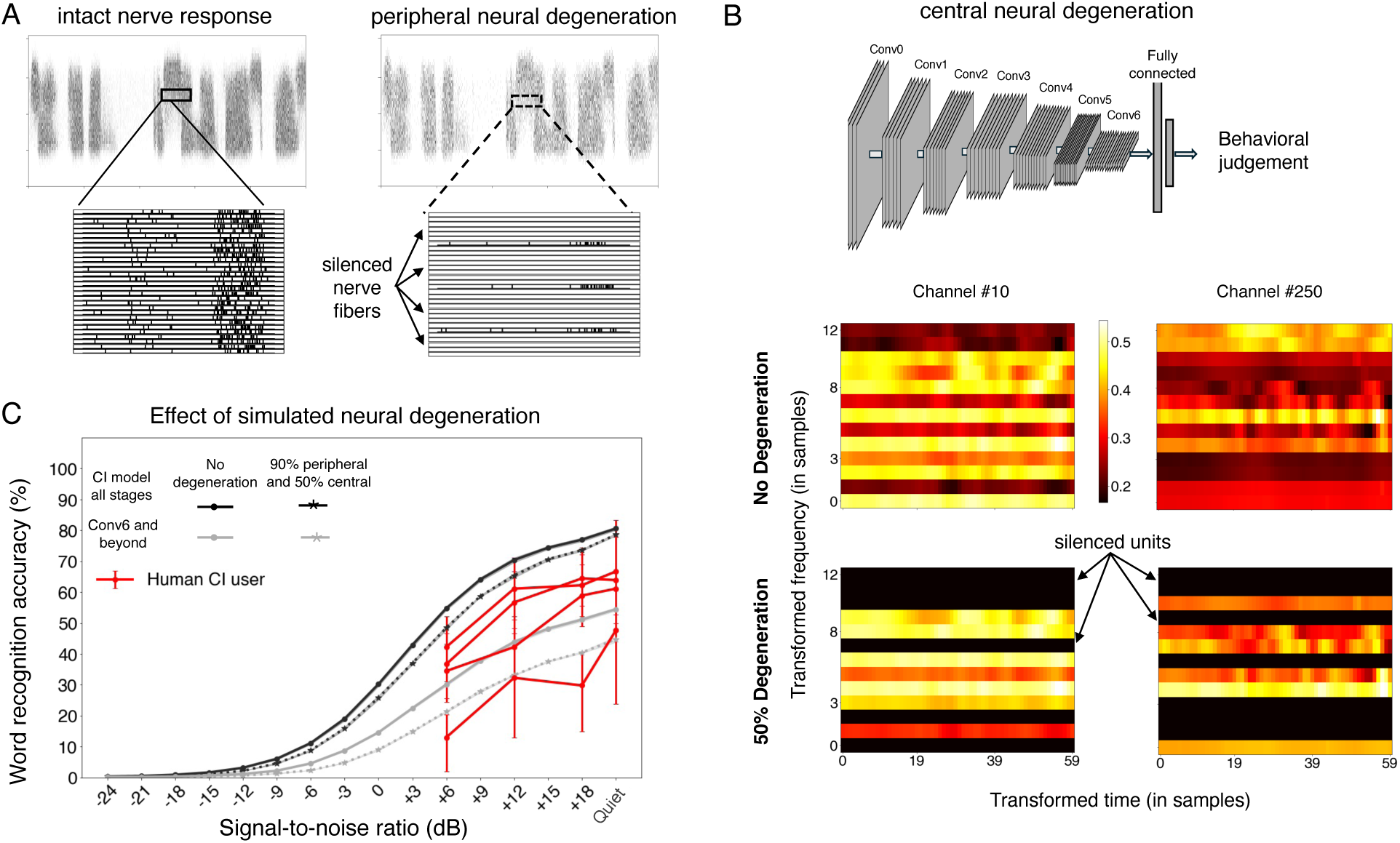
Neural degeneration has modest effects on speech recognition outcomes. **A.** Example of simulated peripheral neural degeneration, showing 90% of nerve fibers silenced. In this condition, fibers were silenced both during reoptimization and evaluation) **B.** Example of simulated central neural degeneration depicted for one stage (Conv3). For the 50% degeneration condition, 50% of the units (chosen randomly) are silenced at all times (both during the model reoptimization and evaluation). Color denotes activation. **C.** Model word recognition performance with and without 90% peripheral and 50% central degeneration (dashed and solid lines, respectively) across two different plasticity conditions. Error bars plot standard error across model architectures or stimuli (for human participants). Human data replotted from Fig. 3B.

In contrast to the large effect of simulated plasticity, the effect of simulated neural degeneration was modest. Even with 90% nerve degeneration, model word recognition performance was only slightly impaired (Fig 9C, solid lines vs dotted lines). With a fully reoptimized decoder, models with degeneration outperformed most human users. This result raises the possibility that neural degeneration on its own cannot explain limits on human outcomes, or the extent of individual differences in outcomes observed in human implant users (see Discussion).

Overall, the results of these experiments show that the factors commonly proposed to constrain human perception through cochlear implants also have some effect on a working computational model.

## Discussion

We developed a computational framework to understand sensory prostheses and applied it to cochlear implants. We optimized neural network decoders of simulated cochlear implant input for the tasks of recognizing speech in noise and localizing natural sounds. We first assessed best-case scenario outcomes for each task by fully reoptimizing a decoder for cochlear implant input and comparing these results to those of human cochlear implant users. For word recognition, fully reoptimized models produced good overall performance in quiet (i.e., close to normal-hearing performance), but were less noise-robust than models with normal hearing input. For sound localization, performance of fully re-optimized models was relatively poor even in quiet. For both tasks, the model performance was on par with the best-performing human cochlear implant users. We also compared best-case scenario outcomes for the three stimulation strategies currently used by the three main cochlear implant manufacturers and found that they all yielded similar results. We then investigated device limitations that might limit behavioral performance. Sound localization limitations were explained by the absence of useful interaural time difference cues in cochlear-implant-stimulated nerve responses, and performance limitations for word recognition were influenced by both spatial spread of stimulation and suboptimal current settings. These findings provide new evidence that improvements to each of these three factors should produce better outcomes, by showing their effect on behavior in a model that performs real-world tasks. Lastly, we investigated the factors that might account for the performance gap between some human implant users and the best-case scenario revealed by the fully reoptimized model. These experiments indicate that reoptimizing the decoder for cochlear implant input was critical – reducing the number of plastic stages eventually brought performance to near-chance levels. By contrast, degeneration produced less variation in outcomes. The results clarify the factors limiting prosthetically mediated perception, and illustrate a framework for predicting real-world behavioral outcomes from candidate prostheses, opening the door to large-scale screening of new device strategies.

### Relation to prior work

We built on a substantial prior literature developing models of how electrical stimulation influences auditory nerve responses^26,97–100^. Relative to this work, our contribution was to build models that mediate real-world behavior using optimized decoders, allowing investigation of the factors that limit behavior. This was only possible given contemporary machine learning methods (previous types of models would not have been able to approach human levels of performance on real-world tasks). Two previous papers have also attempted to use neural networks to predict speech recognition from simulations of cochlear implant input. The first of these used a neural network to predict phonemes from a simulation of a cochlear implant, and reported that the model performed similarly to human cochlear implant users^31^. A second paper also trained a speech recognition system on a cochlear implant simulation^32^. However, the simulation was unconventional, involving noise vocoding of sound followed by a spectrogram transformation, and it was not clear that it accurately captured the information in a typical cochlear implant. Our contribution relative to this work is to optimize models with realistic nerve simulations and to use the models to characterize the factors that determine human behavioral performance.

Another line of prior work has attempted to understand the limitations of cochlear implants using noise-vocoded sounds presented to normal hearing listeners^101,102^. Such papers have reported deficits of various sorts for recognition of noise-vocoded compared to natural speech, attributing them to the information believed to be discarded by cochlear implants. However, we found that a model optimized for normal-hearing nerve input performed on par with normal-hearing humans when presented with noise-vocoded speech (i.e., well above chance; Supplementary Fig. 1), but performed at chance for simulated cochlear-implant-stimulated nerve input (Fig. 7B). This difference plausibly reflects the substantial difference in firing rate statistics between the two types of stimulation (Fig. 2C), such that a decoder optimized for one type of nerve stimulation does not necessarily transfer to the other. Noise vocoding may capture aspects of the limited resolution of current cochlear implants, but likely does not produce the same amplitude statistics in the auditory periphery as sound heard through a cochlear implant. The result suggests that noise-vocoded speech heard by normal hearing listeners should not be taken as a complete proxy for a cochlear implant user presented with natural speech.

Prior models of sound localization via cochlear implants have also provided insight, but to the extent that they have been applied to behavioral judgments, they have been dependent on ad hoc decision stages operating on hard-coded binaural circuitry (receiving simulated cochlear implant input)^103,104^ . As such, they have been unable to model localization of natural sounds, or to explore what is possible with an optimized decoder. By leveraging machine learning (and thus being able to mediate localization behavior for real-world sounds and conditions), we were able to compare humans to an optimized localization system, providing more definitive evidence that factors internal to the brain are not the main limitation for sound localization via current cochlear implants. We were also able to better understand why the residual time difference cues that are preserved by current implants do not meaningfully support localization (in showing that the model did not learn to use them even when optimized for the task, presumably because they are not informative in natural conditions). These results complement the experimental literature that links localization deficits in human cochlear implant users to impaired interaural time difference sensitivity, by linking the deficits to current device strategies (we note that the results also do not preclude factors internal to the brain that might limit behavior with alternative sound coding strategies).

### Limitations

Modeling biological phenomena with mathematical operations necessitates assumptions and approximations. Most notably, we relied on simulations of the process by which a cochlear implant produces action potentials in the auditory nerve. This “electrode-nerve interface” in our models is not completely accurate, in part because our implementation was constrained by the computational demands of training models at a scale sufficient to simulate real-world behavior, and by differentiability (to aid future device optimization that might build on the framework described here). Two notable inaccuracies involve modeling the nerve response as a linear time-invariant system (lacking refractoriness or adaptation), and treating nerve fibers as being excited at only a single point in space rather than potentially at different points along a fiber. These inaccuracies prevent us from modeling some known details of electrically driven nerve stimulation, including the ineffectiveness of higher pulse rates (which likely results in part from refractoriness of nerve fibers). We also made assumptions for some aspects of the interface that likely do not hold in all cases. For instance, we assumed that the electric field from each electrode decayed exponentially at the same rate at all electrode locations, whereas in an actual implant variation in electrode placement and impedance likely introduces some variation across electrodes. Our framework allows such variation to be modeled, but we did not explore it.

The neural network architectures were also constrained by assumptions that could be limiting. For instance, they were strictly feedforward, and lacked the feedback and lateral connectivity present in biological sensory systems. We also built separate models for word recognition and sound localization. Humans, by contrast, perform both tasks using one brain (though there may be considerable segregation of task-specialized representations for the two tasks). It is possible that having to perform both tasks with some of the same neural circuitry constrains the possible task solutions in ways that did not affect our models. The tasks themselves were simplifications of real-world perception. The word recognition task had a large vocabulary, but was still much smaller than the full English language, and required identifying the middle word in a speech excerpt rather than continuous recognition. Both tasks were also restricted to single sources, and did not allow for attentional selection^105^.

Our simulation of neural degeneration was also impoverished. We simulated degeneration as all or none, by setting nerve fibers or units to zero. In actuality, degeneration can be partial, with some nerve fibers remaining somewhat responsive to sound despite having compromised functionality^106^. Our simulations also assumed that degeneration was spread evenly across frequency. Our framework again allows for alternatives to be tested, but we did not do that here.

Plasticity in our models was also limited, being exclusively captured by optimization. Although the plasticity that occurs in cochlear implant users does serve overall to substantially improve task performance, it is not clear that it is well approximated by gradient descent on a task-based objective. Sensory systems likely also exhibit homeostatic and Hebbian plasticity, for instance, the behavioral effects of which remain unclear. Moreover, some plasticity following hearing loss is plausibly maladaptive, including increased neural activity and synchrony, disruption in excitatory-inhibitory balance, and anti-Hebbian plasticity^107^.

### Consequences of model limitations

We next discuss whether these assumptions and approximations limit the conclusions that can be drawn from our results, considering each of the main results in turn. Our first main result concerned best-case scenario performance levels. We found that fully reoptimized decoders for both word recognition and sound localization underperformed models of normal hearing, but substantially outperformed all but the very best human cochlear implant users. In principle inaccuracies in the electrode-nerve interface simulation could cause the performance of the fully reoptimized model to be an overestimation of what is possible given actual nerve responses. In our view this is not likely because we exclusively studied contemporary stimulation strategies, all of which have fairly low pulse rates, for which the most substantial inaccuracy in this model stage (lack of refractoriness) is unlikely to substantially affect nerve responses^108^. It is also difficult to see how the other inaccuracies and approximations would result in a substantial overestimation of performance. It is also possible that the decoder architecture was somehow limited, producing an underestimation of performance, but that seems unlikely given that the same architectures achieved human-like performance when optimized with normal-hearing input. The conclusions about best-case scenario performance thus seems likely to generally hold.

Our second main result was that word recognition and sound localization performance were not substantially different for the three main stimulation strategies in widespread use. This conclusion also seems unlikely to depend sensitively on our modeling assumptions. Because the pulse rates of all three strategies were low, the main inaccuracy in the electrode-nerve interface (lack of refractoriness) is unlikely to affect one strategy more than another. We note that we did not evaluate each strategy with all possible settings of some of the other variables that can affect performance (e.g. current spread and degeneration), and it is possible that a strategy could interact with these variables in some way that could produce larger differences in performance. But the conclusions for a situation without other aggravating factors seem sound (and we note that we obtained similar results for models that were only partially optimized; Fig. 8D).

Our other main results concerned internal factors that limit performance. Two of these results in particular seem to us to be insightful but provisional. The first was that performance depends dramatically on plasticity in the decoder. This result is a function of the decoder for normal nerve responses, which depends on the neural network architectures we used, the optimization procedure used for training, and the training data. In principle some alternative optimization procedure and/or model architecture could result in a normal hearing decoder that generalizes better to cochlear implant input, in which case plasticity might not be as critical to obtaining good performance. At present the possibility cannot be fully excluded, and we thus view this result as suggestive rather than definitive.

A second internal factor we tested was neural degeneration, which had only a modest deleterious effect on performance. We explored only a few configurations of degeneration, but the amount we tested was fairly substantial (Fig. 9; 90% peripheral loss with 50% central loss). The extent of degeneration that we considered is on the far end of what is believed to occur in typical cochlear implant users^109^, and the modest results are consistent with studies suggesting that degeneration (or other changes, such as demyelination) have to be severe in order to produce behavioral deficits^109,110^. However, it seems possible that degeneration could have other effects we did not model, such as plasticity that is partially maladaptive^107,111,112^, which might induce further impairments. For this reason, we regard any conclusions about degeneration to also be suggestive rather than conclusive.

### Future directions

The present results show how machine-learning-based models can be used to understand best-case scenario outcomes for candidate prosthetics, setting the stage to use such models to explore alternative device algorithms^113^. One important avenue will be to evaluate some of the recent proposals for alternative sound coding strategies^114–123^, which have not been tested comprehensively in humans due to the impracticality of large-scale long-term trials. Another will be to use our model framework to optimize sound coding algorithms for task performance. The models presented here were constructed entirely from differentiable components, making it possible to propagate gradients all the way back to the device. Direct optimization of the signal processing in a device should in principle enable better behavioral performance. In both cases the modeling framework can be used to screen candidate device algorithms, the most promising of which might then justify a clinical trial in humans. One goal of such efforts could be to develop new sound coding strategies that improve sound localization abilities while maintaining good speech recognition. Another could be to develop strategies that improve pitch and music perception. A third possibility is to optimize a strategy for unilateral deafness, in which an implant-aided works in concert with a normal ear, or an ear with a hearing aid.

Our work here suggests an important role for plasticity in cochlear implant outcomes. More detailed comparisons between human behavior, brain responses to cochlear implant stimulation, and models of cochlear-implant-mediated hearing have the potential to provide a test bed for theories of plasticity. Such investigations will benefit from models that are more faithful to biology. For instance, we found that performance approaching that of humans was best achieved when only a subset of model stages was reoptimized for cochlear implant input. Such an arrangement could in principle correspond to plasticity limits in early stages of the auditory pathway. Building models that remain optimizable while being constrained to more closely replicate the biological auditory system (e.g. in connectivity, or receptive field extents) could help to more precisely test such ideas.

The constraints imposed by information loss, brain plasticity, and sensory system damage are also relevant to retinal and auditory brainstem implants, which are in earlier stages of development compared to cochlear implants. The approach illustrated here should be equally applicable to these other types of sensory prostheses, and could potentially help guide their development.

## Methods

### Simulation of normal-hearing auditory nerve response

Sound waveforms were first passed through a bank of bandpass filters intended to simulate the filtering imposed by the cochlea. The filters had center frequencies ranging from 60 Hz to 12 kHz. The filters had rounded exponential transfer functions^124^ placed uniformly on an equivalent rectangular bandwidth scale^125^. Filters were applied in the frequency domain via multiplication and resulted in 50 subbands.

The choice of 50 frequency channels resulted from practical constraints. The filter bank was applied at run time as part of the larger model implementation, and memory and compute constraints precluded a larger number of channels. Previous work^18^ has found that increasing the number of frequency channels by a factor of ten did not substantially alter model behavior.

The subbands were rectified and low pass filtered with a cutoff frequency of 3000 Hz (using a finite impulse response filter designed using a Kaiser window of length 50 ms, made using the “firwin” function in the scipy signal library) to simulate the upper limit of phase locking observed in auditory nerve fibers^126–128^. The result of these operations was passed through sigmoidal rate-level functions intended to simulate the response properties of auditory nerve fibers. The lower and upper asymptotes of the sigmoid curve corresponded to the spontaneous and maximum firing rates, respectively. We simulated three different types of auditory nerve fibers^33^: high-spontaneous rate, medium-spontaneous rate, and low spontaneous rate, with corresponding spontaneous rate values of 70.0, 4.0, and 0.1 spikes per second, respectively. The nerve fibers had thresholds of 0.0, 12.0, and 28.0 dB SPL, respectively, and dynamic ranges of 20.0, 40.0, and 80.0 dB respectively. All three fiber types had a maximum firing rate of 250 spikes per second. The rate-level function was given by

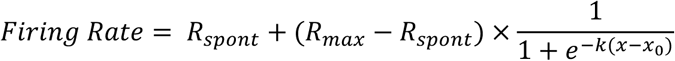

where x and x_0_ are in units of dB SPL, x_0_ = T + (D/2), T is the threshold in dB, D is the dynamic range in dB, and k = 7.32/D.

The resulting nerve firing rate representation was a three-dimensional tensor of shape N x T x C where N is the number of frequency channels (N=50), T is the total number of time samples (T=20000, with stimuli of length 2s and a cochlear sampling rate of 10 kHz) and C is the number of nerve fiber types (C=3). We then stochastically sampled spike trains (each corresponding to an individual nerve fiber) from these instantaneous nerve firing rates. We simulated ∼32,000 nerve fibers in total. At each frequency, we simulated 32000/N fibers, of which 60% were high-, 25% were medium-, and 15% were low-spontaneous rate fibers. This number was chosen to approximate the number of nerve fibers in the human ear. To implement efficient spike sampling, we assumed that each nerve fiber’s spikes result from a binomial process and that the responses of nerve fibers tuned to similar frequencies are summed prior to subsequent processing^129^. We approximated the sum of a set of binomial processes as a normally distributed random variable. Specifically, a sum of binomial processes with parameters n = (fiber-type-proportion * 32000/N) and p = instantaneous firing rate/sampling rate was approximated with a Normal(np, np(1-p)) distribution, samples from which were rounded to the nearest integer. The resulting three-dimensional tensor represents the total number of spikes generated at each time-frequency-nerve fiber bin. This auditory nerve representation was represented with a sampling rate of 10 kHz to reduce the dimensionality of the representation while avoiding information loss.

In previous work^20^, we found that this relatively simple model of auditory nerve responses produced similar behavior to a more complicated model^130^ that captures additional details of peripheral responses. We used the simpler model as it was fast enough to be computed on the fly as part of a larger model with a neural network back-end.

### Sound pressure level (SPL) of model input

The sound pressure level of input signals was defined as:

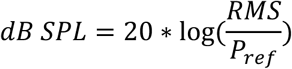

Where P_ref_ was set to 20 × 10^)0^ P.

### Simulation of cochlear-implant-stimulated nerve response

The simulation of CI-stimulated nerve responses involved three key components: (1) the CI signal processor, (2) the interaction between the CI electrodes and the auditory nerve fibers (electrode-nerve interface), and (3) the generation of auditory nerve spiking response following CI stimulation.

We first implemented the signal processing steps performed in an actual CI device to convert sound into electrical signals or electrodograms. Our baseline CI model implemented the continuous interleaved sampling (CIS) based sound coding strategy^38,131^, which modulates the amplitude of electrical pulse trains with the envelopes of a set of sound subbands. Sound waveforms were first passed through a bank of M half-cosine filters, with each filter providing the input to one electrode. Our default CI model used M = 16 electrodes. The center frequencies of the M filters ranged from 150 Hz to 10 kHz, with the filters evenly spaced on an equivalent rectangular bandwidth scale. Low-and high-frequency filters provided input to more apical electrodes and more basal electrodes, respectively. We extracted the envelopes from the M subbands using a rectifier and low-pass filter (with a cutoff of 50 Hz, implemented using a 50 ms Kaiser window made using the “firwin” function in the scipy signal library). We experimented with a higher envelope cutoff (of 160 Hz) and found this did not substantially improve performance. The envelopes were downsampled to the cochlear sampling rate of 10 kHz.

The envelopes were passed through a non-linear loudness growth function that mapped audio amplitudes to corresponding current levels defined by threshold level (TL) and maximum comfortable level (MCL) parameters (in dB re 1 mA). We used a sigmoidal function parameterized such that sound levels between 10 and 70 dB SPL were mapped to currents between TL and MCL. The upper asymptote of the sigmoid captured aspects of the automatic gain control common to CIs in the sense that levels above 70 dB SPL were compressed into a small range. The loudness growth function was specified as:

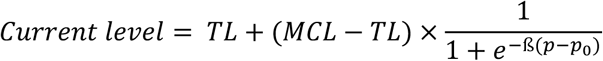

where p_0_ = T + (D/2) = 40 dB, with T being the threshold in dB SPL (T=10) and D being the dynamic range in dB (D=70 -10=60). ß = 0.122. The acoustic dynamic range of 60 dB matched that used in current cochlear implants^7^.

These nonlinearly transformed envelopes were used to amplitude-modulate pulse trains. In the CIS strategy, the pulses on different electrodes are interleaved in time to minimize electrical interference across electrodes. Our default CI model (used in most experiments) used 500 pulses per second and a 200 µs offset between the corresponding pulses in two consecutive electrodes (i.e., the time difference between the kth pulse in the ith electrode and the kth pulse in the (i+1)th electrode). For each stimulus, the starting phase of each electrode’s pulse train was jittered by sampling the time of the initial pulse from the uniform distribution [0, inter-pulse interval] (the inter-pulse interval was the reciprocal of the pulse rate). We used monophasic pulses. Commercial cochlear implants mostly use biphasic pulses (with positive current followed by negative current, or vice versa), but the effects of such pulses were beyond the scope of our model. Biphasic pulses are used in actual implants because they are charge-balanced, such that no residual charge is transferred to the surrounding tissue, which is believed to help maintain the health of the tissue. The two phases of such pulses are believed to stimulate different locations on a nerve fiber^23^. However, because our model of electrical stimulation effectively had only one site of nerve stimulation, positive and negative current phases would simply cancel each other out, unlike what happens in the ear. We thus used monophasic pulses on the assumption that they best replicate the effect of a current pulse on the nerve given the constraints of our modeling approach. Each electrode’s pulse train was modulated with the corresponding envelope by point-wise multiplication. The resulting electrodogram was of shape M x T, where T is the number of time samples (and had the same value as in the normal hearing cochlear model).

To simulate the activation of auditory nerve fibers by electrical stimulation, we implemented an electrode-nerve interface that simulated the effects of spatial spread of the electrical field, temporal imprecision of auditory nerve responses, and the limited dynamic range of auditory nerve responses to electrical stimulation. The electrical stimulation from a CI creates an electrical field that spreads within the cochlea and excites nerve fibers within a spatial neighborhood, with the electrical field and resulting nerve activation dissipating away from the electrode. As in previous modeling work, we simulated this spatial spread of excitation using a double-sided, symmetrical, one-dimensional exponential decay function^45^. We simulated an unwound cochlea of length 35 mm with the M electrodes placed equidistantly from 8.125 mm (more apical) to 23.875 mm (more basal). The simulated electrical field at each nerve fiber location from an individual electrode was set as follows:

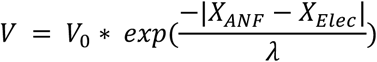

Where 𝑉_0_ denotes the maximum electric field (set to 1), 𝑋_*ANF*_ and 𝑋_*Elec*_ denote the locations of the auditory nerve fiber and electrode, respectively, and 𝜆 (in mm) denotes the spread constant. 𝜆 can be varied to simulate different extents of spread (as might depend on the position of the electrode within the scala tympani). We summed the electrical field from each of the M electrodes at each of the 50 locations corresponding to each auditory nerve fiber channel, resulting in a 50 x T representation of the instantaneous voltage across the cochlea.

Auditory nerve response to electrical as well as acoustic stimulation comprises many complex behaviors such as refractoriness, adaptation at different time scales, etc^47^. Many previous modeling approaches, both biophysical as well as phenomenological, have tried to capture these temporal dynamics using detailed implementations^49–52,132–134^. However, those implementations are often computationally expensive and time-consuming. We used a simpler alternative to enable the modeling of real-world behavior.

We simulated the auditory nerve response to the summed electrical field using a linear time-invariant system. The impulse response of this system was defined by a weighted sum of two Gaussians: an excitatory component (µ = 0.6 ms, 𝜎 = 0.6 ms, weight = 1.0) and a refractory component (µ = 2.2 ms, 𝜎 = 0.33 ms, weight=0.1), normalized to have a peak value of 1. The impulse response duration was 10 ms, intended to match the time scale of the nerve fiber’s temporal dynamics^97,135^. The nerve impulse response was convolved with the voltage time series at its location to obtain the input to the nerve rate-level function at that location. This linear model of the nerve’s response neglects known complexities of the nerve response, in particular refractoriness and adaptation^47,136^. We chose to neglect these phenomena for the sake of computational efficiency (models that implement these phenomena ^49–52,132–134^ cannot currently be implemented without being prohibitively slow; they are also not differentiable, which could be important in future applications of the model).

Electrical stimulation causes several differences in nerve fiber responses relative to what occurs with acoustic stimulation, and we sought to model these differences as they have been proposed to limit the information conveyed by a CI. First, because electrical stimulation bypasses the nerve fiber’s synapse with its hair cell, all nerve fibers, irrespective of their type (i.e., low, medium, high spontaneous rate), exhibit a narrow (∼6 dB each, ∼15 dB collectively) dynamic range. Second, a fiber’s spiking response becomes highly deterministic (again because the stochastic process of neurotransmitter release at the hair cell-auditory nerve synapse is bypassed). To replicate these effects, we gave all nerve fibers the same rate-level response function, and set the spontaneous rate to a near-zero value. The dynamic range was set to 15 dB, the spontaneous firing rate was set to 0.1 spikes per second, and the threshold was set to -5 dB re 1 mA current level. All fibers were given a maximum firing rate of 250 spikes per second.

The resulting CI-stimulated nerve firing rate representation was a three-dimensional tensor of shape N x T x C where N is the number of frequency channels (N=50), T is the total number of time samples (T=20000, with stimuli of length 2s and a cochlear sampling rate of 10,000 Hz) and C is the number of nerve fiber types (C=3). We sampled spikes from this firing rate representation using the same method as was used for the normal cochlea simulation described above.

### Current level settings using variance maximization

The CI model front-end contained free parameters for the threshold (TL) and maximum comfortable (MCL) current levels. Appropriate current settings were critical to ensuring that the CI-stimulated nerve responses carried enough information to perform downstream auditory tasks. For instance, if the current is overall too high or too low, stimulation could result in uniform nervegrams carrying little information about the stimulus. When humans receive a CI, the current settings (threshold and maximum comfortable level) are set by a clinician based on behavioral loudness judgments of the CI user, with the threshold level often set to be a fixed fraction of the maximum comfortable level. At present we do not know how behavioral judgments of loudness relate to the auditory nerve response, and as a result, it is not possible to perform the same procedure in a model. Instead, we developed an approach based on maximizing the variance in the nervegram to compute threshold and maximum comfortable level current settings that would avoid saturation of the nerve’s response.

The goal of our procedure was to maximize variance across stimuli in each of the three dimensions (N, T, C), where N = the number of frequency channels, T = the number of time samples, and C = the number of nerve fiber types. The rationale was that this would increase the distinguishability of the nervegrams of two different stimuli compared to alternative current settings.

We first initialized the maximum comfortable level (MCL) values with minimum and maximum initialization of -60 dB re 1mA and +40 dB re 1mA, respectively, in steps of 5 dB. In clinical settings, the dynamic range between the TL and MCL current levels is often set to a default value, with the clinician setting only the MCL. For instance, in Advanced Bionics cochlear implants, the TL is typically set to 1/10^th^ of the MCL^55^). To ensure that our settings were faithful to this constraint^85^, we hardcoded our TL values to be 20 dB below the MCL values (i.e., TL=MCL -20). This constraint also reduced the search space for the variance maximization algorithm. In each iteration of the algorithm, we sampled a combination of TL and MCL values from this range and computed the variance over the nervegram for each of a small subset of the evaluation dataset, containing 100 randomly sampled speech-in-noise examples at SNRs between -24 dB and +18 dB as well as clean speech. We calculated variance across all 100 stimuli resulting in a three-dimensional variance tensor from which we computed the mean of the variance tensor across all dimensions. Specifically, we took a tensor of shape [BxNxTxC], where B=100 is the number of stimuli, N=50 is the number of frequency channels, T=20000 is the number of time samples, and C=3 is the number of nerve fiber types. Then, for each of the NxTxC bins, we calculated the variance across the 100 stimuli, resulting in a tensor of shape NxTxC. We then computed the mean across all three dimensions of this tensor. We used the TL and MCL combination that resulted in the maximum value of this variance as the current settings.

### Intensity discrimination experiment

To validate our choice of the threshold and maximum comfort current levels arrived through the variance maximization procedure, we simulated an intensity change detection experiment similar to that described in a previous publication^57^. In the original experiment, experienced CI users listened to stimuli presented through pulse trains on a single electrode. Stimuli either had a constant intensity, or a momentary increase in intensity followed by a decrease back to the original level. Their intensity discrimination threshold, represented as the Weber fraction, was measured using an adaptive 2-down, 1-up, three-interval forced choice (3IFC) procedure that estimated the intensity change corresponding to 70.7% correct response in the psychometric function. The intensity discrimination threshold was analyzed as a function of the pedestal level (i.e., the baseline intensity to which the intensity change was added), represented in a percentage current dynamic range (% DR).

We started by sampling a range of current settings and computed the nervegram variance for each setting. We chose current settings that resulted in variance values greater than or equal to 60% of the maximum variance. For each chosen current setting, we then calculated the intensity discrimination threshold.

To simulate a discrimination judgment, we used a linear binary classifier composed of a 128-unit dense layer followed by dropout regularization and sigmoid activation. The classifier received the flattened CI nervegram as input and judged if the stimuli contained a change in the intensity (label 0: no change in intensity; label 1: change in intensity). We trained the linear classifier with a binary cross-entropy loss until asymptotic performance was reached and then evaluated the model for a range of pedestal levels and intensity changes.

In both the training and evaluation phases, we employed continuous pedestal stimuli with a total duration of 1000 ms. Stimulus with class label “1” consisted of three temporal segments: an initial 250 ms baseline period presenting the signal at pedestal intensity, followed by a 500 ms period presenting the signal at the altered (pedestal + intensity change) intensity, and concluding with a 250 ms return to baseline pedestal intensity. Stimuli with class label “0” were 1000 ms long sound stimuli maintained at a constant intensity. We used the baseline CIS simulator to generate the pulsetrains from the input sound stimuli and kept all CI parameters except the current settings unchanged across conditions. At every iteration of the training, we first generated the audio stimuli and obtained the corresponding nervegram which was then passed to the linear model. Linear model weights were updated via gradient descent using the optimization parameters as for training the neural network models. Stimuli were generated on the fly by sampling a pedestal value uniformly from the range (20 dB SPL, 80 dB SPL) and frequency value from the range (80 Hz, 8 kHz). The models were trained and validated on intensity changes ranging from (0.02 dBSPL, 12.38 dBSPL). All stimuli were sampled at 20 kHz. We evaluated the trained models on another set of stimuli that spanned the same pedestal level as training data but only 1 kHz frequency. The intensity variation in the evaluation data ranged from 0.01 dB SPL to 14.75 dB SPL.

For each current setting model, we generated the psychometric function by plotting the detection accuracy (%) as a function of the intensity change value (ΔI). For each pedestal condition (I), we obtained the intensity threshold by calculating the ΔI required to achieve 70.7% detection accuracy. The intensity discrimination limen, denoted by the Weber function, was given by 10log(ΔI/I). Note that both I and ΔI were first converted from the dB SPL value to the current value using the loudness growth function for the specific current settings. Similar to the original publication, we calculated %DR as

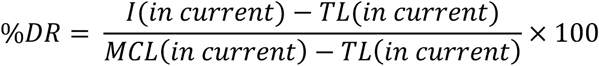

### Deep neural network architectures

The simulated auditory nerve response from either the normal cochlea or the CI was passed through a deep artificial neural network. The neural network consisted of a series of hierarchical feed-forward stages. Each stage was comprised of four operations: (1) 2-dimensional convolution using linear kernels, (2) point-wise rectification using a rectified linear unit activation function, (3) pooling, and (4) layer normalization. The very last stage of the neural network was a fully connected layer, mapping the 3-dimensional output from the convolution stage to a vector. Classification was performed on this vector using the softmax operation. Softmax converts the model output representations into a probability distribution over the output class labels such that the class label attaining the highest probability is interpreted as the model prediction for a given stimulus. We also included a dropout layer in the last stage (immediately following the fully connected layer, before the classification heads) that helped regularize the model and avoid overfitting. The dropout rate was 0.5, and dropout was used only during training (not during evaluation).

For each task (word recognition and sound localization), we trained 3 unique neural network architectures. These architectures included the same set of operations described above but had different hyperparameters (e.g., number of stages, size and shape of the convolution kernels, the extent of pooling, size of the fully connected layer, etc.). The architectures were identified through a large-scale architecture search^17,19^ to find architectures that performed one of the tasks well. The behavior of the different architectures was analyzed analogously to that of individual human participants. For practical reasons, in three of the experiments, only one of the three architectures was used, as described below. Behavioral results were generally similar across the three architectures. The architectures of the word recognition and sound localization tasks differed in the number of stages, and the pooling operation (Hanning-weighted average pooling^64^ for speech task vs. max pooling for localization task) used.

The speech models operated on monaural input, with input to the neural network from an auditory nerve representation of shape [N frequencies x T time samples x C nerve fibers] obtained from a single channel audio. By contrast, the localization models operated on binaural input. The audio signals for the left and right were separately processed by either the cochlear-to-nerve or the CI-to-nerve simulation, each producing an auditory nerve representation of shape [N frequencies x T time samples x C nerve fibers]. These representations were then concatenated along the fiber dimension to obtain a [N frequencies x T time samples x 2C nerve fibers] representation that was the input to the neural network.

### Peripheral neural degeneration

To simulate peripheral neural degeneration, we eliminated fibers at the spike sampling stage by limiting the maximum number of spikes that could be sampled at each time-frequency-fiber bin in the nerve representation. In the no degeneration conditions, we simulated a maximum of 640 fibers per frequency channel (32000/50). To simulate 90% peripheral degeneration, for example, we reduced the number of maximum spikes to 64 (10% of 640). We imposed uniform degeneration across all frequencies and nerve fiber types.

### Central neural degeneration

We simulated neural degeneration in the central auditory pathway by silencing a group of randomly chosen frequency-channel combinations at each stage of the deep neural network. The output from each stage of the neural network was a 3-dimensional tensor with shape [N frequency bins, T timesteps, C channels] whose dimensionality that varied across stages. For instance, to simulate 50% central degeneration, we randomly chose 50% of the N x C units (channels within each frequency bin) at each stage’s output and silenced the responses for all T timesteps. The silenced activations were included as part of the input to the next stage of the model. The selection of units to be silenced at each stage was performed prior to any retraining of the CI model, and remained unchanged during retraining.

### Training and validation dataset – word recognition in noise

The neural network models of word recognition were trained on all three recognition tasks for the Word-Speaker-Noise dataset (i.e., word recognition, speaker recognition, and environmental sound recognition). This dataset ^20,62,63^ was inspired by the task originally used by Kell et al.^17^ and is detailed in previous papers from our lab^20,63,64^. We trained models on all three tasks despite only evaluating the model on word recognition because we found empirically that this aided optimization. The description of the dataset here is adapted from the original paper^64^.

Speech segments were obtained from two large-scale speech recognition corpora: Wall Street Journal (WSJ; speakers reading excerpts from Wall Street Journal) and Spoken Wikipedia (SWC; speakers reading articles from Wikipedia). Feather et al. counted the occurrences of different words in these corpora and only included words at least four characters long. They then sampled two second-long speech segments containing the chosen words overlapping at the mid-point of the sentence excerpt. The ground truth class label for each sentence excerpt corresponded to the target middle word.

To generate a balanced dataset, speaker and word class labels were chosen such that each word class label contained approximately 50 unique speakers and vice versa. Class labels were additionally constrained such that there were at least 200 unique utterances within each word and speaker class. Also, to ensure diversity of stimuli even within a single class label, a maximum of 25% of the stimuli were allowed to come from a single cross-class label (e.g., for a given word label, at most 25% of the stimuli were spoken by the same speaker). The resulting stimuli were then sub-sampled so that there were at most 2000 stimuli for each word or speaker class. This process resulted in a training dataset containing a total of 230356 clean speech segments covering 793 unique word classes and 472 unique speaker classes. A similar process was performed to generate the validation dataset containing a total of 40650 speech segments.

During model training, each training example was generated by randomly sampling a foreground speech segment from the above-described dataset and combining it with a randomly selected background noise sample. The background noises were extracted from AudioSet, a large-scale dataset of a wide variety of sound events^137^. To avoid ambiguity with the foreground signal, AudioSet sounds tagged as “speech”, “whisper” or “silence” were excluded. To help ensure that the noise was present throughout a training example, sounds that had more than 10% zeros were excluded. Additionally, AudioSet contains a large proportion of “music” stimuli. Therefore, to maintain class balance, clips labeled only “music” without any additional branch labels were also excluded. This resulted in a background noise dataset of 718625 unique clips covering 516 sound categories.

Speech and noise excerpts were combined at a signal-to-noise ratio (SNR) uniformly sampled from -10 dB to 10 dB. 9% of training examples did not have background noise (i.e., SNR = infinite). Each stimulus was rescaled to a dB SPL value sampled uniformly from the range of 30 to 90. This random sampling of foreground and background stimuli pair as well as the SNR and sound level results in an arbitrarily large dataset that was generated in batches on the fly during the model training. The validation data were also generated on the fly via the same process.

### Evaluation dataset – word recognition in noise

To evaluate the model performance following training, we used an evaluation dataset containing ∼64,000 stimuli generated using a procedure similar to that used for the training/validation dataset. The key difference in this dataset was that we used five specific types of background noise rather than sampling randomly from a set of natural backgrounds. There were two main rationales for this choice. The first was that evaluating the models on conditions on which they were not directly trained allowed us to get a sense of generalization ability. The second was that it allowed us to compare the performance on different noise types to that of humans. The five noise types were: (1) Auditory scenes from the IEEE AASP CASA Challenge dataset (2) 8-talker speech babble from CommonVoice (3) instrumental music from MUSDB18 (4) Speech shaped noise and (5) amplitude-modulated speech-shaped noise. These noise types were the same as those in the speech-in-noise experiment in Kell et al^17^. To generate the speech-shaped noise, we randomly chose clean speech samples from the evaluation data, computed the speech spectrum, and imposed this spectrum on randomly generated samples of white noise. To generate the amplitude-modulated noise makers, we first generated the speech-shaped noise carrier. Then we randomly chose another clean speech sample from the evaluation data, extracted its envelope (by rectifying and lowpass filtering the waveform, using a cutoff of 40 Hz) and imposed this envelope on the speech-shaped noise carrier^138^. Models were tested on stimuli with SNRs varying from -24 dB to +18 dB in steps of 3 dB, as well as clean speech stimuli. Audio presentation level was fixed at 60 dB SPL.

### Training and validation dataset – sound localization

Training data were the same as those used in a previous publication from our lab^20^, and this section is reproduced, with minor changes, from that paper. Sound localization models were trained on binaural audio signals: spatialized “foreground” natural sounds embedded in natural noise recordings. The foreground natural sounds were obtained from the Glasgow Isolated Sound Events (GISE-51)^139^ dataset subset of the Free Sound Dataset 50k (FSD50k)^140^. This dataset contained variable-length sounds covering 51 categories out of which we selected those having a 44.1 kHz original sampling rate (in order to support the frequencies needed for elevation cues). This filtering procedure resulted in 12465 training and 1716 validation source clips.

Texture-like background noises were obtained from the AudioSet dataset by excluding non-stationary stimuli such as speech and music. We defined stationarity by measuring the texture statistics (e.g., envelope means, correlations, and modulation power in and across cochlear frequency channels) and quantifying the stability of such statistics^141,142^. We excluded sounds whose statistical variability (denoted by the standard deviation of the texture statistics parameter) was greater than 0.5. This stationarity screening resulted in excluding 11.9% of the original AudioSet from being used for model training/validation. This procedure resulted in 26515 and 562 training and validation noise clips, respectively. To increase the variability of the training data, bandpass filters were applied to 50% of the source stimuli. The center frequencies of the filters were sampled log-uniformly between 160 Hz and 16 kHz. Bandwidths and filter orders were log-uniformly sampled between 2 and 4 octaves and 1 and 4, respectively.

To generate spatialized audio, both target and noise stimuli were passed through binaural room impulse responses generated using a virtual acoustic room simulator^143^. The simulator used the image-source method to simulate room impulse responses and include KEMAR HRTFs. We used these HRTFs for both the normal-hearing and cochlear-implant models. We note that the pinna cues in these HRTFs are not available to human cochlear implant users because the device microphone is located outside the ear canal. We used these HRTFs for lack of a better option, reasoning that the pinna cues would not be preserved by the cochlear implant stimulation and thus would be unlikely to influence the results.

The simulator outputs two impulse responses which when convolved with a source signal reproduce the sound that would arrive at the ears of a listener at a specified position in a room with the source signal at a specified location in the room. Using this simulator, we generated binaural room impulse responses for a set of listener and source positions for each of 2000 unique shoe-box-shaped rooms with height varying between 2.2 and 10m and length and width varying between 3 and 30m. The listener head positions (x, y, z coordinates) and head angles in each room were sampled uniformly subject to the constraint that the head was positioned no higher than 2m from the floor and at least 1.45 m away from each wall. This enabled us to synthesize binaural signals that spatially rendered the foreground sound at one position in a room with noise signals presented at a set of other locations.

We rendered sounds over an azimuth range of 0 to 360 degrees in steps of 5 degrees (72 unique azimuth locations) and an elevation range of 0 to 60 degrees in steps of 10 degrees (7 unique elevation locations), resulting in a total of 504 (=72×7) unique azimuth-elevation positions. The model performed a classification task over this set of positions. For each training stimulus, a target was rendered at one of the 504 locations, chosen randomly. 3 to 12 different noise stimuli were rendered at 3 to 12 different locations, chosen randomly. For the training dataset, the room was chosen randomly from a set of 1800 rooms subsampled from the original set of 2000. The remaining 200 rooms were used for the validation dataset. After spatialization, the target and noise stimuli were added at SNRs uniformly drawn between -15 and +25 dB, except for 5% of scenes that included no noise. Overall, this process resulted in 1,814,400 and 201,600 auditory scenes in the training and validation dataset, respectively. Each scene was rescaled to a sound pressure level sampled uniformly from the range of 30 to 90 dB SPL.

### Model optimization

We trained the word recognition model jointly on three different tasks: word recognition, speaker recognition, and background sound recognition. Although we only analyzed model behavior for the word recognition task, this multi-task training paradigm aided optimization and led to faster convergence of the CI models, specifically the ones with the HiRES120 CI sound coding strategy. For multitask training, we used three cross-entropy loss values for the three tasks, and the final loss was defined by the weighted sum of the three losses (with weights 1.0, 0.25, and 300.0 for the word, speaker, and background sound recognition tasks respectively; these weights were determined empirically as values that caused all three losses to decrease). We used the sparse categorical cross-entropy loss function for all tasks, for both the word recognition and sound localization models.

Model parameters were initialized either using the predefined distributions in Tensorflow (for the normal hearing model and CI model with random initialization) or learned weights from the normal hearing model (for all other CI models). Parameters were learned iteratively using stochastic gradient descent. We used the ADAM optimizer with 𝛽_1_ = 0.9, 𝛽_2_ = 0.999, 𝜖 = 10^−7^, and a batch size of 32. A learning rate of 1e-5 was used for all models except for those that used the HiRES120 CI strategy (for which this learning rate did not reliably cause the loss to decrease). For HiRES120 models, a local hyperparameter search was performed by varying the learning rate between 1e-4 and 1e-7 in factors of 10. This resulted in a final learning rate choice of 1e-6 for that sub-group of models.

Each model was trained until its validation accuracy reached an asymptote. Validation set performance and training set loss were evaluated after every 10,000 training steps for the word recognition task and after every 5,000 steps for the sound localization task. The number of training steps needed to achieve asymptotic performance varied depending on the tasks, model architecture, and conditions (plasticity, neural degeneration, different initializations, etc.). Learned parameters were saved as checkpoints, with the checkpoint producing the best validation performance used for model evaluation. All model training was performed on NVIDIA A100 GPUs available through the MIT OpenMind Computing cluster. Models took, on average, around 5 days of training on a single GPU.

### Model evaluation

Once training was completed, each of the architectures was tested using the evaluation dataset for the model’s task. To obtain the model prediction, each test stimulus was first passed through the nervegram generator (for either the normal cochlea or CI) and then through the learned neural network. The model yielded a probability distribution over all the possible class labels. The class label with the highest probability was taken as the model response. The model prediction was compared to the ground truth label for each stimulus within a condition to obtain either word recognition accuracy (proportion correct) or localization error (the mean absolute angular difference between the model predicted location and true location). The mean response for an experimental condition was obtained by averaging performances across the 10 architectures for a task. During evaluation, all model parameters were frozen.

### Human cochlear implant user word recognition experiment

To evaluate how well our CI models replicate the behavior of human CI users, we measured CI users’ word recognition performance on the same task the models were tested on.

#### Informed consent

All participants provided informed consent. The Massachusetts Institute of Technology Committee on the Use of Humans as Experimental Subjects (COUHES) approved all experiments.

#### Participants

We recruited four (three female) postlingually deafened adult CI users in the age range of 61 to 80 years (mean = 69.5 years, S.D. = 7.85 years) through the Audiology Clinic of Massachusetts Eye and Ear Infirmary. All participants had normal hearing for a significant part of their life and then were implanted with a CI following deafness (see Table 1). All had lived with their CIs for at least a few years prior to the experiments and used spoken language to communicate. Three participants provided their Consonant-Nucleus-Consonant (CNC) word recognition scores from previous clinical assessments; these scores are provided as percentiles in Fig. 3B.

**Table 1.** Details of the CI participants who completed the word recognition task

| Patient ID | Age as of 2024 (Gender) | Etiology | Hearing status |  | CI device | Implantation type | CI Experience (years) |  | Stimulus presentation level (RMS amplitude) |
| --- | --- | --- | --- | --- | --- | --- | --- | --- | --- |
|  |  |  | Left | Right |  |  | Left | Right |  |
| 003 | 61 (female) | Hereditary | Hearing loss, aided with CI | Hearing loss, aided with hearing aid | Advanced Bionics Marvel | Unilateral | 10 | NA | 0.2000 |
| 005 | 80 (male) | Unknown | Hearing loss, aided with CI | Hearing loss, aided with CI | Advanced Bionics Marvel | Bilateral (asynchronous implantation) | 6 | 2 | 0.1100 |
| 006 | 68 (female) | Unknown | Hearing loss, aided with CI | Hearing loss, aided with CI | Advanced Bionics Marvel | Bilateral (asynchronous implantation) | 10 | 2 | 0.0356 |
| 008 | 69 (female) | Meniere's disease | Hearing loss, aided with CI | Hearing loss, aided with hearing aid | Advanced Bionics Marvel | Unilateral | 8 | NA | 0.0356 |

#### Procedure

On each trial, participants listened to a 2s long speech segment embedded in noise and reported the word in the middle of the speech segment (i.e., that overlapped the 1s mark). Stimuli were presented directly to the participant’s cochlear implant from the experiment computer via Bluetooth. Individual subjects’ implant settings were kept unchanged except that all noise reduction algorithms were disabled for a fair comparison with the models.

The target words were drawn from the model vocabulary (a set of 793 words). Participants were familiarized with the word set before the start of the experiment, by reading a list of the words. After each stimulus, participants typed their response into an interface that autocompleted with the words in the experiment vocabulary that were consistent with the entered text. Responses could only be entered if the entered text string matched a word in the vocabulary. In this way the participants performed the same task as the models, which were also constrained to only report words within the vocabulary. Sound was delivered exclusively via Bluetooth. For unilaterally implanted participants, this meant that their other ear could not contribute to task performance. For participants who were bilaterally implanted, half of the experiment was performed using their left ear and the remaining half using their right ear, and results for the two ears were analyzed together. Each participant completed a 2-hour listening session.

Participants were tested in 4 SNR conditions: +6 dB, +12 dB, +18 dB and clean. There were five different background noise types (auditory scene, instrumental music, 8-speaker babble, speech-shaped noise, and amplitude-modulated speech-shaped noise), resulting in 16 total conditions (3 SNRs x 5 background noise types + clean speech). The experiment contained twice as many clean speech trials as one of the noise conditions. The SNRs were selected based on pilot experiments to produce a range of performance. There were 18 unique stimuli within each condition (36 for clean speech), thus resulting in a total of 306 stimuli. Each stimulus contained a different word. Each participant heard the same set of 306 stimuli. The experiment was divided into 6 blocks, each consisting of 51 trials. Before the main experiment, participants completed a set of practice trials (20 trials with clean stimuli and 15 trials with speech in noise, one for each of the 3 SNRs x 5 background noise type combinations) to familiarize themselves with the task. Participants received feedback during the practice trials but not the main experiment.

Before the start of the experiment, we calibrated the level of the stimuli for each participant individually. To do so, we played a sample stimulus with an RMS amplitude of 0.07 with the computer volume setting at the maximum level. We then increased the level in steps of 5 dB while receiving verbal feedback from the participants as to whether the stimulus was audible and whether it was uncomfortably loud. We chose a sound level that was audible as well as comfortable for the individual participant and presented all stimuli for the rest of the experiment at this pre-identified sound level. We did not make any changes in the settings of the participant’s device.

All participants used the Advanced Bionics Marvel CI device. This CI has built-in algorithms for performing noise cancellation that were by default enabled in the participants’ devices. To enable human-model comparisons on a level playing field, we manually disabled all noise cancellation in the participants’ CI using the TargetCI software provided by Advanced Bionics. All other device settings remained unchanged.

### Normal hearing human listener word recognition experiment

The normal hearing listeners’ word recognition data were taken from a previous publication^20^. Subjects performed the same task as the cochlear implant users, however through an online experiment platform. Extensive research conducted in our laboratory has consistently demonstrated that online data can match the quality of data collected in traditional laboratory settings^17,144–149^ provided measures are taken to ensure standardized sound presentation, encourage participant compliance, and promote task engagement. Online participants were recruited using the Prolific platform. To ensure data quality, participants were prescreened to have at least a 95% submission approval rate, Participants again typed responses into a textbox and, as they typed, the displayed list of 793 words was filtered to include only words that matched the entered string. Only responses from the word list could be submitted. Recognition data were collected for 6 SNR conditions (−9, -6, -3, 0, +3 dB). Individual participants heard only 376 stimuli (each unique speech excerpt was presented once), uniformly sampled across SNRs and noise types. All 44 subjects (30 male) self-reported as normal hearing individuals, passed a headphone check^144^, completed at least 100 trials and responded correctly to the catch trials at least 85% of the times. Participants heard a calibration sound and adjusted the presentation level to be comfortable prior to the start of the experiment. Participants ages were between 18 and 62 (median 36) years.

### Evaluation dataset – noise-vocoded speech

To generate noise-vocoded speech, the speech signal was first converted into a set of narrow subbands using a bank of 320 filters. Filters were zero phase and were the positive portion of a single period of a cosine function, with center frequencies ranging from 20 Hz to 10 kHz. Adjacent filters overlapped by 50% so that the filters perfectly tiled the frequency spectrum. Filters were evenly spaced on an Equivalent Rectangular Bandwidth (ERB) scale^125,150^. These subbands were summed in various combinations to produce the desired number of channels (i.e., 1, 4, 16, and 64 bands). For instance, to generate a single band, all of the subbands were summed, whereas to generate 4 bands adjacent sets of 80 filters were summed. The envelopes of these summed subbands were extracted as the magnitude of the analytic signal (obtained via the Hilbert transform). These envelopes were imposed on the subbands of a Gaussian noise signal (obtained via the same filtering and summing operations used to obtain the speech signals) by multiplication with the noise subbands, after dividing the noise subbands by their original envelope. The resulting subbands were refiltered with a filterbank of either 1, 4, 16, or 64 filters (obtained by summing together subsets of the transfer functions of the filters used to obtain the original set of narrow subbands) and summed to generate the noise-vocoded speech signal.

### Normal hearing human data – noise-vocoded speech

Human speech intelligibility with noise vocoded speech was measured as part of a larger human psychophysics study performed in the lab^150^. 14 participants (4 female, mean age: 28 years, range: 22-33 years) from this study performed the noise-vocoded word recognition experiment. As in the other human experiments described above, participants listened to 2s long stimuli and typed their response in a text box. Sounds were presented using the Psychtoolbox for MATLAB at 70 dB SPL with a sampling rate of 16 kHz and were presented via the soundcard on a MacMini and Behringer HA400 amplifier through Sennheiser HD280 headphones. All experiments were conducted in a soundproof booth and were approved by the Massachusetts Institute of Technology Committee on the Use of Humans as Experimental Subjects and all participants provided informed consent.

### Simulating the effect of number of electrodes (Fig. 3C)

We analyzed the dependence of CI model word recognition on the number of active electrodes by simulating an experiment similar to one previous run with human CI users^67^. In this previously published experiment, 11 post-lingually deafened adult cochlear implant users (>= 6 months experience) performed word recognition on clean speech recordings. Each participant listened to speech stimuli using 4, 8, 10, 16, and “all” electrodes. In the “all” condition, all electrodes in the device were active; the number varied from 19 to 22 depending on the participant. In all conditions the electrodes were mapped to a constant frequency range of 188 to 7938 Hz. All other device settings were unmodified from their clinical settings. Participants did not receive training with the altered number of electrodes. Stimuli were presented at 60 dB SPL in a single-walled sound booth over a loudspeaker located at 0-degree azimuth and 1m distance from the participant. The test stimuli included Consonant Nucleus Consonant (CNC) words, and AzBio sentences in quiet and at + 5 dB SNR. We used only the results from the AzBio sentence recognition task in quiet. We scanned human results from the original paper (black bar graphs in Figure 1A in the paper by Berg and colleagues) using WebPlotDigitizer.

We simulated a similar experiment in a model of cochlear-implant-mediated word recognition. The model was optimized for nervegrams generated with 22 simulated electrodes using the CIS sound coding strategy (using normal-hearing weight initialization, and either re-optimizing all or a subset of model stages), and then were evaluated on simulated nerve representations from 1, 2, 4, 8, and 16 active electrodes. This scenario was intended to replicate that of the human participant who normally used the full set of device electrodes but then was tested in the lab using fewer electrodes. The models performed the middle word recognition task on the evaluation dataset described above (as a result, the models were not tested on the same stimuli, or with exactly the same task, as the humans to which they are compared). We used only a single model architecture (arch0_0000) in this experiment. Because the human participants and models were not tested on the same stimuli and task, the results should not be expected to be quantitatively consistent. The human data were reported in the “Rationalized Arcsine Unit” (RAU), whereas our model evaluation quantified performance with proportion correct.

### Noise burst localization experiment

We replicated a localization experiment previously conducted with cochlear implant users as participants. In this experiment, normal hearing listeners^65^ and cochlear implant users^66^ listened to noise bursts containing frequencies between 300 Hz and 10 kHz, presented in an anechoic room. On each trial, a noise burst was played from a randomly chosen speaker location (at 0, ±15, ±30, ±45, ±60, or ± 75 degrees for the normal hearing listeners and at ±10, ±30, ±50, or ± 70 degrees for cochlear implant users). Participants identified the source location of the target sound. We ran our localization models on a simulation of this experiment, using the same noise burst stimuli virtually rendered in a simulated anechoic room.

The localization models predicted the azimuthal location of the noise burst rendered in the simulated environment detailed above. The models were originally trained to predict azimuth locations at 5 degrees spacing. To simulate the experiment (in which locations were drawn from a smaller set), we performed post-processing on the model location judgements. Specifically, the model judgment was taken to be the experimental location closest to the model’s prediction. If the model prediction was equidistant between two experimental locations, then the judgment was assigned equiprobably to one of the two locations.

### Simulating variation in the extent of central plasticity

We simulated different extents of plasticity by varying the number of stages in the neural network that were reoptimized for CI-stimulated nerve representations. For the main plasticity manipulation (Fig. 8), we first initialized the neural network using the parameters learned with normal cochlea input. This initialization could be conceptualized as analogous to the effects of an initial period of auditory experience prior to the onset of deafness. Then during retraining with CI-stimulated nerve responses as input, we specified which stages of the network would be further updated. During the gradient-based weight update process, only the parameters of the specified “plastic” stages were modified. All learnable parameters of all operations in the specified stages were modified. This implementation of plasticity assumes that plasticity is all-or-none within an individual stage. We also assumed hierarchical continuity of plasticity, meaning if one model stage as plastic then all deeper stages were also plastic. For example, in the plasticity condition “stages conv4 and beyond”, the conv4 stage and all model stages beyond that (e.g., conv5, conv6, etc.) were reoptimized for CI input.

### Normal hearing vs random weight initialization

As an alternative to initializing the neural network with normal hearing weights and then retraining those weights with CI input, we also analyzed the effects of random weight initialization. In this case each learnable parameter of the network was initialized with a random value drawn from a predefined distribution (default settings in Tensorflow; for example, glorot uniform distribution to initialize convolution kernels and zeros for bias terms, ones and zeros for the scaling and bias parameters in layer normalization, etc.). Then, the neural network parameters for all or a subset of model stages were updated during training with CI input, as described in the previous section.

### Simulation of different sound coding strategies

In addition to the continuous interleaved sampling strategy that we used as a baseline, we implemented and compared the strategies used by the three main current CI manufacturers. Our implementations were based only on publicly available information (published descriptions and code from one manufacturer).

#### Advanced Combination Encoder

Advanced Combination Encoder (ACE) is a CI sound coding strategy widely used in CIs manufactured by Cochlear Ltd. ACE combines the CIS strategy with N-of-M, another sound coding strategy used in early CIs that dynamically selected a subset of electrodes to stimulate at each point in time. ACE aims to combine the temporal resolution benefits of CIS with the spectral resolution benefits from N-of-M. In ACE, audio is analyzed in frames, with electrodes selected dynamically based on the peaks in the sound amplitude in different frequency bands (similar to N-of-M). The electrodes selected in each frame are activated in an interleaved fashion (similar to CIS). We simulated a 16-electrode ACE model in which only 8 electrodes were active at a time. The initial CI processing used the subband envelope extraction and loudness growth function as used in the baseline CIS model, and then divided the envelopes into 2 ms frames. We chose the pulse rate to be 800 Hz. In each frame, we identified the 8 frequency bands whose envelopes had the highest mean amplitudes. The pulse trains for the corresponding 8 electrodes were amplitude-modulated using the corresponding frame of the compressed envelopes of the 8 selected bands, whereas the pulse trains of the other 8 electrodes were set to zero. The electrode signals for all audio frames were concatenated to yield an electrodogram of shape 16 x T.

#### Fine Structure Processing

Fine Structure Processing (FSP) is a sound coding strategy used in MED-EL CIs. Unlike CIS-based strategies that encode only envelope information, FSP strategies aim to encode the temporal fine structure of the sound, particularly in the low to mid-frequency range. FSP as implemented in current MED-EL CIs uses channel-specific sampling sequences at the most apical 2-4 electrodes. Instead of having a fixed pulse rate in those electrodes, the stimulation rate is continuously varied to capture the instantaneous fine structure information in the corresponding frequency band of the audio input. We implemented a version^40^ of the FSP strategy in which the 3 most apical electrodes encoded fine structure information while the remaining 13 electrodes were stimulated using envelope information as in CIS. Incoming waveforms were divided into M=16 subbands. The subbands for these remaining 13 electrodes were further processed in the same way as the baseline CIS, as described above. The time and amplitude of pulses in the 3 most apical electrodes were determined by the local temporal maxima within frames of the corresponding subbands^104^. We used a pulse rate of 800 Hz for the 13 most basal electrodes. For the 3 most apical electrodes, we used an analysis frame length of 1ms to compute the fine structure information (frames did not overlap). This again yielded an electrodogram of shape 16 x T.

#### High Resolution with Fidelity 120

High Resolution with Fidelity 120 (HiRES120) is a sound coding strategy developed by Advanced Bionics LLC. HiRES120 is an extension of an earlier strategy known as HiRES that was a particular instantiation of the CIS strategy that at the time offered a higher stimulation rate and more channels (e.g., 16) than had been previously available. Our implementation of CIS used for the baseline cochlear implant model is in fact the original HiRES strategy. HiRes120 augments HiRES with additional “virtual channels” by a mechanism known as current steering. The goal is to increase frequency resolution, enabling 120 different virtual channels. Current steering works by concurrently stimulating two adjacent electrodes in a weighted fashion such that the maxima of the resulting electric field occur somewhere in between the locations of the two electrodes. As a consequence, the peak of stimulation can be “steered” to points other than the electrode locations, potentially resulting in improved frequency resolution. HiRES120 uses 15 pairs of electrodes (i.e. all pairs of adjacent electrodes from the set of 16 electrodes, assuming all 16 are active). Current steering creates 8 additional stimulation sites between each electrode pair, totaling 120 virtual stimulation sites. As discussed in detail elsewhere^41^, HiRES120 uses an analysis filter for each electrode pair to determine the amplitude envelope and spectral peak to be conveyed by each pair of electrodes at each of successive time frames. The two electrodes in a pair emit synchronized pulses whose amplitudes are determined by the envelope, with current steering weights for the two electrodes to convey the spectral peak. All electrodes apart from the first and last emit two pulses per frame, one for each of the two pairs the electrode is part of. The pulses for different pairs are interleaved in time. This sound coding strategy also includes a carrier synthesis stage which modulates the Hilbert envelope of each analysis band with the spectral peak frequency for that band and is intended to represent the temporal structure in that analysis band. We used an open-source implementation of this strategy available from Advanced Bionics (https://github.com/jabeim/AB-Generic-Python-Toolbox). The only changes we made were to modify the pulses to be monophasic instead of biphasic. Also, since our other CI models did not include any explicit automatic gain control or noise cancellation, we disabled these in the HiRES120 simulation as well. This again yielded an electrodogram of shape 16 x T. We trained a single model architecture for this stimulation strategy (one for each task; arch0_0000 for word recognition and arch0 for sound localization) as the other two model architectures trained prohibitively slowly. Results for the other strategies were similar across architectures and so it seemed likely that a single model architecture would provide an accurate estimate of performance.

#### Current settings

The threshold and maximum comfort current levels were optimized separately for each strategy using the variance maximization approach described above to make sure that the resulting nervegram was not entirely saturated or inactive. The electrode-nerve interface in all cases remained the same to help ensure a fair comparison across strategies.

### Word recognition and sound localization with different sound coding strategies

To simulate model word recognition and sound localization performance with different sound coding strategies discussed above, we followed the same approach to our baseline CIS-based model except that the pulse rates were set to 800 pulses per second (to be faithful to their current standard implementations). For each strategy, we implemented a model stage to generate the CI-stimulated nervegram specific to that strategy. This model stage provided the input to the neural network model stages. Each of the architectures discussed previously was trained separately for each sound coding strategy and each task (word recognition or sound localization). The training, validation, and evaluation dataset remained unchanged across cases. The model training paradigm for simulating different extents of plasticity was identical to that for the baseline CIS model.

### Manipulation of spatial spread of electrical excitation

We manipulated the spatial spread of electrical excitation by varying the spread constant of the exponential decay function that governed spatial spread. We tested two scenarios intended to reproduce the current spread for monopolar and bipolar stimulation, respectively, that are known to differ in their spatial spread of excitation. The broad spatial spread (monopolar) condition was intended to reproduce current that decays by approximately 1.2 dB/mm, via a spread constant of 6. The medium (bipolar) and narrow (tripolar) spread conditions were intended to reproduce current that decays at approximately 3.5 dB/mm and 5 dB/mm, via a spread constant of 2.35 and 1.5, respectively.

For each spatial spread condition, we also tuned the threshold and maximum comfort current settings using the variance maximization algorithm so that the nervegram in each condition had the best chance of conveying information about sound stimuli. This process is analogous to how current levels are readjusted for human participants when the type of stimulation used in their device is changed.

### Word recognition and sound localization with different current level settings

To simulate model word recognition and sound localization performance with different current level settings, we first chose two groups of current levels: 1) settings that resulted in intensity discrimination thresholds similar to those of human cochlear implant users and 2) settings that resulted in substantially worse intensity discrimination threshold than those of cochlear implant users (Figure 2E). We then reoptimized a neural network using each current setting (for either the task of word recognition or sound localization). The two plasticity conditions were implemented as described previously. We only trained one architecture (arch0_0000 for word recognition and arch0 for sound localization) with alternative current settings.

### ITD/ILD cue weighting

This section is reproduced with minor changes from an earlier publication from our lab^20^ that used the same experiment on a different set of models. To analyze the CI model sensitivity to ITD and ILD cues, we simulated the localization cue weighting experiment by Macpherson and Middlebrooks (2002)^73^ on our model using the stimuli described in an earlier modeling paper^19^. In the original human experiment, 13 participants (5 female; age range 18-35 years) listened to virtually rendered stimuli played over headphones and reported the perceived azimuth by orienting their heads towards the estimated source locations. The stimuli they heard contained artificially imposed additional ITD and ILD cues, therefore allowing the authors to evaluate human response bias as a function of the imposed localization cues. The stimuli included two sets of noise bursts: high pass (4-16 kHz) and low pass (0.5-2 kHz). Each stimulus was of length 100 ms with 1 ms squared-cosine ramps at onset and offset.

Our normal hearing and CI localization models were presented with identical noise burst stimuli spatialized using a room simulator that simulated an anechoic room. All sounds were located at 0 degrees elevation and 0-360 degrees azimuth in steps of 5 degrees. We analyzed model behavior for imposed ITD biases of ±300 µs and ±600 µs time delays; and imposed ILD biases for ±10 dB and ±20 dB level differences between the two ears. Model responses were front-back folded to make it directly comparable to the human experimental setup (in which stimuli were only presented in front of participants).

To compute the ITD response bias, we subtracted the ITD values associated with the true azimuth of the stimuli (i.e, the azimuth location from where the stimuli were actually played) from the ITD values associated with the model-predicted azimuth (i.e, the azimuth location that, the model thinks, stimuli were played from). Similarly, the ILD response bias was calculated by subtracting the ILD values of the true azimuth from the ILD values associated with the model-predicted azimuths. In Figure 6D, we plot these response biases as a function of imposed biases (ITD and ILD) for two noise bands, resulting in a total of four conditions. If the model response was strongly dependent on the manipulated localization cue (ITD or ILD), the response bias would vary linearly with a slope close to 1.0 as a function of the imposed bias. Alternatively, a slope close to 0.0 would indicate the localization cue did not affect the localization response. The bias weights reported in Figure 6E are these slopes (derived from fitting lines to the data).

### Analysis of ITD/ILD electrodogram cues

We quantified the ITD/ILD cues encoded in the electrode signals from CIS, ACE, FSP, and HiRES-120 sound coding strategies by measuring ITDs and ILDs from electrode responses and comparing these to the ITDs and ILDs in the stimulus. We analyzed each electrode separately using narrow-band noise stimuli centered at the filter used for the electrode (with a bandwidth of 80 Hz). For the ITD cue analysis, we imposed ITDs of -1000µs to +1000 µs in steps of 100µs by delaying the stimulus in one ear with respect to the other ear. For the envelope ITD analysis, we first imposed amplitude modulation on the noise (40 Hz modulation rate, with a depth of 100%). However, the ITD was imposed on the entire waveform, such that the audio stimulus contained both envelope and fine structure ITDs. For the ILD cue analysis, we imposed ILDs of -10 dB to +10 dB in steps of 1 dB on the noise bands in the two ears. These synthetically generated binaural stimuli were then processed by the sound coding strategy under analysis to obtain a binaural electrodogram.

To compute the ITD cues encoded in the electrodogram fine structure, we cross-correlated the right and left ear electrode signals and identified the lag (in µs) corresponding to the maximum similarity in the two channels, restricting analysis to lags within +-1 ms. The resulting cross-correlation should be dominated by interaural differences in pulse timing. To compute the ITD cues encoded in the electrodogram envelopes, we lowpass filtered the electrode signals (using a Butterworth filter with a cutoff of 60 Hz) before performing the cross-correlation. To compute the electrode ILD, we measured the difference in root-mean-squared current (in µA) between the left and right ear electrode signals. These steps were performed for each electrode separately. The graphs in Fig. 6F and Supplementary Figs. 5-7 compare the ITD or ILD from electrode responses to those imposed on the noise band.

## Acknowledgments

The authors thank Charles Hem for assistance setting up the experiments with cochlear implant users, Alex Kell and Erica Shook for collecting the noise-vocoded speech data in normal hearing participants, Bertrand Delgutte for insightful discussions and Advanced Bionics for providing the software and hardware to collect data from cochlear implant users. Work supported by the K. Lisa Yang Brain-Body Center, a John W. Jarve Seed Fund Award, and NIH grant R01 DC017970.

---

**Supplementary Figure 1.**
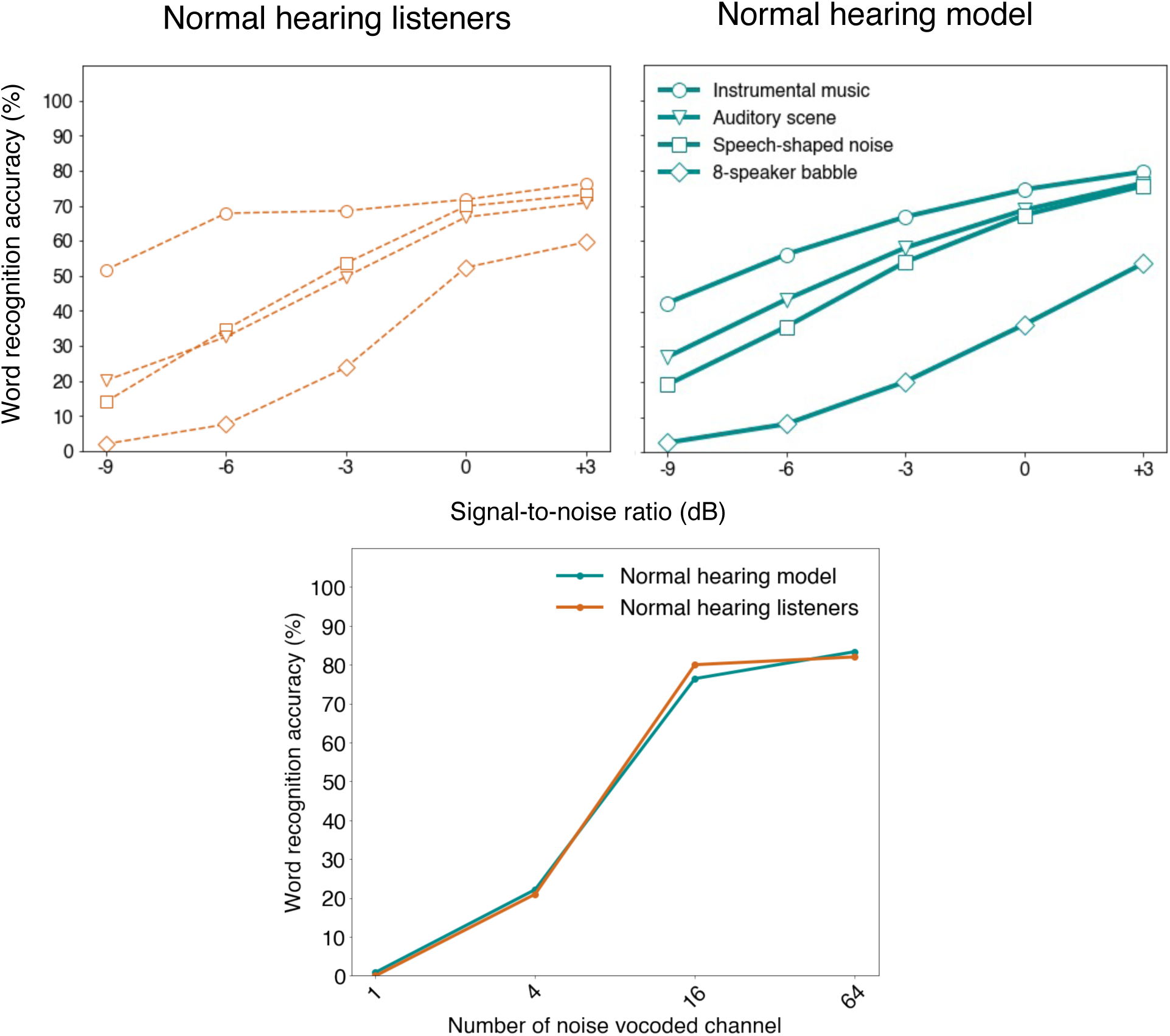
Comparison of word recognition by normal hearing model and normal hearing humans, for speech in noise and noise-vocoded speech.

**Supplementary Figure 2.**
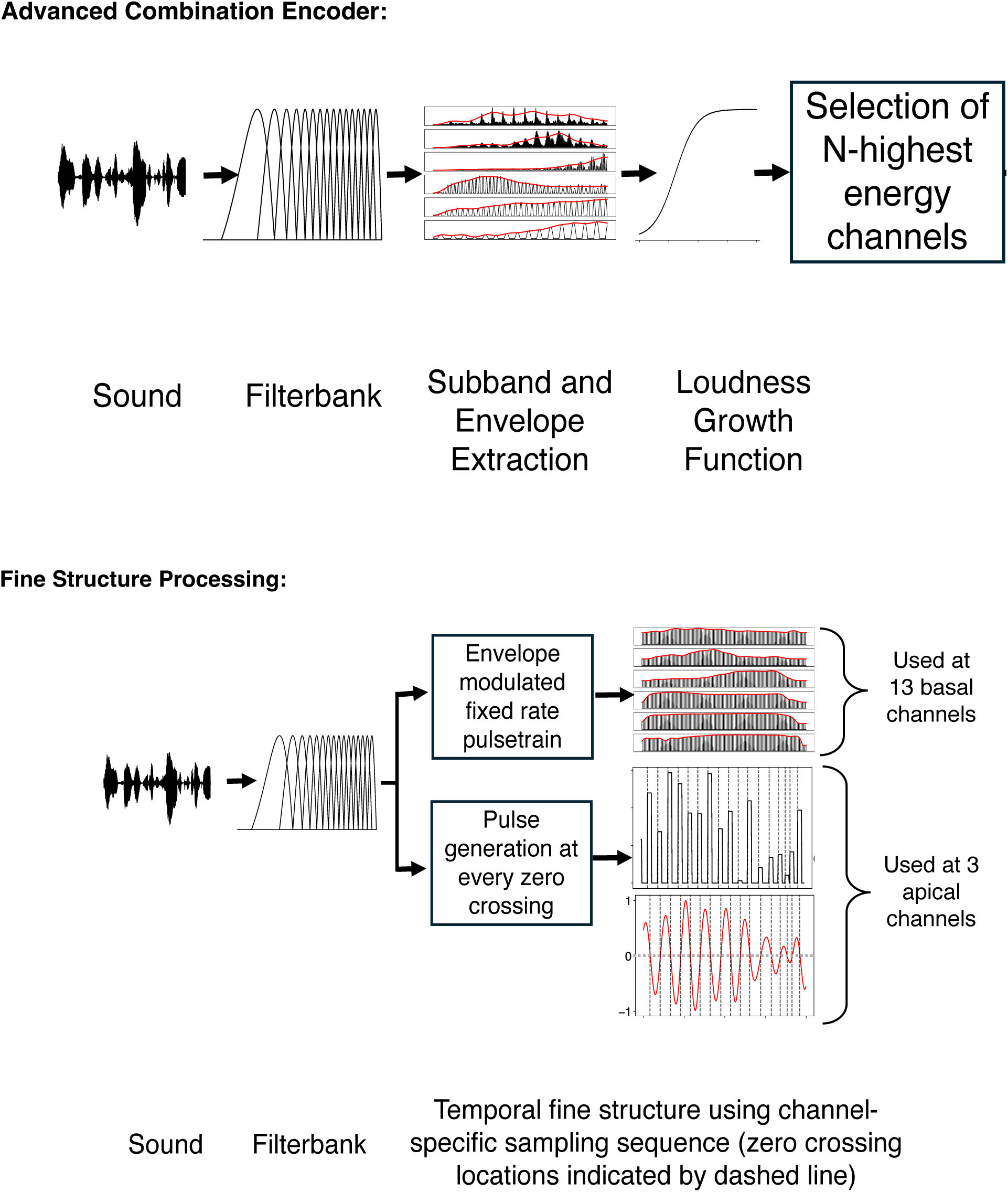
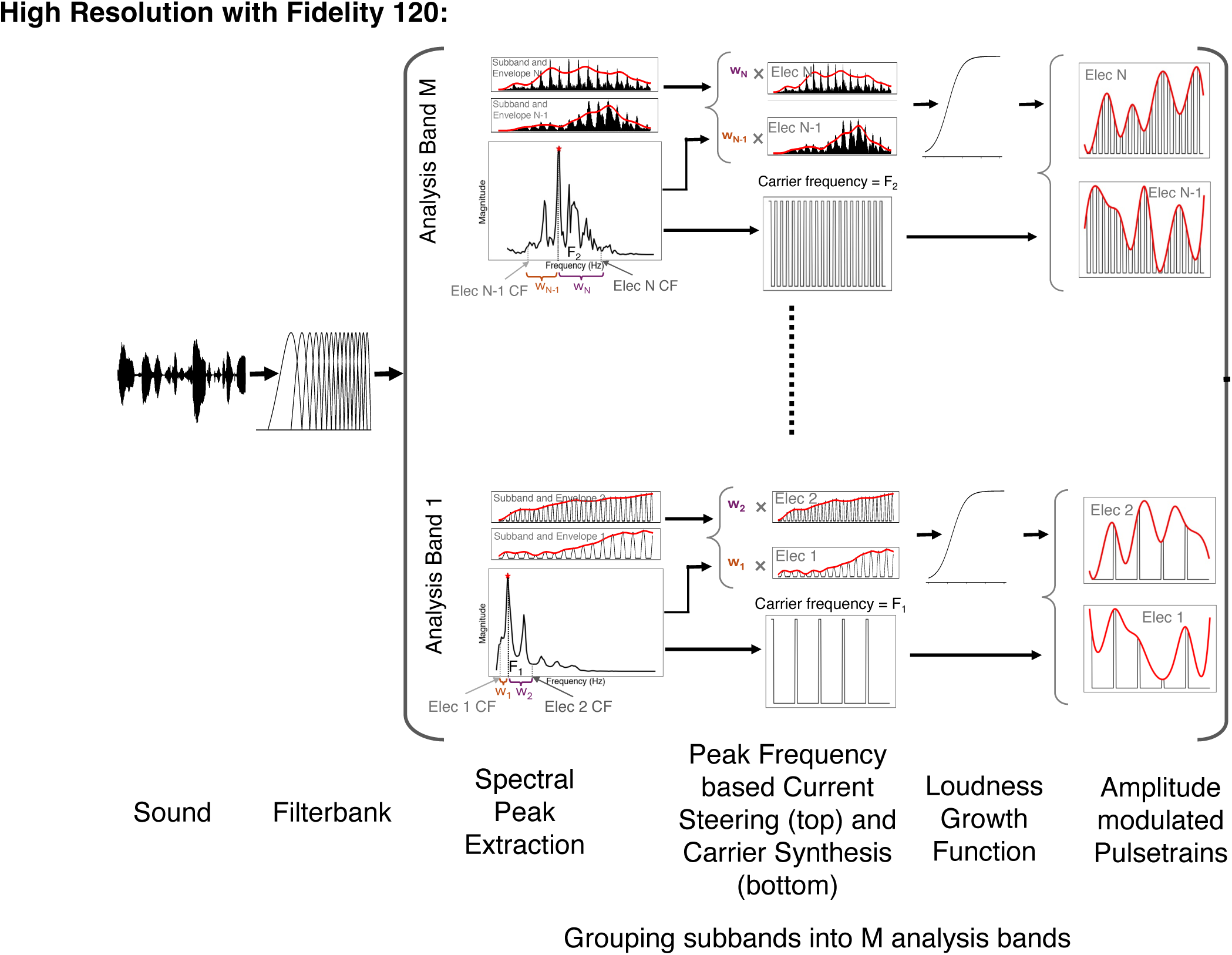
Schematic of different sound coding strategies

**Supplementary Figure 3.**
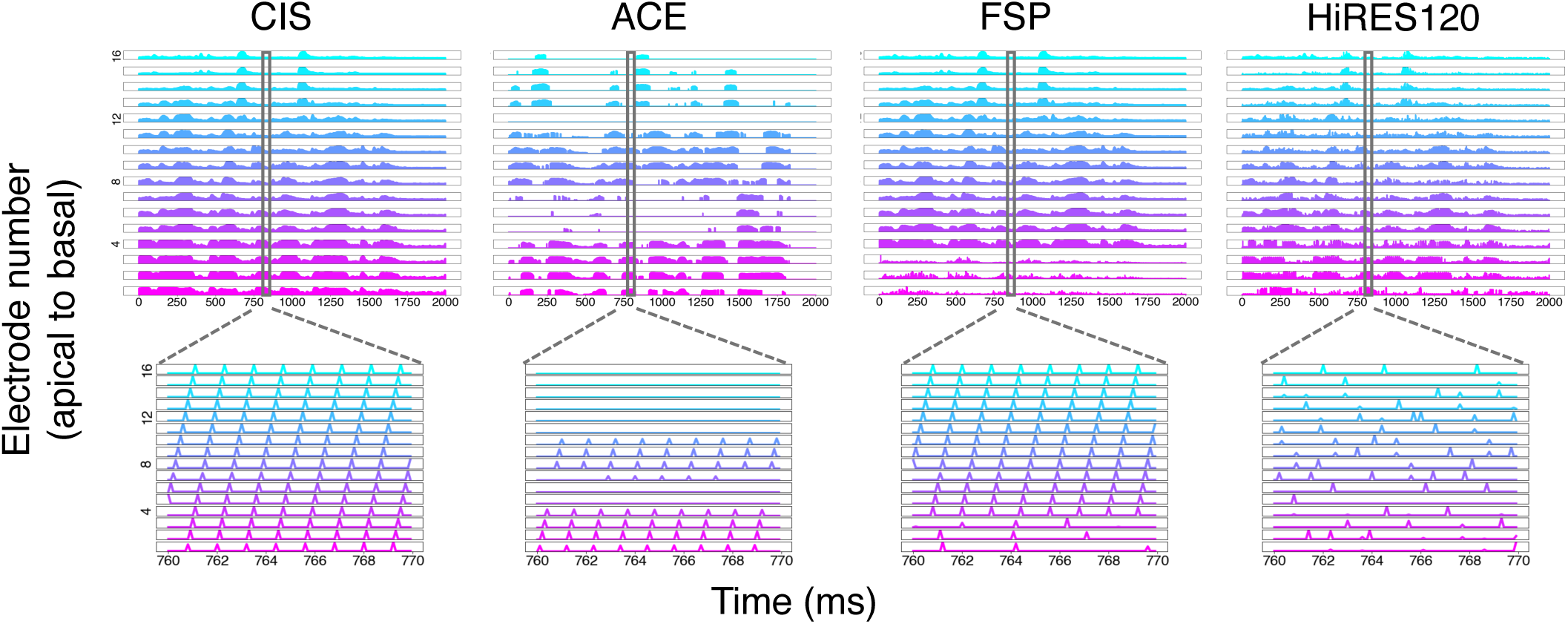
Example electrodograms from different sound coding strategies

**Supplementary Figure 4.**
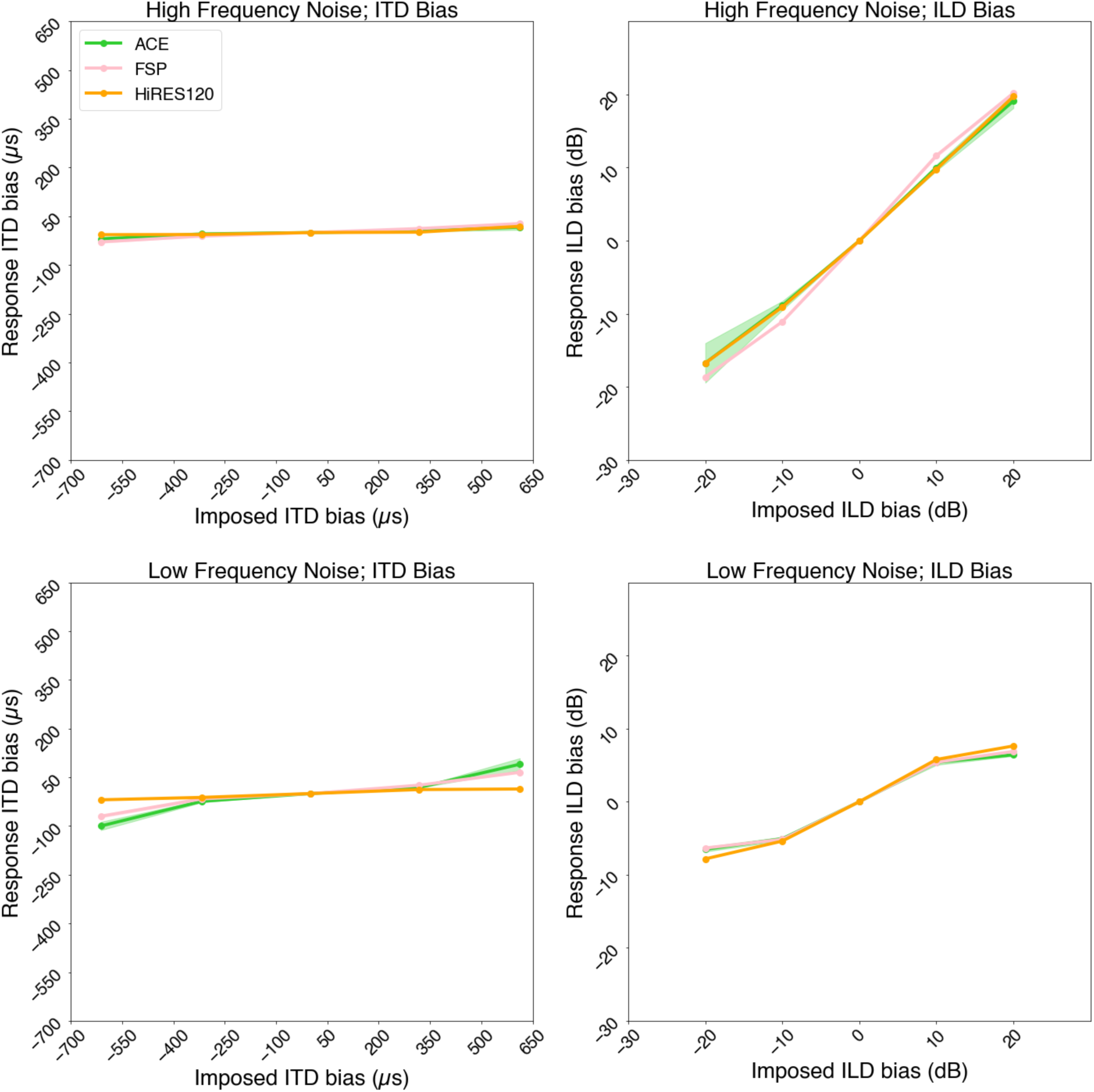
ITD-ILD bias for CI models with different sound coding strategies

**Supplementary Figure 5.**
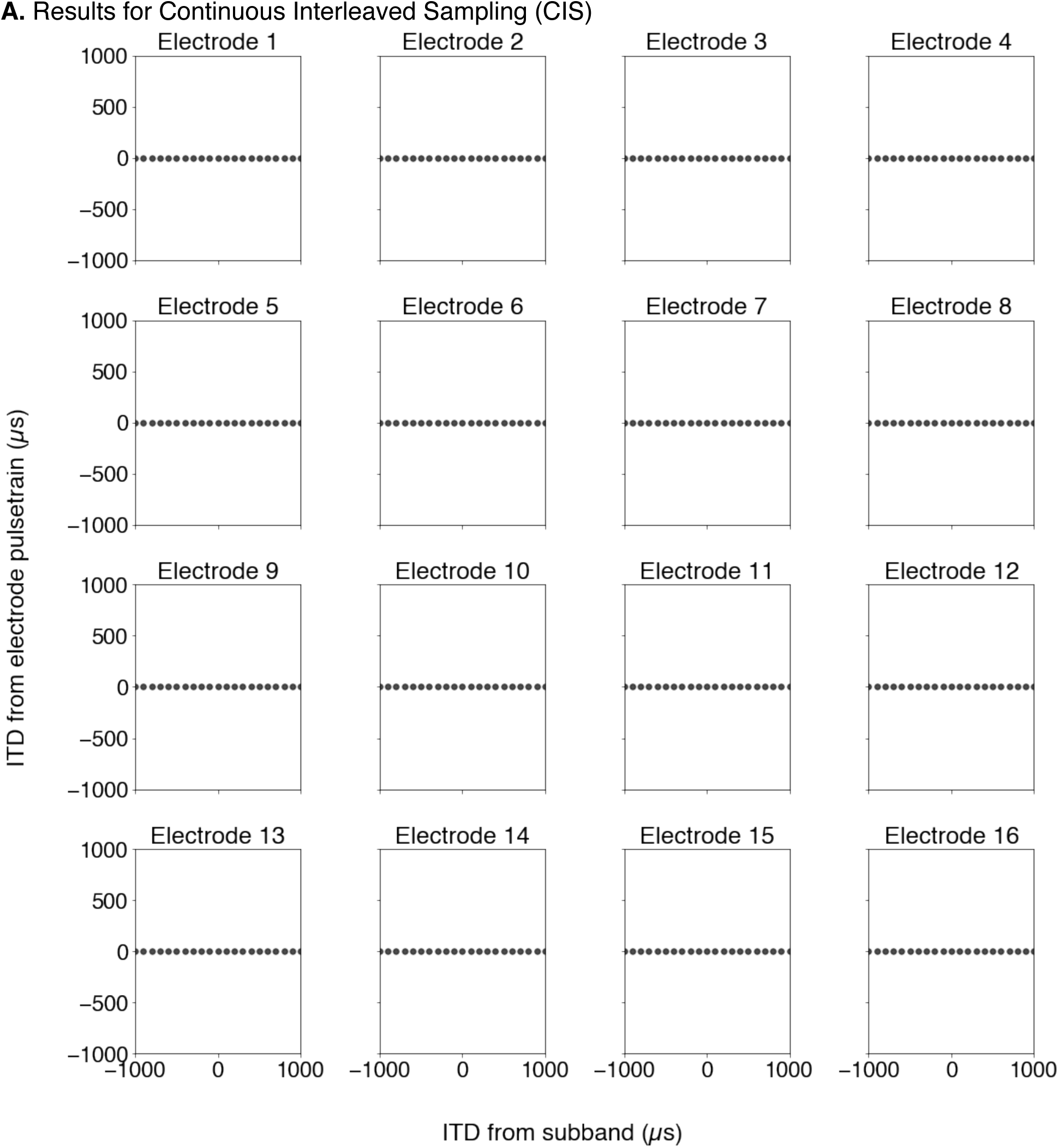
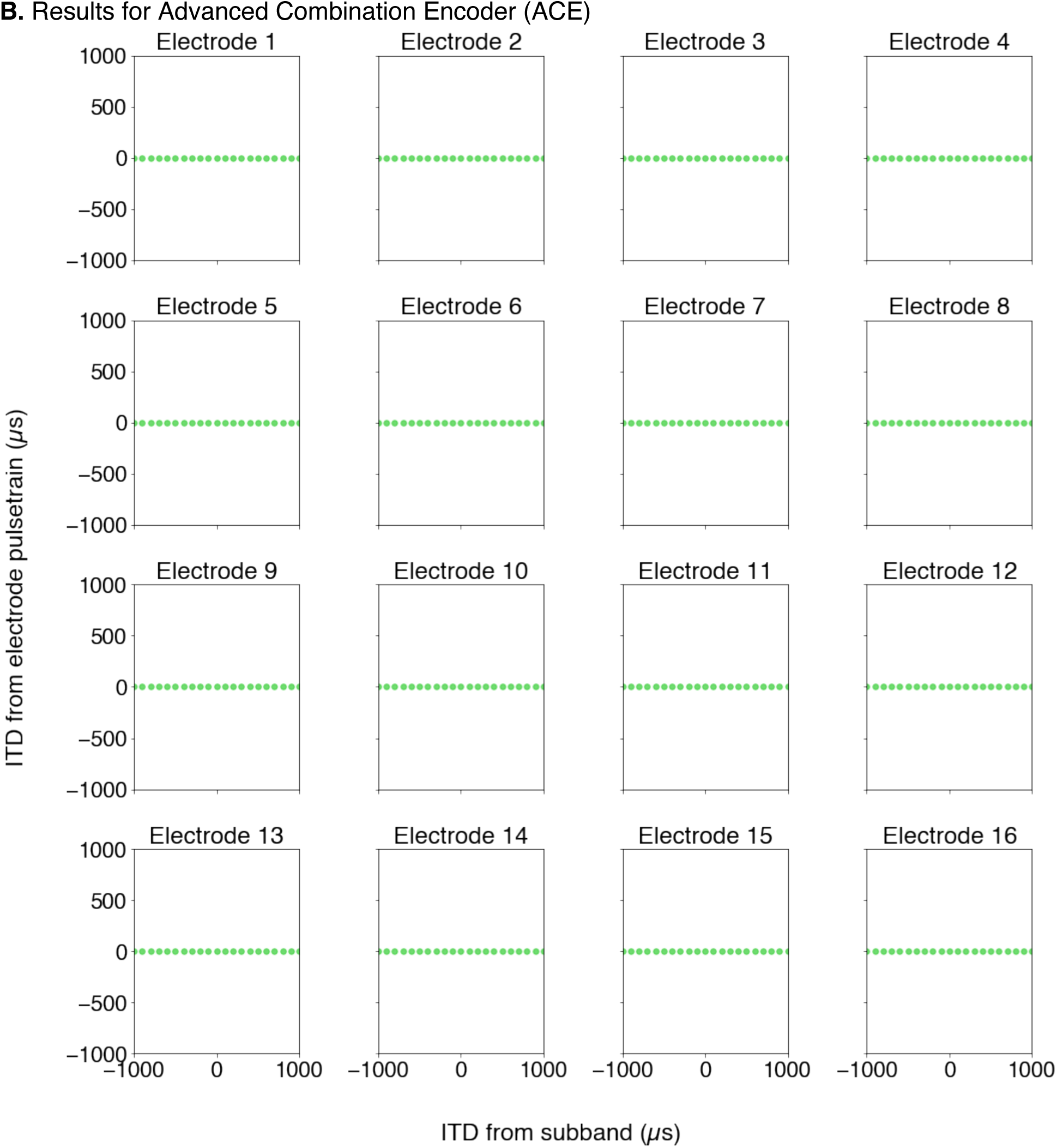
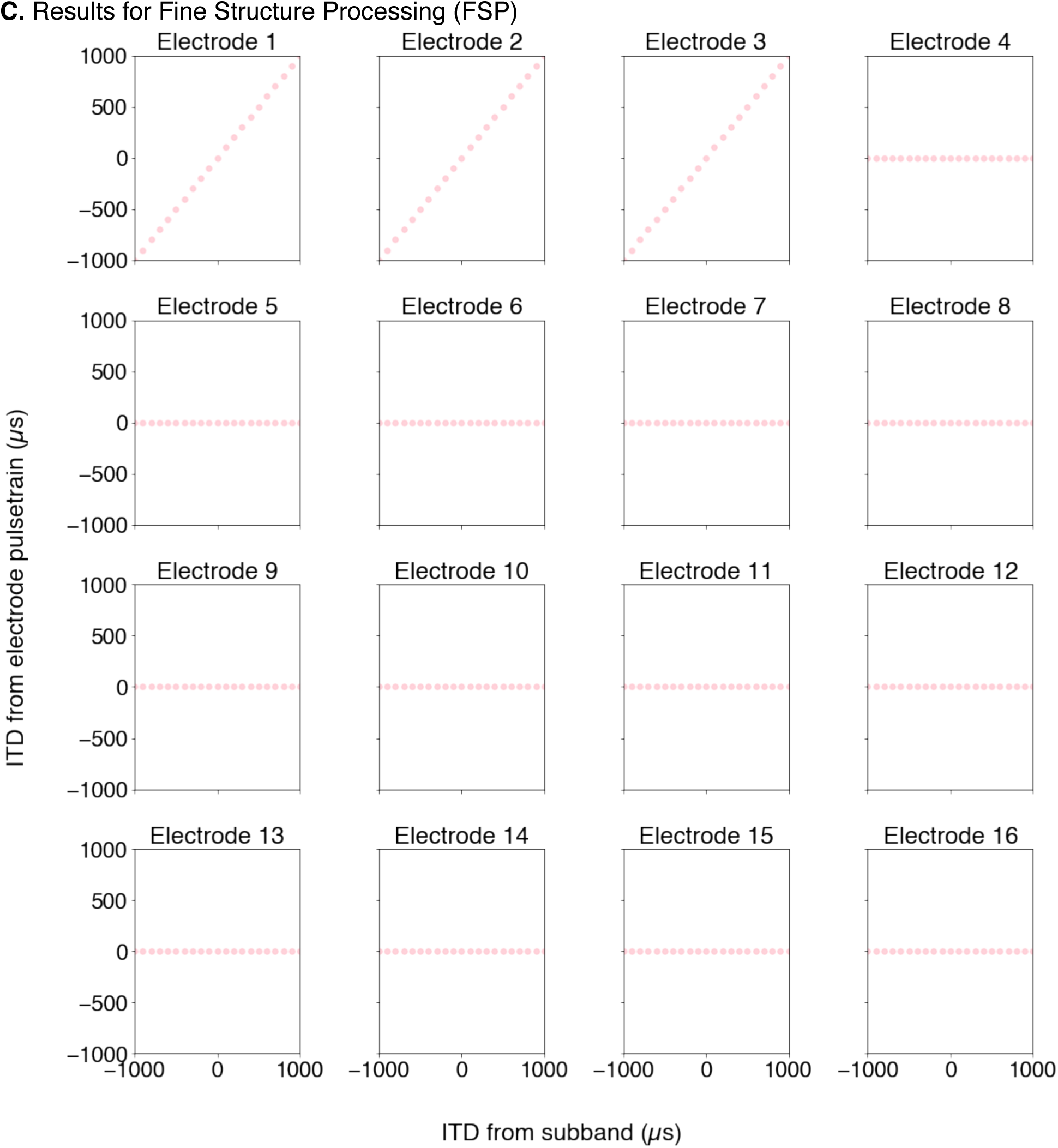
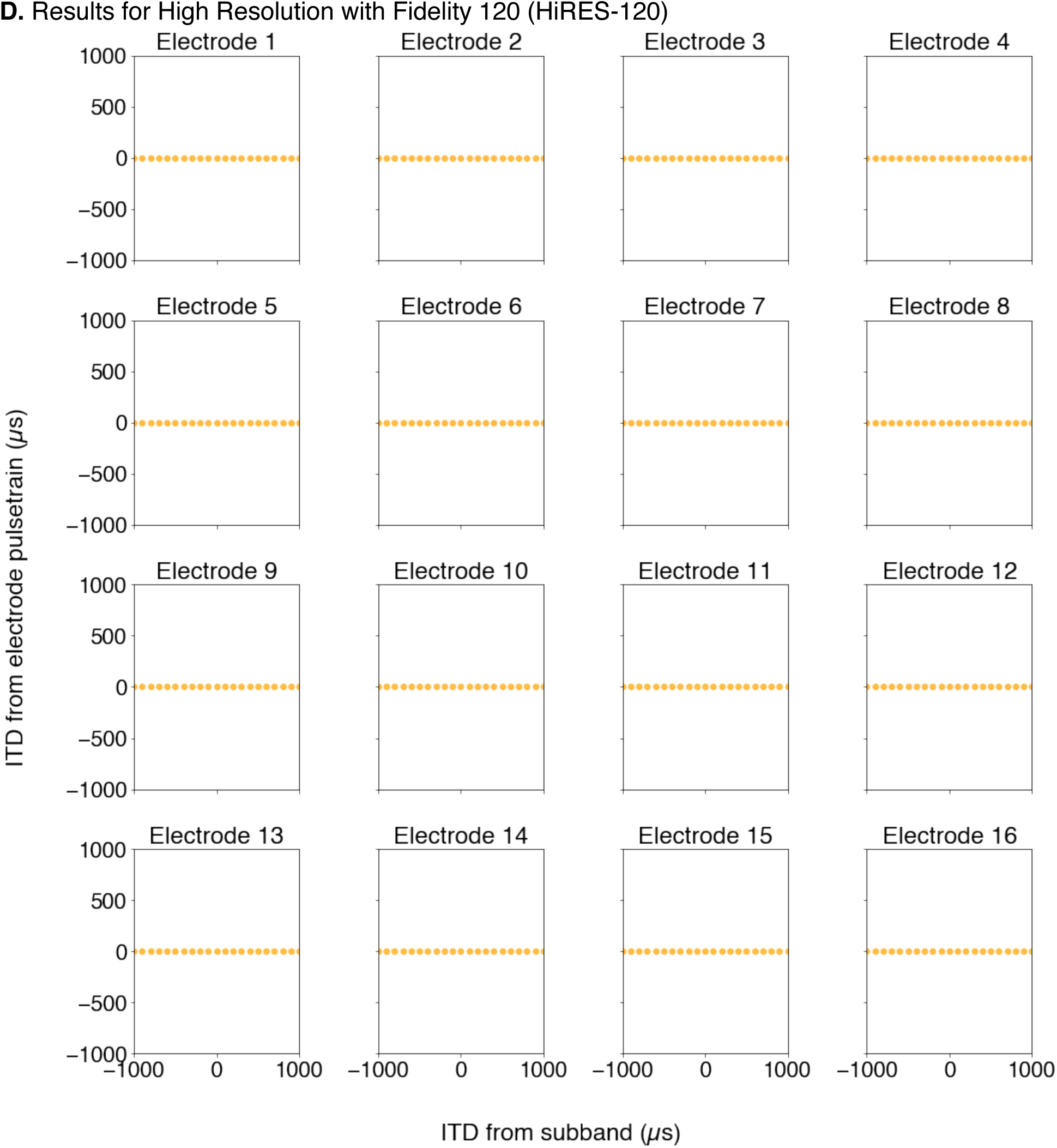
Analysis of fine structure ITDs obtained from electrode signals **A.** Results for Continuous Interleaved Sampling (CIS)

**Supplementary Figure 6.**
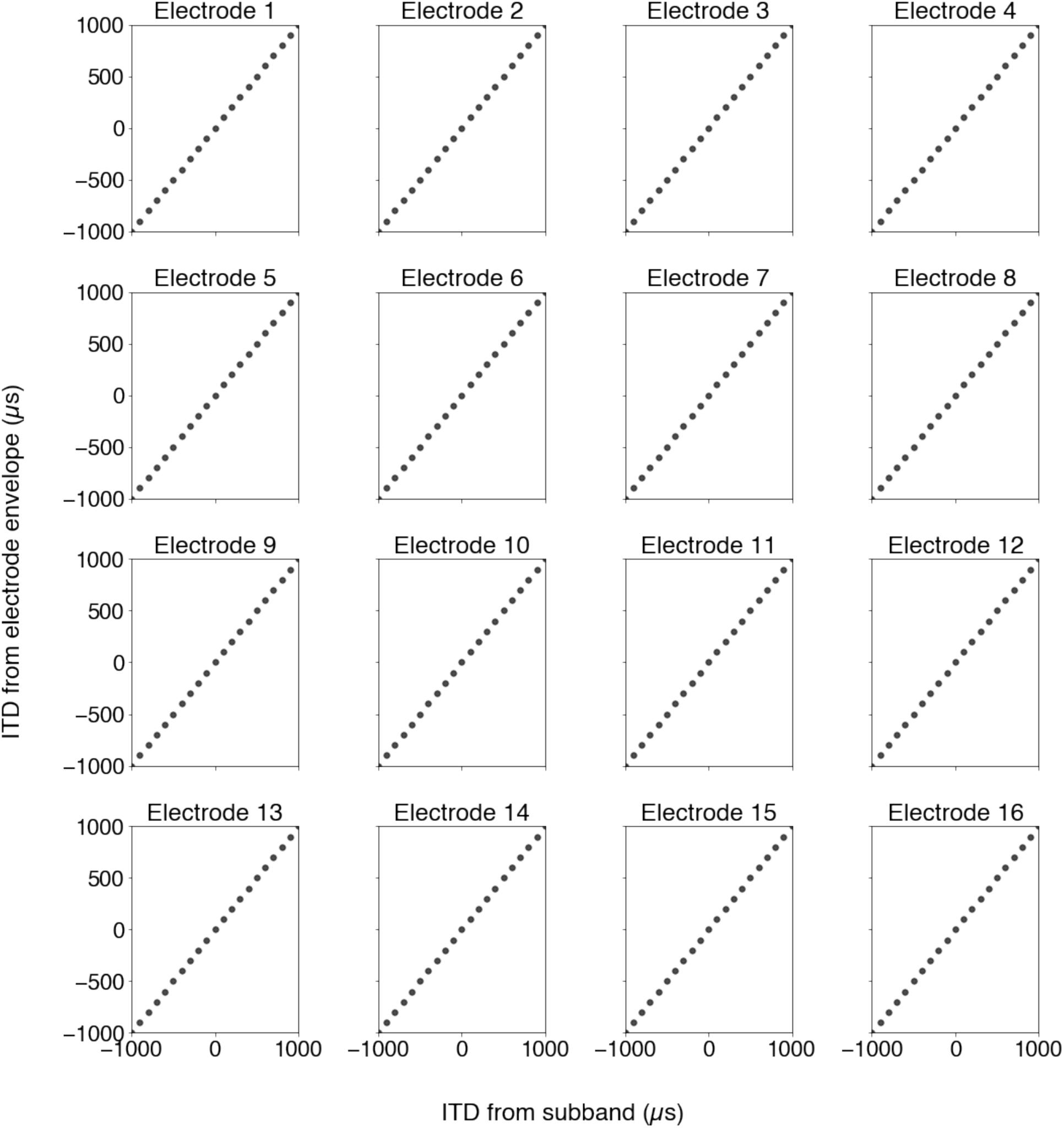
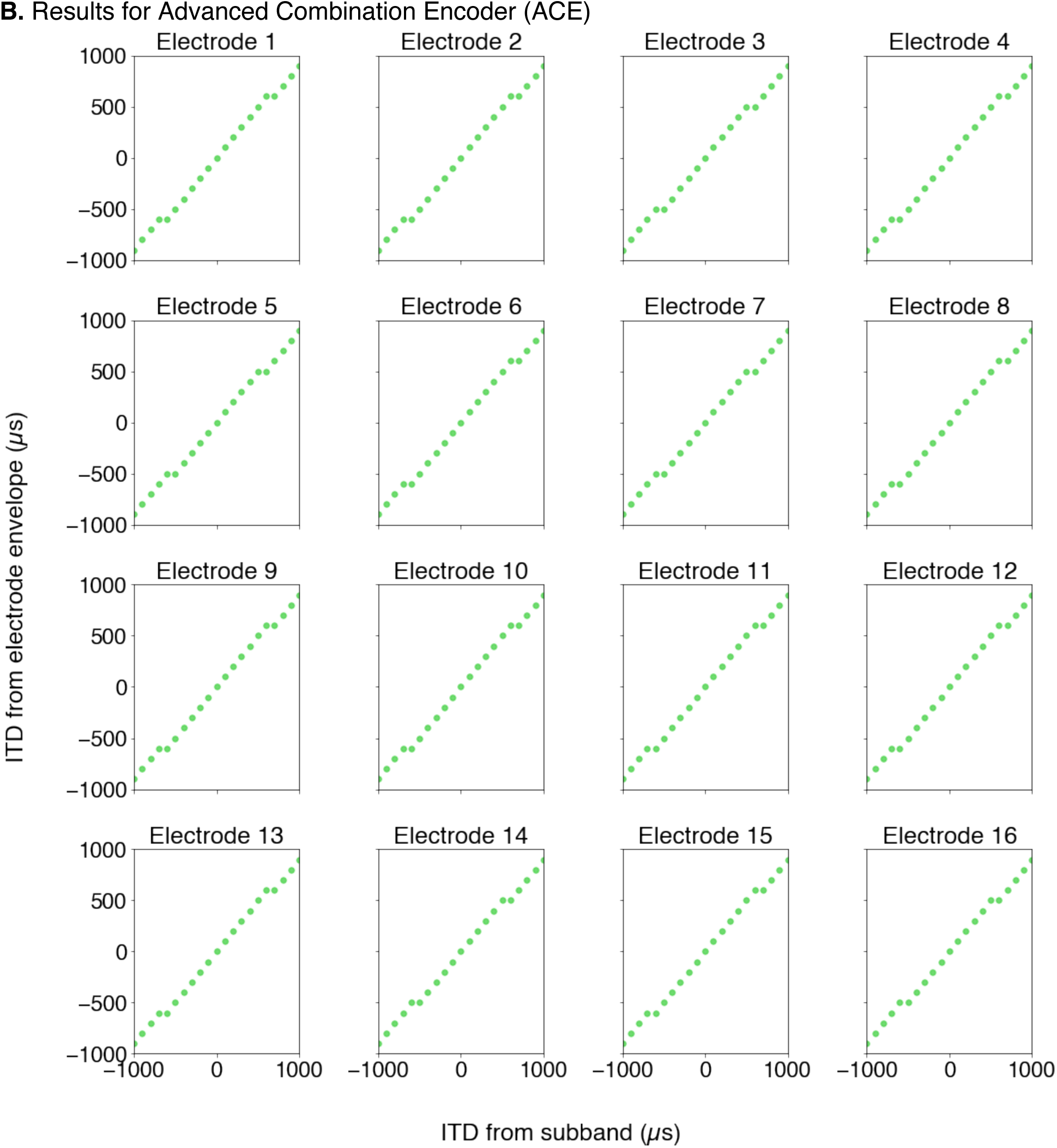
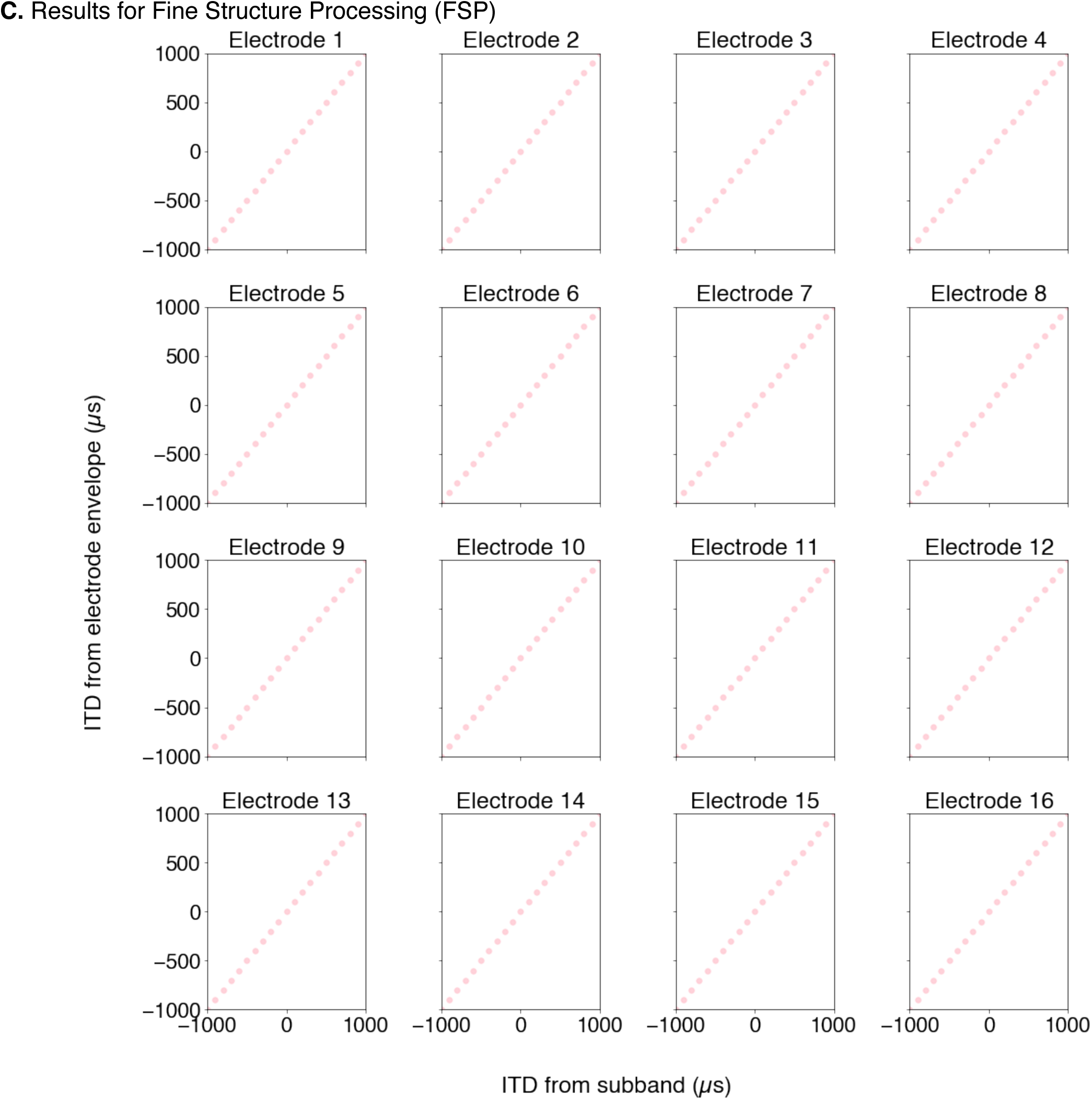
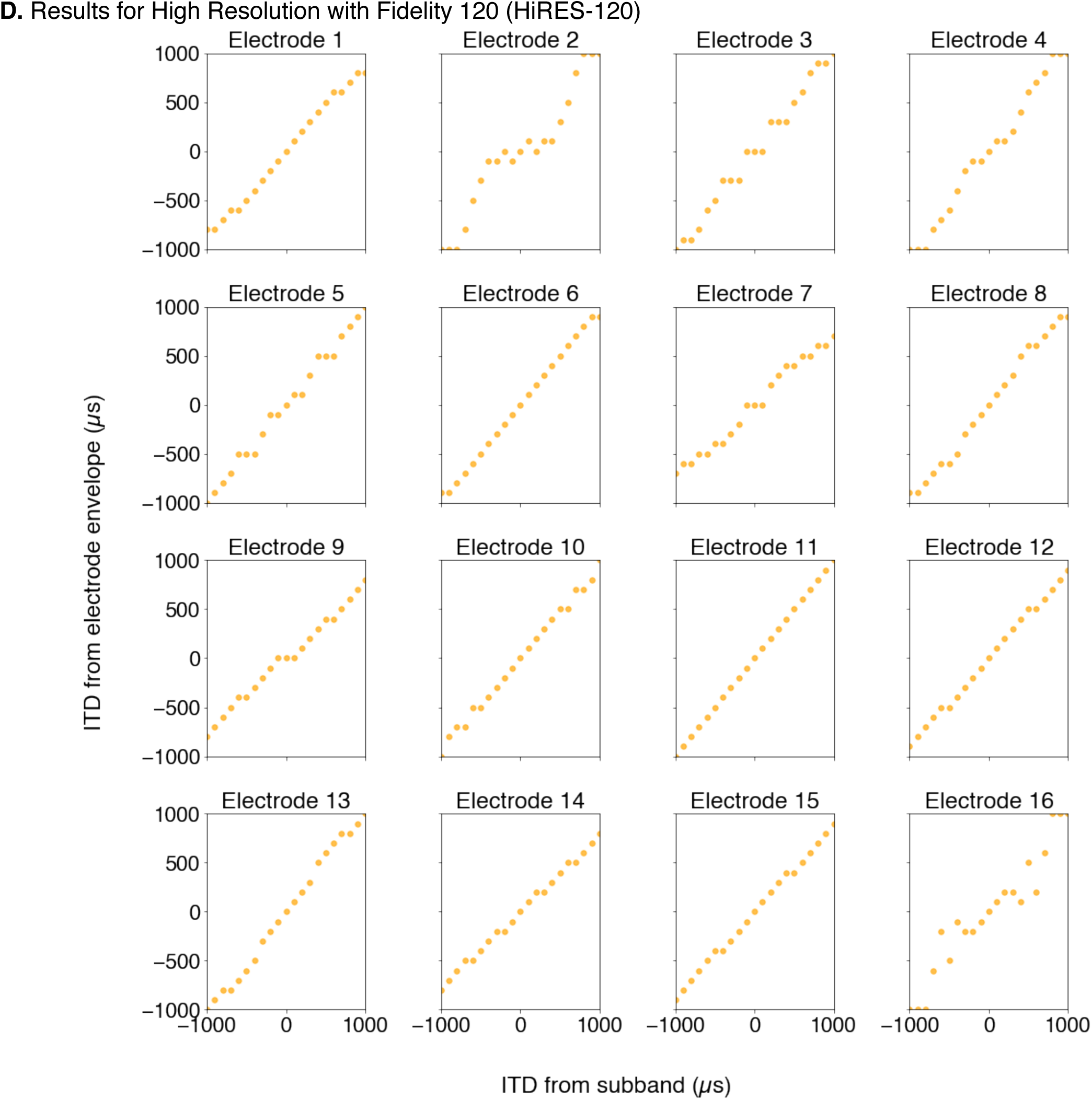
Analysis of envelope ITDs obtained from electrode signals

**Supplementary Figure 7.**
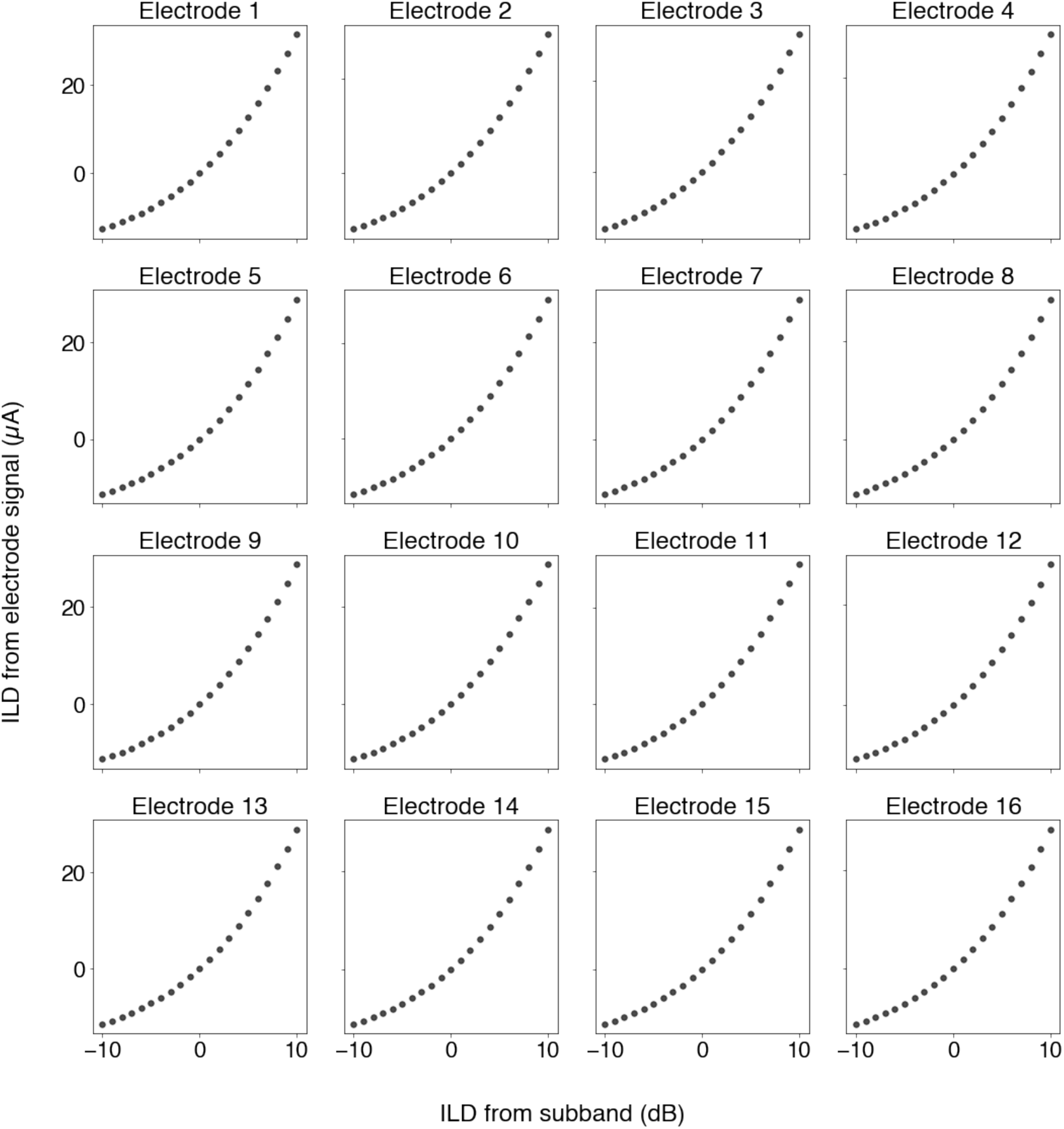
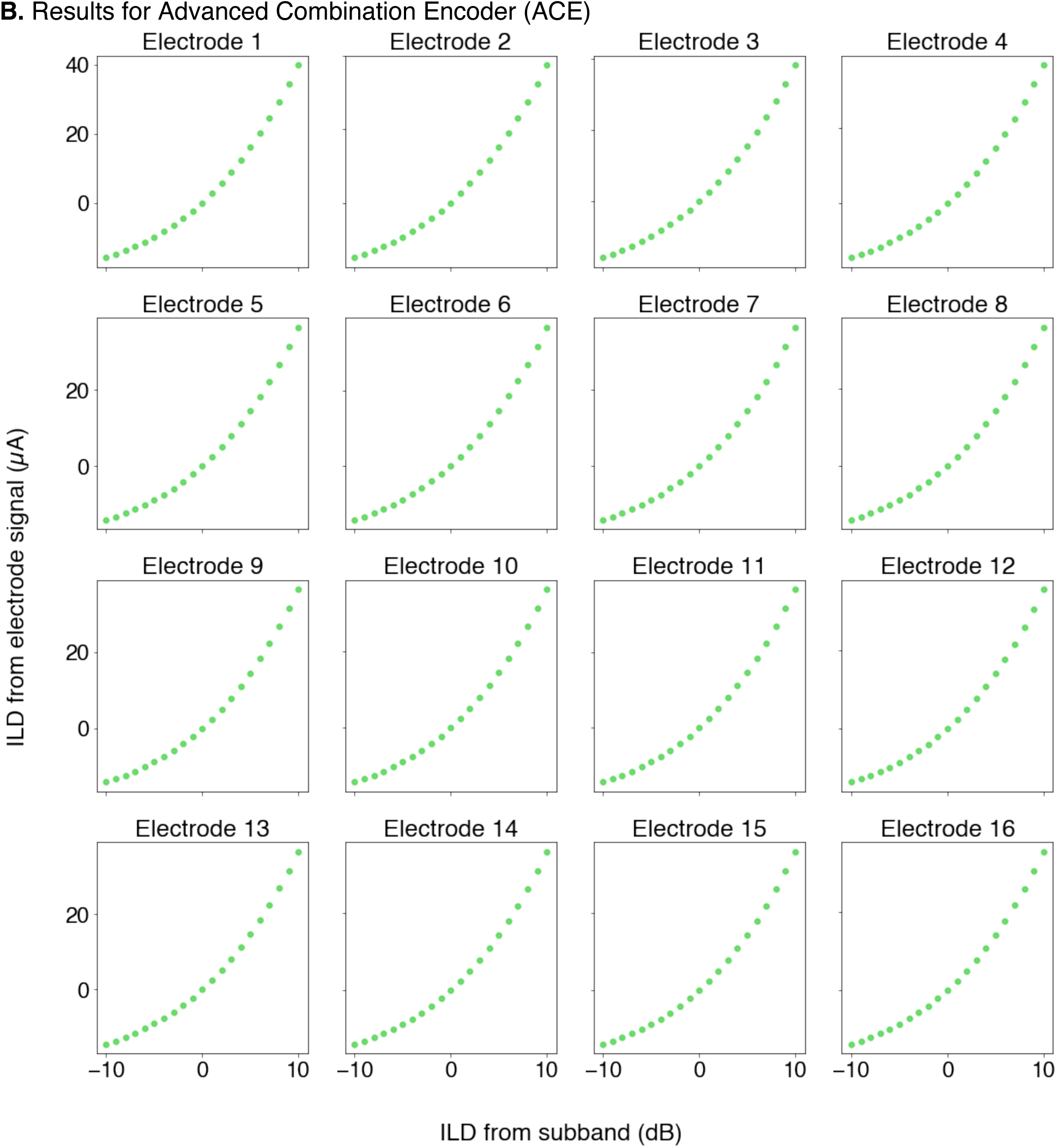
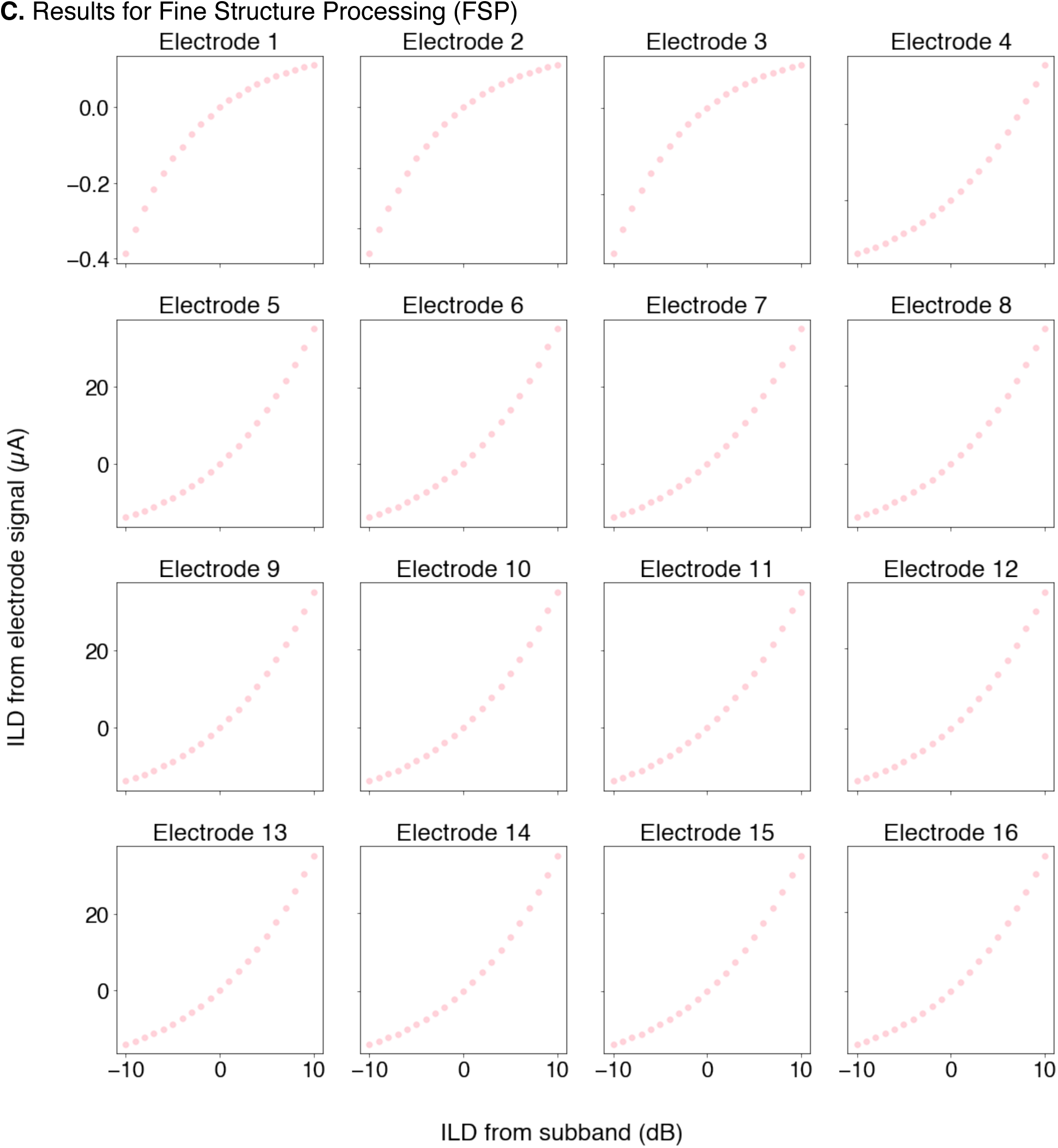
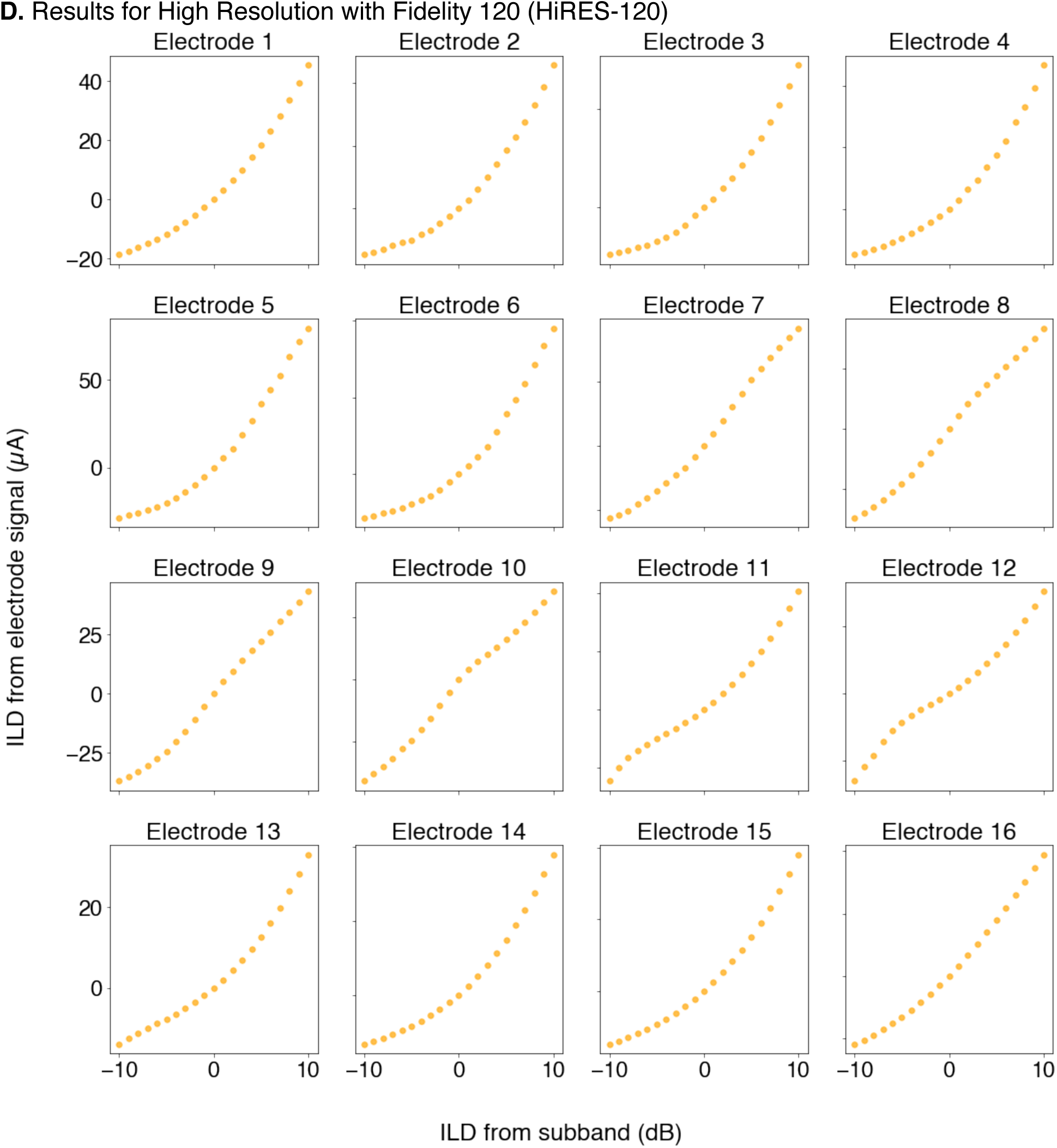
Analysis of ILDs obtained from electrode signals

**Supplementary Table 1.** Details of the three neural network architectures used for the word recognition task

| **arch0_0000** | **arch0_0001** | **arch0_0002** |
| --- | --- | --- |
| input_lnorm | input_lnorm | input_lnorm |
| conv0 [2, 42, 32] | conv0 [1, 84, 32] | conv0 [4, 21, 32] |
| relu0 | relu0 | relu0 |
| hpool0 [2, 4] | hpool0 [2, 4] | hpool0 [2, 4] |
| lnorm0 | lnorm0 | lnorm0 |
| conv1 [2, 18, 64] | conv1 [2, 18, 64] | conv1 [2, 18, 64] |
| relu1 | relu1 | relu1 |
| hpool1 [2, 4] | hpool1 [2, 4] | hpool1 [2, 4] |
| lnorm1 | lnorm1 | lnorm1 |
| conv2 [6, 6, 128] | conv2 [6, 6, 128] | conv2 [6, 6, 128] |
| relu2 | relu2 | relu2 |
| hpool2 [1, 4] | hpool2 [1, 4] | hpool2 [1, 4] |
| lnorm2 | lnorm2 | lnorm2 |
| conv3 [6, 6, 256] | conv3 [6, 6, 256] | conv3 [6, 6, 256] |
| relu3 | relu3 | relu3 |
| hpool3 [1, 4] | hpool3 [1, 4] | hpool3 [1, 4] |
| lnorm3 | lnorm3 | lnorm3 |
| conv4 [8, 8, 512] | conv4 [8, 8, 512] | conv4 [8, 8, 512] |
| relu4 | relu4 | relu4 |
| hpool4 [1, 1] | hpool4 [1, 1] | hpool4 [1, 1] |
| lnorm4 | lnorm4 | lnorm4 |
| conv5 [6, 6, 512] | conv5 [6, 6, 512] | conv5 [6, 6, 512] |
| relu5 | relu5 | relu5 |
| hpool5 [1, 1] | hpool5 [1, 1] | hpool5 [1, 1] |
| lnorm5 | lnorm5 | lnorm5 |
| conv6 [8, 8, 512] | conv6 [8, 8, 512] | conv6 [8, 8, 512] |
| relu6 | relu6 | relu6 |
| hpool6 [2, 4] | hpool6 [2, 4] | hpool6 [2, 4] |
| lnorm6 | lnorm6 | lnorm6 |
| flatten | flatten | flatten |
| fc0 [512] | fc0 [512] | fc0 [512] |
| relu_fc0 | relu_fc0 | relu_fc0 |
| norm_fc0 | norm_fc0 | norm_fc0 |
| dropout | dropout | dropout |
| fc [517, 433, 794] | fc [517, 433, 794] | fc [517, 433, 794] |

**Supplementary Table 2.**
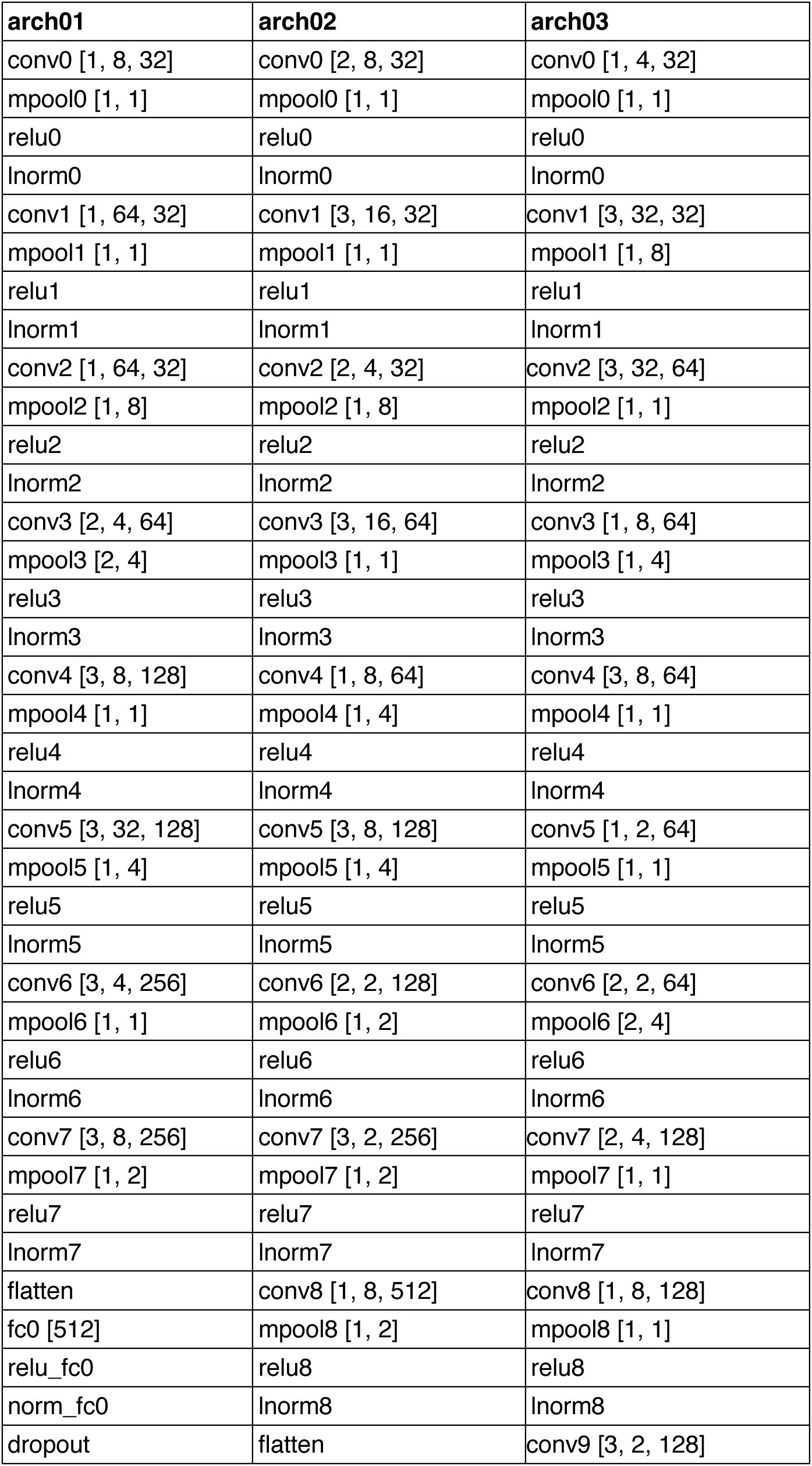
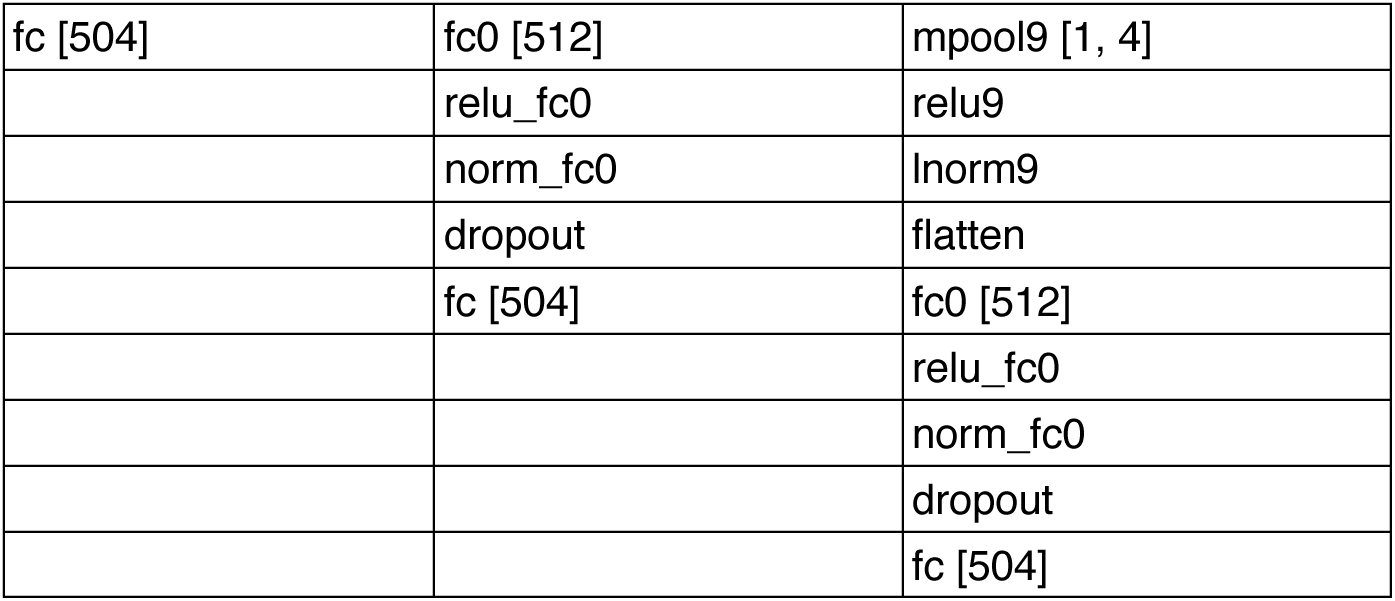
Details of the three neural network architectures used for the sound localization task

| arch01 | arch02 | arch03 |
| --- | --- | --- |
| conv0 [1, 8, 32] | conv0 [2, 8, 32] | conv0 [1, 4, 32] |
| mpool0 [1, 1] | mpool0 [1, 1] | mpool0 [1, 1] |
| relu0 | relu0 | relu0 |
| lnorm0 | lnorm0 | lnorm0 |
| conv1 [1, 64, 32] | conv1 [3, 16, 32] | conv1 [3, 32, 32] |
| mpool1 [1, 1] | mpool1 [1, 1] | mpool1 [1, 8] |
| relu1 | relu1 | relu1 |
| lnorm1 | lnorm1 | lnorm1 |
| conv2 [1, 64, 32] | conv2 [2, 4, 32] | conv2 [3, 32, 64] |
| mpool2 [1, 8] | mpool2 [1, 8] | mpool2 [1, 1] |
| relu2 | relu2 | relu2 |
| lnorm2 | lnorm2 | lnorm2 |
| conv3 [2, 4, 64] | conv3 [3, 16, 64] | conv3 [1, 8, 64] |
| mpool3 [2, 4] | mpool3 [1, 1] | mpool3 [1, 4] |
| relu3 | relu3 | relu3 |
| lnorm3 | lnorm3 | lnorm3 |
| conv4 [3, 8, 128] | conv4 [1, 8, 64] | conv4 [3, 8, 64] |
| mpool4 [1, 1] | mpool4 [1, 4] | mpool4 [1, 1] |
| relu4 | relu4 | relu4 |
| lnorm4 | lnorm4 | lnorm4 |
| conv5 [3, 32, 128] | conv5 [3, 8, 128] | conv5 [1, 2, 64] |
| mpool5 [1, 4] | mpool5 [1, 4] | mpool5 [1, 1] |
| relu5 | relu5 | relu5 |
| lnorm5 | lnorm5 | lnorm5 |
| conv6 [3, 4, 256] | conv6 [2, 2, 128] | conv6 [2, 2, 64] |
| mpool6 [1, 1] | mpool6 [1, 2] | mpool6 [2, 4] |
| relu6 | relu6 | relu6 |
| lnorm6 | lnorm6 | lnorm6 |
| conv7 [3, 8, 256] | conv7 [3, 2, 256] | conv7 [2, 4, 128] |
| mpool7 [1, 2] | mpool7 [1, 2] | mpool7 [1, 1] |
| relu7 | relu7 | relu7 |
| lnorm7 | lnorm7 | lnorm7 |
| flatten | conv8 [1, 8, 512] | conv8 [1, 8, 128] |
| fc0 [512] | mpool8 [1, 2] | mpool8 [1, 1] |
| relu_fc0 | relu8 | relu8 |
| norm_fc0 | lnorm8 | lnorm8 |
| dropout | flatten | conv9 [3, 2, 128] |
| fc [504] | fc0 [512] | mpool9 [1, 4] |
|  | relu_fc0 | relu9 |
|  | norm_fc0 | lnorm9 |
|  | dropout | flatten |
|  | fc [504] | fc0 [512] |
|  |  | relu_fc0 |
|  |  | norm_fc0 |
|  |  | dropout |
|  |  | fc [504] |

## Notes

### Competing Interest Statement

The authors have declared no competing interest.

### Summary of Updates

The updated pre-print includes additional model results, analyzing the effect of spread of excitation expected from tripolar stimulation on word recognition and sound localization performance. It also includes additional analysis of the interaural time and level difference cues that are availabe from the cochlear implant electrodograms with different sound coding strategies.

